# HSV-1 infection induces a downstream shift of the +1 nucleosome

**DOI:** 10.1101/2024.03.06.583707

**Authors:** Elena Weiß, Adam W. Whisnant, Thomas Hennig, Lara Djakovic, Lars Dölken, Caroline C. Friedel

## Abstract

Herpes simplex virus 1 (HSV-1) infection induces a loss of host transcriptional activity and widespread disruption of host transcription termination, which leads to an induction of open chromatin downstream of genes. In this study, we show that lytic HSV-1 infection also leads to an extension of chromatin accessibility at promoters into downstream regions. This is most prominent for highly expressed genes and independent of the immediate-early proteins ICP0, ICP22, and ICP27 and the virion host shutoff protein *vhs.* ChIPmentation of the noncanonical histone variant H2A.Z, which is strongly enriched at +1 and −1 nucleosomes, indicated that these chromatin accessibility changes are linked to a downstream shift of +1 nucleosomes. In yeast, downstream shifts of +1 nucleosomes are induced by RNA Polymerase II (Pol II) degradation. Accordingly, irreversible depletion of Pol II from genes in human cells using α-amanitin altered +1 nucleosome positioning similar to lytic HSV-1 infection. Consequently, treatment with phosphonoacetic acid (PAA) and knockout of ICP4, which both prevent viral DNA replication and alleviate the loss of Pol II from host genes, largely abolished the downstream extension of accessible chromatin in HSV-1 infection. In the absence of viral DNA replication, doxycycline-induced expression of ICP27, which redirects Pol II from gene bodies into intergenic regions by disrupting transcription termination, induced an attenuated effect that was further enhanced by co-expression of ICP22. In summary, our study provides strong evidence that HSV-1-induced depletion of Pol II from the host genome leads to a downstream shift of +1 nucleosomes at host promoters.

**Importance:** Lytic herpes simplex virus 1 (HSV-1) infection leads to a profound host transcription shutoff. Loss of RNA Polymerase II (Pol II) in yeast has previously been shown to relax +1 nucleosome positioning to more thermodynamically favorable sites downstream of transcription start sites. Here, we show that a similar phenomenon is likely at play in lytic HSV-1 infection. Sequencing of accessible chromatin revealed a widening of nucleosome-free regions at host promoters into downstream regions. By mapping genome-wide positions of the noncanonical histone variant H2A.Z enriched at +1 and −1 nucleosomes, we demonstrate a downstream shift of +1 nucleosomes for most cellular genes in lytic HSV-1 infection. As chemical depletion of Pol II from genes also leads to a downstream shift of +1 nucleosomes in human cells, changes in chromatin architecture at promoters in HSV-1 infection are likely a consequence of HSV-1-induced loss of Pol II activity from the host genome.

## Introduction

Herpes simplex virus 1 (HSV-1) is one of nine human herpesviruses, and more than half of the global population are latently infected with HSV-1 (1–4). HSV-1 does not only cause the common cold sores but is also responsible for life-threatening diseases, particularly in the immunocompromised (5). The HSV-1 genome consists of 152 kilobases (kb) of double-stranded DNA and encodes for at least 121 large open reading frames (ORFs) and >100 small ORFs (6). Upon lytic infection, four of the five viral immediate early (IE) genes, i.e., ICP0, ICP4, ICP22 and ICP27, hijack the host gene expression machinery to ensure viral replication during early stages of infection (7–11). This results in the recruitment of RNA polymerase II (Pol II) from the host chromatin to viral genomes and a general shutoff of host transcription (7, 9, 11–13). Pol II recruitment to viral genomes is predominantly facilitated by the viral IE protein ICP4 (13), which effectively sequesters Pol II into the viral replication compartments (14). The global host transcription shutoff is further exacerbated by ICP22-mediated inhibition of host transcription elongation (15) and the viral *vhs* (virion host shutoff) protein. *vhs* is a nuclease delivered by the tegument of the incoming viral particles that cleaves both viral and host mRNAs (16–18). We recently demonstrated that *vhs* continuously degrades about 30 % of host mRNAs per hour during the first 8 h of high multiplicity lytic infection of primary human fibroblasts (19).

HSV-1 further disturbs host transcription by inducing a widespread disruption of host transcription termination (DoTT) resulting in read-through transcription for tens of thousands of nucleotides (nt) beyond cellular polyadenylation (pA) sites (20). Read-through transcription commonly extends into downstream genes, leading to a seeming transcriptional induction of these genes. DoTT negatively affects host gene expression in two ways: First, as read-through transcripts are not exported (21), DoTT prevents the translation of newly transcribed mRNAs and thus dampens the host transcriptional response to infection. Second, it redirects a large fraction of ongoing Pol II transcription from host gene bodies into downstream intergenic regions and, thus, further exacerbates the depletion of Pol II from host genes. We recently showed that ICP27 is sufficient for inducing DoTT (22). Interestingly, transcription downstream of genes (DoG) is also induced in cellular stress responses (21, 23–25). Accordingly, infection with an ICP27-null mutant still induces read-through transcription, albeit at much lower levels, presumably due to a virus-induced cellular stress response (26). In contrast to stress-induced DoGs, HSV-1-induced DoTT is associated with a massive increase of chromatin accessibility downstream of genes (denoted as downstream open chromatin regions (dOCRs)) detectable by ATAC-seq (Assay for Transposase-Accessible Chromatin using sequencing) (21). dOCR induction requires strong absolute levels of transcription downstream of genes, thus it is predominantly observed for highly transcribed genes with strong read-through. Treatment with phosphonoacetic acid (PAA) during HSV-1 infection, which inhibits viral DNA replication (27, 28), results in increased dOCR induction (29). As viral DNA redirects Pol II from the host chromatin, PAA treatment mitigates the loss of Pol II from the host genome and thus increases absolute levels of transcription on host genes and – importantly for dOCR induction – read-through regions. We recently showed that ICP22 is both required and sufficient for inducing dOCRs in the presence of ICP27-induced read-through (29).

As we previously only investigated chromatin accessibility *downstream* of genes, we now further explore our previously published ATAC-seq data to examine changes in chromatin accessibility around host gene promoters. This revealed a broadening of open chromatin regions at promoters into regions downstream of the transcription start sites (TSS) for most host genes. While it was consistently observed in the absence of ICP0, ICP22, ICP27, and *vhs*, PAA treatment and ICP4 knockout, both of which inhibit viral DNA replication and alleviate the HSV-1-induced loss of host transcription, substantially reduced the extension of open chromatin at the TSS. Weiner *et al.* previously showed that Pol II depletion in yeast induces downstream shifts of +1 nucleosomes, in particular for highly expressed genes (30). Transcribing Pol II disturbs histone-DNA interactions and, at high transcription rates, leads to eviction of nucleosomes from the chromatin (31). Weiner *et al.* proposed that Pol II depletion relaxes chromatin, allowing nucleosomes to shift to more thermodynamically favorable sites. The noncanonical histone H2A variant H2A.Z is strongly enriched at gene promoters at +1 and −1 nucleosome positions (32, 33), likely as H2A.Z deposition increases nucleosome mobility and makes DNA more accessible to the transcriptional machinery (34). We thus performed ChIPmentation for H2A.Z during lytic HSV-1 infection to map +1 nucleosome positions. This showed that changes in chromatin accessibility in HSV-1 infection reflected a downstream shift of +1 nucleosomes. We observed similar changes in H2A.Z occupancy upon treatment with α-amanitin, a deadly toxin found in Amanita mushrooms, which inhibits transcription by preventing translocation of Pol II and triggering degradation of Rpb1, the largest subunit of Pol II (35). Our findings thus provide strong evidence that depletion of Pol II from host genes during HSV-1 infection leads to the downstream shift of +1 nucleosomes to more thermodynamically favorable sites.

## Results

### Widespread extension of chromatin-free regions downstream of transcription start sites

To investigate changes in chromatin accessibility within promoter regions during HSV-1 infection, we re-analyzed our recently published ATAC-seq data (29) for mock and wild-type (WT) HSV-1 infection of human fetal foreskin fibroblasts (HFF) as well as infection with null mutant viruses for the HSV-1 IE proteins ICP0, ICP22, and ICP27 as well as the tegument-delivered late protein *vhs*. Samples for mock, WT, Δ*vhs,* and ΔICP27 infection were collected at 8 h p.i., while samples for ΔICP0 and ΔICP22 infection were collected at 12 h p.i. To assess chromatin accessibility changes in promoter regions, we identified genomic regions with differential ATAC-seq read density using our recently published RegCFinder method (36). RegCFinder searches for genomic subregions for which the read distribution differs between two conditions (e.g., mock and WT infection). It is targeted towards regions of interest by specifying genomic windows as input (**Sup. Fig 1a** in supplemental material). For each input window, subregions that show differences in the distribution of reads within this window are identified. Statistical significance and log2 fold-changes for identified subregions are determined by DEXSeq (37). We defined the windows of interest as the ± 3kb promoter region around the TSS of 7,649 human genes. These TSSs were previously defined in our analysis of promoter-proximal Pol II pausing in HSV-1 WT infection (38) based on a re-analysis of published precision nuclear run-on sequencing (PRO-seq) and PROcap-seq (a variation of PRO-seq) data of flavopiridol-treated uninfected HFF. Inhibition of the CDK9 subunit of the positive transcription elongation factor b (P-TEFb) by flavopiridol arrests Pol II in a paused state at the TSS, allowing the mapping of Pol II initiation sites with both PRO- and PROcap-seq. Filtering for high-confidence TSSs confirmed by both data sets and within a maximum distance of 500 bp to the nearest annotated gene on the same strand identified the major TSS for 7,649 genes. Using these 6 kb promoter windows as input, we applied RegCFinder to our ATAC-seq data for WT and all null-mutant infections in comparison to mock to identify promoter subregions that show differences in the distribution of open chromatin between infection and mock.

It should be noted that dOCRs can extend into downstream genes following very strong read-through transcription. We nevertheless did not filter promoter windows beforehand to exclude genes for which dOCRs extended from an upstream gene into their promoter window for two reasons: (i) Since dOCRs represent changes in chromatin accessibility, RegCFinder should, by design, detect them; (ii) The farther the distance from the upstream gene, the less pronounced are changes in chromatin accessibility in dOCR regions, making it difficult to distinguish them from background. Any approach we tested to filter promoter windows overlapping with dOCRs from upstream genes before the RegCFinder analysis either excluded too many or too few windows. Thus, we decided to first run RegCFinder and then investigate which detected changes were due to the induction of dOCRs from upstream genes.

RegCFinder identified 23,000-24,000 differential subregions (in the following also denoted as RegC, short for regions of change) for each virus infection compared to mock (WT: 23,357 RegC, ΔICP0: 23,532, ΔICP22: 24,640, ΔICP27: 23,523, Δ*vhs*: 23,716, **Sup. Fig 1b**). Between 29 and 37 % of these showed a statistically significant difference in read density within the corresponding 6 kb promoter windows upon infection (multiple testing adjusted p-value (adj. p.) ≤ 0.01, WT: 8,552 RegC, ΔICP0: 6,740, ΔICP22: 5,710, ΔICP27: 7,188, Δ*vhs*: 6,860). These represented between 3,129 (ΔICP22 infection) and 4,391 (WT) genes (**Sup. Fig 1c**). The lower fraction of statistically significant differential regions in ΔICP22 infection is likely due to the relatively low number of ATAC-seq reads mapping to the host genome compared to the other infections (∼3.8-fold fewer, see **Sup. Table 1** in supplemental material). The main reason for this was the high fraction of reads mapping to the HSV-1 genome in ΔICP22 infection (70-94%) compared to WT infection (∼52%). We confirmed the higher proportion of viral reads in ΔICP22 infection compared to WT in an independent experiment for both 8 and 12 h infection with the ΔICP22 mutant and its parental WT strain F (WT-F, **Sup. Table 1**). Previously, McSwiggen *et al.* showed that the viral genome remains largely nucleosome-free and thus highly accessible (14). In contrast, the human genome is predominantly inaccessible except at promoters, gene bodies of highly expressed genes and enhancers (39). Accordingly, ATAC-seq coverage on the HSV-1 genome is essentially uniform (**Sup. Fig 2a,b**). The exception are the inverted repeat regions as we masked the terminal repeat copies from read alignment. Thus, the internal repeats exhibit approximately twice the coverage of the unique HSV-1 genome regions. The flat ATAC-seq coverage is observed for all null mutants, indicating that viral chromatin accessibility is not dependent on individual viral proteins.

**Fig 1a** visualizes the positions of identified RegC for the 4,981 promoter windows containing at least one statistically significant (adj. p. ≤ 0.01) differential region for at least one virus infection compared to mock. Log2 fold-changes for statistically significant regions are illustrated in **Sup. Fig 1d**. Here, blue indicates that the relative read density in that subregion is increased during virus infection compared to mock (infection>mock, denoted as i-RegC in the following), and red indicates that relative read density in that subregion is higher in mock compared to infection (mock>infection, denoted as m-RegC). Strikingly, our results showed a highly homogeneous picture for WT and null-mutant infections compared to mock, with changes in the same direction observed at approximately the same locations. To identify distinct patterns of changes in chromatin accessibility, we performed hierarchical clustering on the RegC location heatmap in **Fig 1a**. We selected a cutoff on the dendrogram to obtain 14 clusters identified by visual inspection. To visualize the average read density for each cluster, we performed metagene analyses of promoter windows combined with statistics on RegC locations for each cluster (**Fig 1b,d,f,g, Sup. Fig 3**). Each 6 kb input window was divided into 101 bins of ∼59 bp for metagene analyses. For each bin, ATAC-seq read counts were determined, normalized to sequencing depth, normalized to sum up to 1 for each input window, averaged across all windows in each cluster, and then averaged across replicates. In addition, we show which fraction of windows have an m-(red) or i-RegC (blue) for each bin. Based on the location heatmap and the metagene analyses, we identified three major patterns of changes in chromatin accessibility around promoters during WT infection: (I) ATAC-seq peaks at the TSS that shift and/or broaden into regions downstream of the TSS during HSV-1 infection (clusters 1, 2, 4, 5, 9, 10, 12, 14, a total of 3,472 genes, **Fig 1b, Sup. Fig 3a-g**, examples in **Fig 1c, Sup. Fig 4a-h**), (II) ATAC-seq peaks at the TSS that shift and/or broaden into regions upstream of the TSS (clusters 6, 7, 11, a total of 919 genes, **Fig 1d, Sup. Fig 3h,i**, examples in **Fig 1e, Sup. Fig 4i-k**), and (III) an increase in chromatin accessibility both up- and downstream of the TSS peak in infection compared to mock (cluster 13, 126 genes, **Fig 1f**, example in **Sup. Fig 4l**). Two clusters (3 and 8, a total of 464 genes) exhibited a combination of patterns I and II with an extension of the TSS peak in both up- and downstream direction (**Fig 1g, Sup. Fig 3j, Sup. Fig 4m,n**).

**Fig 1:**
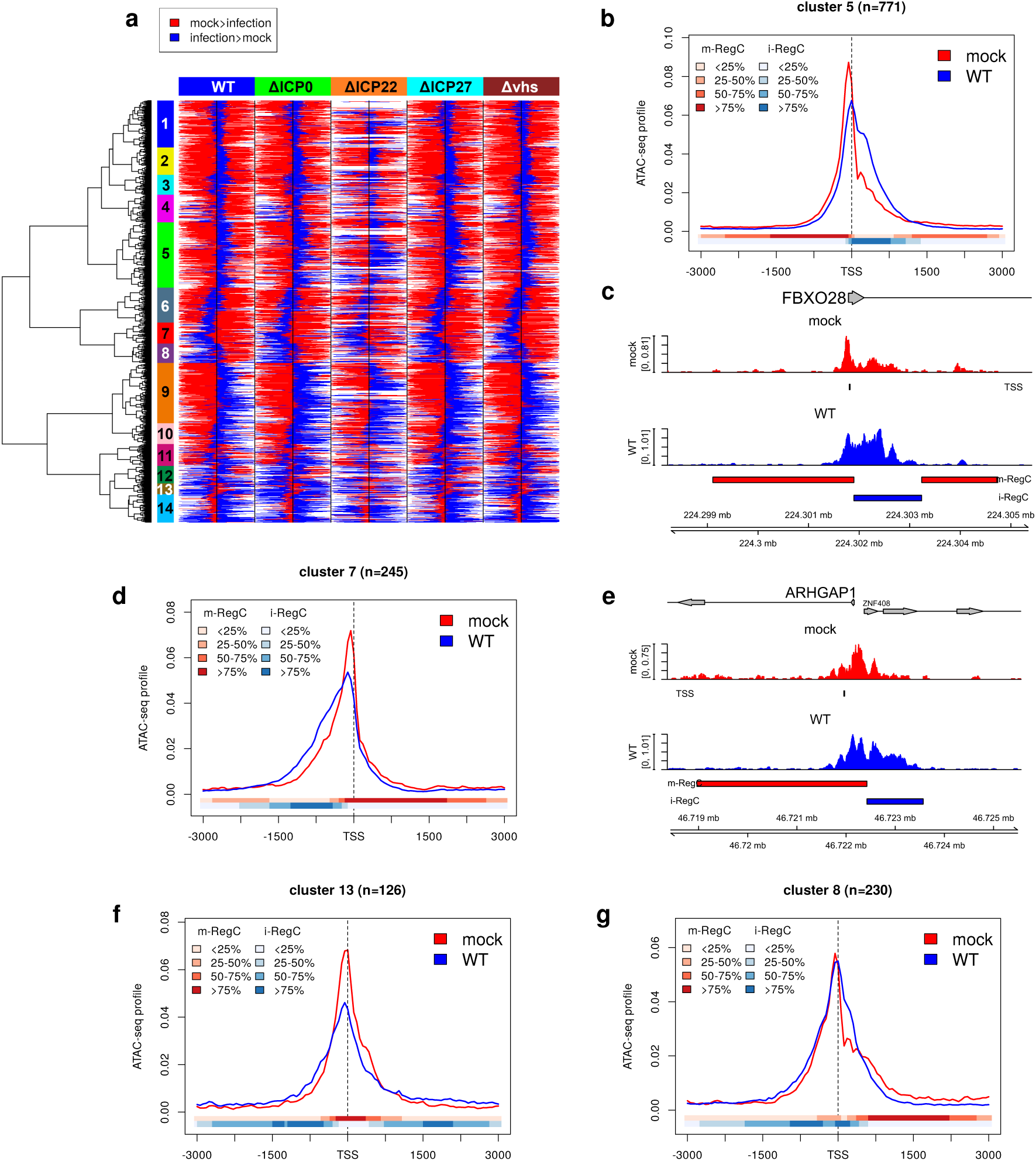
HSV-1 infection impacts chromatin accessibility around promoters for the majority of host genes. **(a)** Heatmap illustrating the location of differential regions (m- and i-RegC) identified by RegCFinder on the ATAC-seq data for WT, ΔICP0, ΔICP22, ΔICP27, and Δ*vhs* infection compared to mock infection. Results are shown for 4,981 (out of 7,649) promoter windows containing at least one statistically significant (adj. p. ≤ 0.01) differential region for at least one virus infection compared to mock. Red and blue colors represent m-RegC (mock>infection) and i-RegC (infection>mock) locations, respectively, within promoter windows. Here, each heatmap row shows results for the same input window in different comparisons of virus infection to mock. Colored rectangles on top of the heatmap indicate which virus infection was compared to mock. Black vertical lines in the center of each part of the heatmap indicate the position of the TSS. Hierarchical clustering was performed according to Euclidean distances and Ward’s clustering criterion, and the cutoff on the hierarchical clustering dendrogram was selected to obtain 14 clusters (marked by colored rectangles between the dendrogram and heatmap and numbered from top to bottom as indicated). Log2 fold-changes for differential regions are shown in **Sup. Fig 1d. (b)** Metagene plot showing the average ATAC-seq profile around the TSS ± 3 kb in mock (red) and WT (blue) infection for cluster 5, an example for pattern I. For a description of metagene plots, see Materials and Methods. The colored bands below the metagene curves in each panel indicate the percentage of genes having an m- or i-RegC (red or blue, respectively) at that position. **(c)** Read coverage in a ±3.6 kb window around the TSS in ATAC-seq data of mock (red) and WT (blue) infection, for example, gene FBXO28 with pattern I. Read coverage was normalized to the total number of mapped reads for each sample and averaged between replicates. A short vertical line below the read coverage track for mock infection indicates the TSS. Gene annotation is indicated at the top. Boxes represent exons, lines represent introns, and the direction of transcription is indicated by arrowheads. Below the read coverage track for WT infection, m-RegC (red bars) and i-RegC (blue bars) are shown for the comparison of WT vs. mock infection. **(d)** Metagene plot as in **(b)** for cluster 7, the most pronounced case of pattern II. **(e)** Read coverage plot as in **(c)** for example gene ARHGAP1 with pattern II. Here, the promoter window also contains the TSS of the ZNF408 gene on the opposite strand, and the accessible chromatin region is extended upstream of the ARHGAP1 TSS, i.e., downstream of the ZNF408 TSS. **(f,g)** Metagene plots as in **(b)** for **(f)** cluster 13, which exhibits pattern III, and **(g)** cluster 8, one of two clusters exhibiting the combined I+II pattern.

### Most chromatin accessibility changes at promoters are independent of dOCR induction

First, we evaluated which observed changes were due to dOCRs extending into promoter regions. For this purpose, we determined the fraction of promoter windows in each cluster that overlapped with dOCRs of the 1,296 genes for which we previously showed consistent dOCR induction across different HSV-1 strains (29) (**Fig 2a**). dOCR regions in mock, WT and all null mutant infections (average across two replicates) were identified as previously described (21, 29) (see also Materials and Methods). It is important to note that some relatively short dOCRs are already observed in mock infection, but these substantially extend during HSV-1 infection in read-through regions. In this analysis, we also included ATAC-seq data obtained in the same experiment for 8 h p.i. WT infection combined with PAA treatment (WT+PAA) to identify dOCR regions that are not as clearly detectable without PAA treatment but may still bias promoter analyses. WT (± PAA) and null mutant infections are ordered according to the overall extent of dOCR induction in **Fig 2a**. While no differences to mock infection were observed for ΔICP22 infection as expected, some clusters showed enrichment for dOCR overlaps upon WT(± PAA), ΔICP0, ΔICP27, and Δ*vhs* infection. This increased with the overall extent of dOCR induction in these viruses. Notably, since read-through in ΔICP27 infection is strongly reduced, dOCR induction is also reduced – but not abolished – as dOCRs require strong read-through transcription.

**Fig 2:**
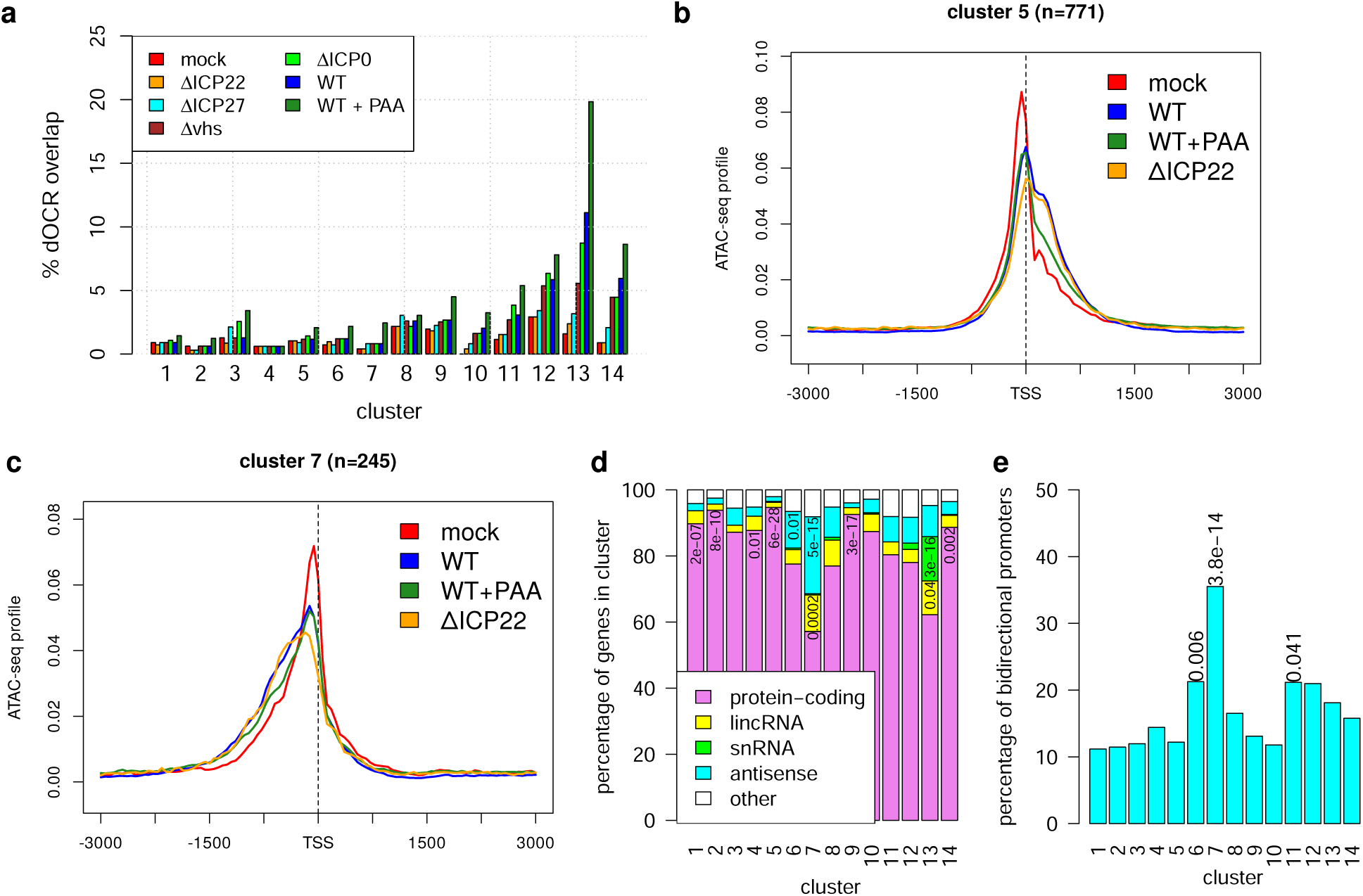
Changes in chromatin accessibility at promoters are mostly independent of dOCR induction. **(a)** Bar plots showing the percentage of promoter regions for each cluster overlapping with dOCRs downstream of the 1,296 genes for which we previously showed consistent dOCR induction across different HSV-1 strains (29). dOCR regions in mock, WT (± PAA treatment), and null mutant infection were calculated as previously described (29) (see Materials and Methods for further details), and the overlaps of promoter windows to dOCR regions originating from an upstream gene were determined for each cluster. WT (± PAA treatment) and null mutant infection are ordered according to the overall extent of dOCR induction. Only a few clusters (12–14) showed some enrichment for dOCRs from upstream genes, which was abolished in ΔICP22 infection. **(b-c)** Metagene curves showing average ATAC-seq profiles in promoter windows for mock (red), WT (blue), WT+PAA (green), and ΔICP22 (orange) infection for clusters 5 **(b)** and 7 **(c)**. (**d**) Barplot showing the percentage of genes in each cluster annotated as either protein-coding, long intervening/intergenic noncoding RNAs (lincRNA), snRNA, antisense, or others in Ensembl. The significance of enrichment of each gene type in each cluster was determined using a one-sided Fisher’s exact test (with alternative = greater), and p-values were corrected for multiple testing using the Benjamini-Hochberg method. Adj. p. < 0.05 are indicated in the corresponding field of the barplot. (**e**) The percentage of bidirectional promoters in each cluster was calculated as the percentage of promoter windows containing a protein-coding, lincRNA, or antisense gene (according to Ensembl annotation) on the opposite strand to the target gene starting within 1 kb upstream of the TSS of the target gene. Enrichment and significance analysis and multiple testing correction were performed as for (**d**) and adj. p. < 0.05 are indicated on top of bars.

However, no cluster had more than 20% of promoter windows overlapping with dOCRs even with PAA treatment, and most had <5% overlap, indicating that dOCRs represent only a very small fraction of observed changes in chromatin accessibility around promoters. Most importantly, the extension of accessible chromatin regions at promoters was also observed upon infection with an ICP22-null mutant. Moreover, in contrast to dOCRs, which increase upon PAA treatment, the extension of open chromatin at promoters was strongly reduced by PAA treatment, indicating that it is not linked to dOCRs (**Fig 2b,c**, **Sup. Fig 5-18**). Even for clusters 12-14, which exhibited some enrichment for dOCRs, most differential regions were also detected in ΔICP22 infection (**Sup. Fig 16-18**) and thus not linked to dOCR induction. This was confirmed in the independent ATAC-seq experiment for 8 and 12 h infection with WT-F and its ΔICP22 mutant, which also confirmed the chromatin changes at promoters for a different HSV-1 strain (**Sup. Fig 19**). Notably, there was little change between 8 and 12 h infection in both WT-F and ΔICP22 infection. The absence of ICP0, ICP27, or *vhs* also had little impact on changes in chromatin accessibility around promoters (**Sup. Fig 5-18**) despite the longer duration of infection for the ICP0-null mutant and the different parental virus strains for the null mutants. These observations suggest that there is an upper limit to the extent of chromatin accessibility changes at promoters in HSV-1 infection. We conclude that neither dOCR induction nor the activity of ICP0, ICP22, ICP27, or *vhs* alone explains most observed changes in chromatin accessibility at promoters in HSV-1 infection.

### Extension of accessible chromatin regions around promoters is linked to transcription

To correlate observed changes to potential transcription factor binding, we next performed motif discovery for novel and known motifs in m- and i-RegC grouped either by cluster or pattern compared to the background of all promoter windows using HOMER (40) but found no enriched motifs. Functional enrichment analysis for Gene Ontology (GO) terms for each cluster also yielded no significant results, except for cluster 13 (pattern III). Cluster 13 was enriched for “mRNA splicing via the spliceosome” and related terms, however, this was mainly due to several snRNA genes in this cluster. Accordingly, cluster 13 was significantly enriched for snRNA genes (**Fig 2d**, adj. p. = 3 × 10^−16^, see Materials and Methods). In contrast, pattern I clusters tended to contain high fractions of protein-coding genes but no or very few snRNAs. Manual investigation of these few snRNAs either showed no shift or an overlap with other protein-coding genes. Thus, most snRNAs either showed no significant changes or a relative increase in chromatin accessibility on both sides of the TSS (= pattern III in cluster 13) but no down- or upstream shifts in chromatin accessibility. Notably, snRNAs are transcribed by RNA Polymerase III (Pol III), not Pol II. However, snRNA loci and other genes not transcribed by Pol II (rRNAs, tRNAs) are repeated several times in the human genome. The quality of read mappings to these loci is thus insufficient to draw a definitive conclusion. Consequently, few snRNA (55), almost no rRNA (12) and no tRNA loci were included in our analysis. In summary, this analysis provides clear evidence for an extension of accessible chromatin at promoters of many genes transcribed by Pol II, particularly protein-coding genes.

Strikingly, cluster 7 (= the most pronounced case of pattern II) was significantly enriched for antisense transcripts (adj. p. = 5 × 10^−15^). The other pattern II clusters 6 and 11, as well as cluster 8 (combined pattern I + II), also contained a relatively high fraction of antisense transcripts. Consistently, we found that cluster 7 (adj. p. = 3.8 × 10^−14^) and to a lesser degree clusters 6 and 11 (adj. p. = 0.006 and 0.041, respectively) were enriched for bidirectional promoters, i.e., promoters containing an annotated gene start within 1 kb upstream of the TSS of the target gene (= the gene around whose TSS the promoter window was originally defined, **Fig 2e**, see **Fig 1e** for an example). We thus hypothesized that pattern II essentially represented the mirror image of pattern I (mirrored at a vertical axis through the TSS) with the “downstream” broadening/shift of chromatin accessibility occurring in antisense direction for bidirectional promoters. To confirm this, we calculated the sense-to-antisense transcription ratio in promoter windows for all genes. For this purpose, we analyzed RNA-seq data of chromatin-associated RNA, which depicts nascent transcription, for mock and 8 h p.i. WT HSV-1 infection from our recent study (19). We also included in this analysis the 2,668 genes (denoted as NA group) without significant chromatin accessibility changes that were excluded from **Fig 1a**. Interestingly, metagene analyses for these genes also revealed a slight broadening of the TSS peak into downstream regions. However, this was much less pronounced than for pattern I clusters and, therefore, not detected by RegCFinder (**Sup. Fig 20a**). To calculate the sense-to-antisense transcription ratio, promoter windows were divided into the regions down- and upstream of the TSS, and expression in chromatin-associated RNA was determined in sense direction for the downstream region (=DSR) and in antisense direction for the upstream region (=UAR). We then compared log2 ratios of DSR to UAR expression between clusters (**Fig 3a)**. Positive log2(DSR:UAR) ratios indicate that sense transcription downstream of the TSS is stronger than antisense transcription upstream of the TSS, and negative values indicate the opposite. Strikingly, all pattern II clusters had significantly lower DSR to UAR ratios than the remaining genes, while pattern I tended to have considerably higher ratios. While for pattern II clusters 6 and 11 median values were still positive, values for cluster 7 with the most pronounced pattern II were commonly negative. Thus, transcription upstream of the TSS in antisense direction was the dominant mode of transcription for cluster 7 promoters, which means pattern II essentially just represents pattern I for the genes on the antisense strand.

**Fig 3:**
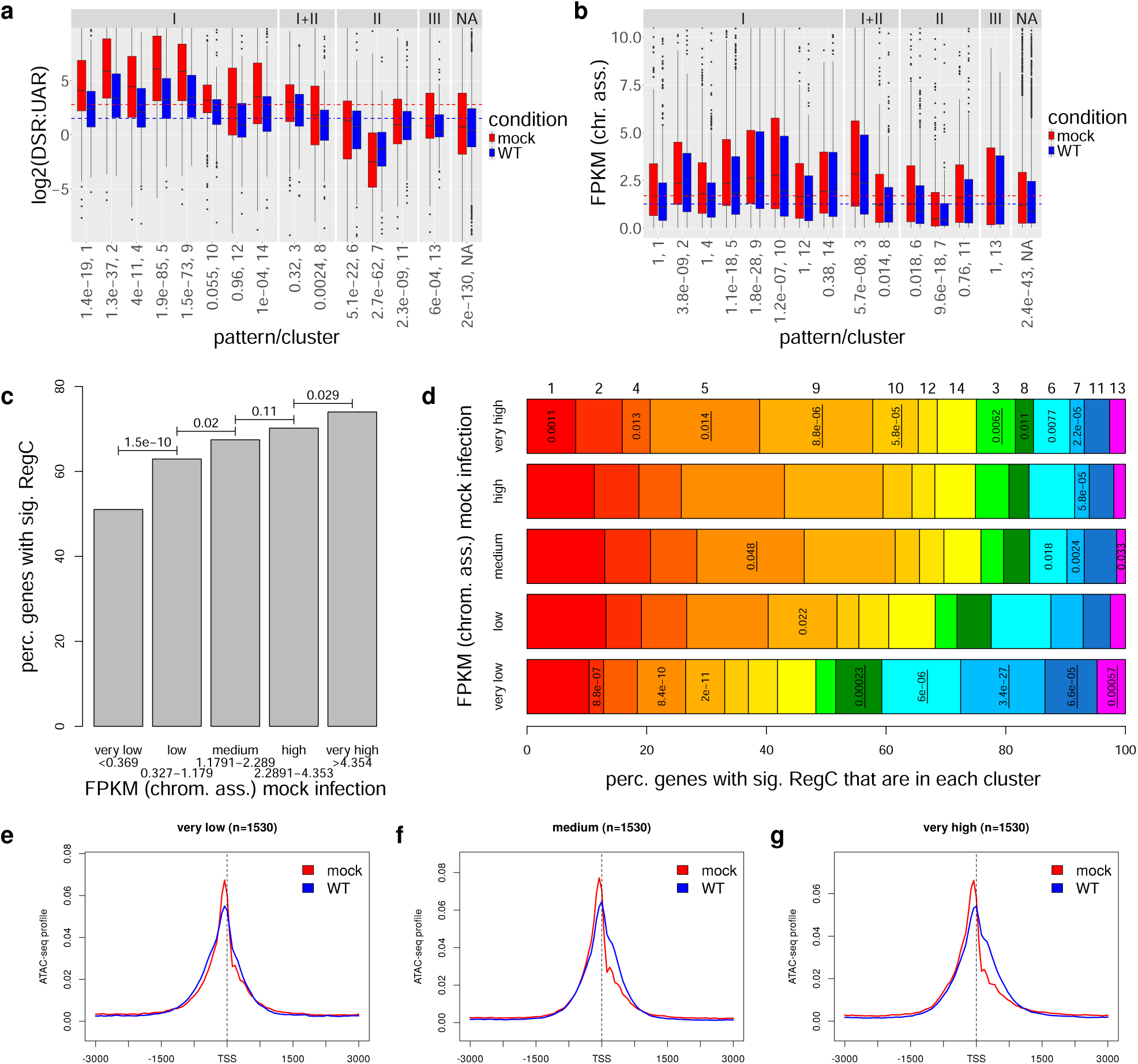
Changes in chromatin accessibility at promoters are linked to transcription. **(a,b)** Boxplots showing the distribution of **(a)** log2(DSR:UAR) ratios and **(b)** gene expression (FPKM) values in chromatin-associated RNA in mock (red) and 8 h p.i. WT (blue) HSV-1 infection for all clusters (grouped by pattern, cluster numbers are shown below each boxplot) and remaining genes without significant chromatin accessibility changes (NA group). DSR was calculated as the expression in chromatin-associated RNA for the region downstream of the TSS in the sense direction of the target gene and UAR as the expression in the upstream region in the antisense direction. P-values for Wilcoxon rank sum tests comparing values in mock infection for each cluster against all other analyzed genes are indicated below cluster numbers and were corrected for multiple testing using the Bonferroni method. **(c)** Barplot showing the percentage of genes with very low, low, medium, high and very high gene expression with at least one significant differential region (RegC). Here, gene expression cutoffs for the five groups were determined such that each group contains the same number of genes and are indicated below the labels on the x-axis. Multiple testing corrected p-values for two-sided Fisher’s exact tests comparing the fraction of genes with RegC between subsequent gene expression groups are indicated on top of bars. **(d)** Percentage of genes within each cluster among those genes with at least one significant RegC for the five gene expression groups from **(c)**. Significance of enrichment or depletion for each cluster in each gene group was determined with two-sided Fisher’s exact tests and multiple testing correction was performed with the method by Benjamini and Hochberg. Adj. p. < 0.05 are shown and are underlined in case the cluster is enriched and not underlined if it is depleted. Clustered are ordered and colored according to pattern (red-yellow: pattern I, green: pattern I + II, blue: pattern II, magenta: pattern III). **(e-g)** Metagene curves showing average ATAC-seq profiles in promoter windows for mock (red) and WT infection (blue) for genes with very low, medium, and very high expression from **(b)**.

Next, we investigated differences in gene expression changes in WT compared to mock infection between patterns (**Sup. Fig 20b**). This showed a significant difference (adj. p < 0.001) for cluster 1, for which gene expression was more strongly down-regulated than for all other genes. No other consistent trends were observed between the different patterns. In contrast, gene expression (quantified as FPKM = fragments per kilobase million mapped reads, **Fig 3b**) differed considerably between patterns. Most pattern I clusters as well as cluster 3 (combined pattern I+II) exhibited median expression levels above average in mock (**Fig 3b**), which was statistically significant for 5 clusters. In contrast, clusters 6, 7 (pattern II) and 8 (combined pattern I+II) exhibited relatively low expression values, with cluster 7 genes having significantly lower expression. The latter is consistent with these genes being less expressed than their antisense counterpart in these bidirectional promoters. Genes without significant chromatin accessibility changes (NA group) also showed significantly lower expression, albeit not as low as cluster 7. These differences between clusters/patterns were generally maintained in HSV-1 infection at lower overall expression levels, consistent with the absence of differences in gene expression fold-changes between most clusters. When stratifying analyzed genes into five equal-sized groups according to FPKM in chromatin-associated RNA in uninfected cells, we observed that the fraction of genes with at least one significant RegC increased with gene expression (**Fig. 3c**). Consistently, pattern II was significantly enriched among the lowliest expressed genes and depleted among genes with medium to high expression (**Fig. 3d**). In contrast, cluster 5, 9 and 10 with pattern I were enriched among the most highly expressed genes. Metagene analyses on the five gene expression groups confirm this trend, with the downstream extension of chromatin accessibility increasing with gene expression (**Fig. 3e-g**). Interestingly, pattern I cluster 1, which showed a significantly higher reduction in gene expression during HSV-1 infection than the other analyzed genes, was most frequent among genes with low to high expression but weakly significantly depleted among the most highly expressed genes. In summary, these results indicate that HSV-1 infection extends the accessible chromatin around promoters in the dominant direction of transcription for most host genes. Here, highly expressed genes and moderately expressed genes with stronger transcription reduction in HSV-1 infection are most strongly affected. Notably, although highly expressed genes do not generally exhibit a stronger reduction of transcription *relative* to their original expression levels in mock, the *absolute* drop in Pol II occupancy between mock and HSV-1 infection is more pronounced than for more lowly expressed genes. Thus, our results show parallels to observations for yeast that Pol II depletion induces downstream shifts of +1 nucleosomes, which extends nucleosome-free regions and accordingly accessible chromatin at promoters, most prominently for highly expressed genes (30). We thus hypothesized that the loss of Pol II during HSV-1 infection causes the extension of accessible chromatin at promoters during HSV-1 infection.

### Changes in chromatin accessibility manifest between 4 and 6 h of HSV-1 infection

To investigate how early in infection changes in chromatin accessibility around promoters can be detected, we ran RegCFinder on an ATAC-seq time-course for 1, 2, 4, 6, and 8 h p.i. WT infection from our previous publication (21) (**Fig 4a,b**). While barely any significant differential regions were identified at 1 and 2 h p.i., 1,963 significant regions in 1,249 promoter windows were identified by 4 h p.i. and 7,558 significant regions in 3,989 promoter windows by 8 h p.i. (**Fig 4b**). Log2 fold-changes for significant differential regions generally reflected the patterns observed in the analysis for WT and null mutant infections (**Fig 4a**). The same applied when we separately determined fold-changes and significance on the time-course data for the differential regions determined for WT vs. mock from our initial analysis (**Sup. Fig 21**) and when we performed metagene analyses for the clusters from **Fig 1a (Fig 4d-f**, **Sup. Fig 22**, example genes in **Sup. Fig 23**). Although the changes in chromatin accessibility at 8 h p.i. of the time-course were less pronounced than for the original WT vs. mock comparison at 8 h p.i., this nevertheless confirms the changes in chromatin accessibility around promoters in HSV-1 infection in an independent experiment. The lower fraction of viral reads at 8 h p.i. in the time-course experiment (∼25%, **Sup. Table 1**) than both for the first WT experiment (∼52%) and WT-F at 8 h (∼61%) suggests a slower progression of infection in the time-course experiment, which likely explains why the effect was less pronounced. Again ATAC-seq read coverage on the HSV-1 genome remained essentially uniform throughout the time-course (**Sup. Fig 2c,d**).

**Fig. 4:**
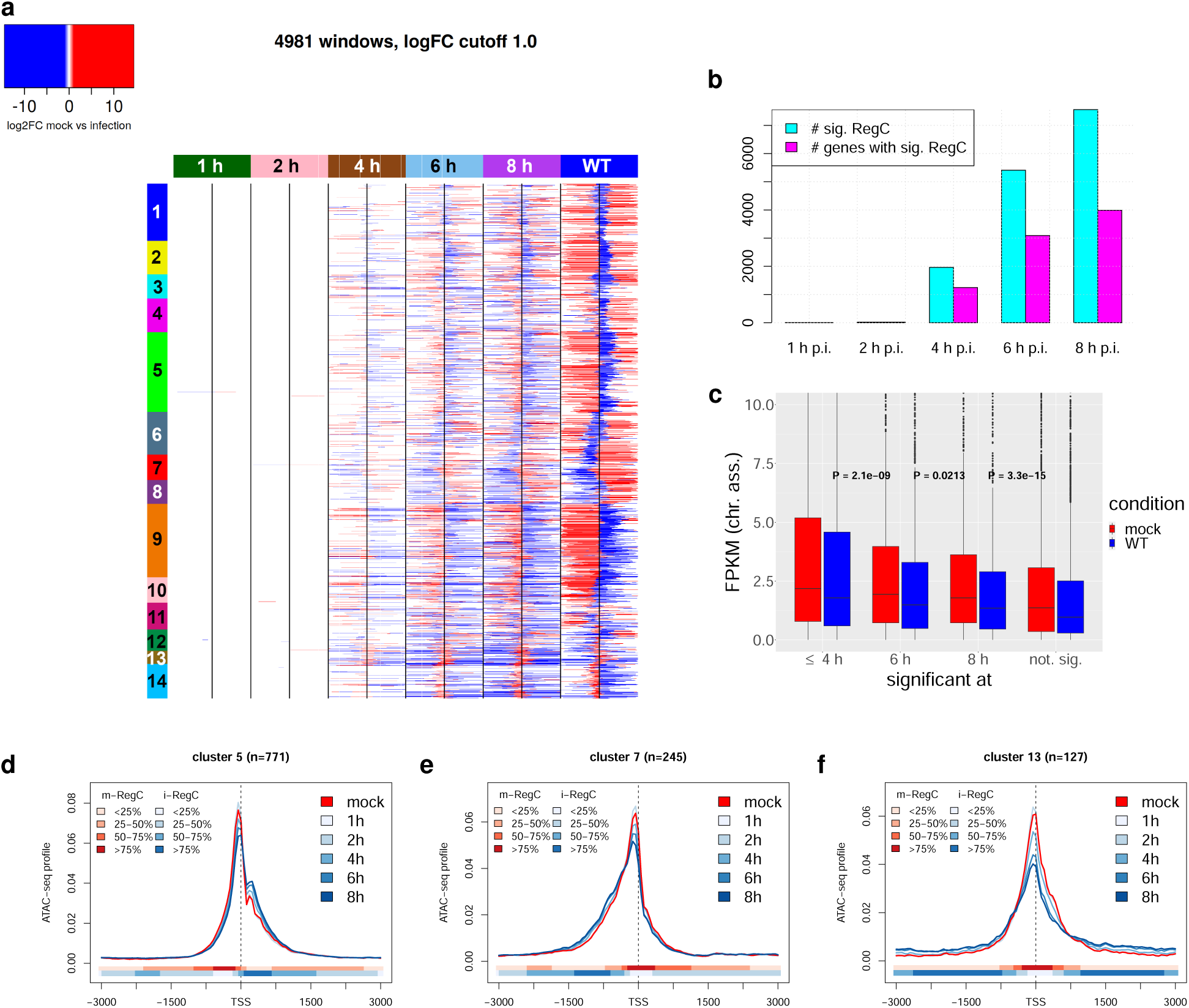
Changes in chromatin accessibility begin to manifest between 4 and 8 h p.i. depending on gene expression. **(a)** Heatmap showing the log2 fold-changes determined with DEXSeq on the ATAC-seq time-course data (1, 2, 4, 6 and 8 h p.i. WT infection compared to mock infection) for statistically significant differential regions (m- and i-RegC) identified by RegCFinder. For comparison, log2 fold-changes for the WT vs. mock comparison from Fig 1a are also shown. Each row represents the results for one of the input windows included in Fig 1a. Colored rectangles on top indicate the time-point of infection or whether the original WT vs. mock comparison is shown. Statistically significant differential regions are colored according to the log2 fold-change determined by DEXSeq. Here, the color scale is continuous between −1 and 1, and all log2 fold-changes >1 are colored the same red, and all log2 fold-changes <1 the same blue. Promoter windows are ordered as in Fig 1a, and clusters are annotated by colored and numbered rectangles on the left. **(b)** Number of statistically significant differential regions (m- and i-RegC) and number of genes with at least one statistically significant differential region identified for each time-point of infection in the ATAC-seq time-course data. **(c)** Boxplot showing the distribution of gene expression (FPKM) values in chromatin-associated RNA for mock and 8 h p.i. WT infection for genes with significant changes in chromatin accessibility in promoter windows (i) at 4 h p.i. or earlier, (ii) at 6 h p.i. but not yet at 4 h p.i., (iii) at 8 h p.i. but not yet at 6 h p.i. and (iv) in the analysis shown in Fig 1a, but not yet at 8 h p.i. in the time-course ATAC-seq experiment. P-values for Wilcoxon rank sum tests comparing FPKM values in mock infection between subsequent groups are also indicated. **(d-f)** Metagene curves of ATAC-seq profiles in mock infection (red) and all time-points of infection (blue shades) from the time-course experiment, for example, clusters 5 (pattern I, **d**), 7 (pattern II, **e**) and 13 (pattern III, **f**). The colored bands below the metagene curves indicate the percentage of genes having an i- or m-RegC (blue or red, respectively) at that position in the comparison of 8 h p.i. to mock infection from the time-course experiment. Metagene plots for all clusters are shown in **Sup. Fig 22**.

The time-course analysis also reveals that these changes begin to manifest by 4 h p.i., with 1,249 genes already showing a significant change. Moreover, genes that showed an early effect (by 4 h p.i. or earlier) were significantly more highly expressed than genes that showed an effect by 6 h p.i. or later (**Fig 4c**). The lowest expression was observed for genes with a significant effect in the first WT experiment but not in the time-course. This again confirms the link between expression levels of a gene and the change in chromatin accessibility. Notably, we previously showed that transcriptional activity on host genes drops substantially in the first 4 h of infection to only 40% of transcription in uninfected cells, which was further halved until 8 h p.i. (19). Thus, the onset of the extension of accessible chromatin regions at promoters into downstream (for pattern I) and upstream (for pattern II) regions follows the onset of the drop in host transcription during infection. This provides further evidence for the hypothesis that loss of Pol II from host genes drives changes in chromatin accessibility at promoters. Since the slower progression of virus infection in the time-course experiment likely leads to a less pronounced loss of Pol II from host genes, this would explain the differences in effect between the time-course and the null mutant experiment.

### ICP4 knockout reduces, and combined expression of ICP22 and ICP27 induces the extension of accessible chromatin

We already showed above that PAA treatment substantially reduced the broadening and/or downstream shift of accessible chromatin regions at promoters (**Fig 2b,c, Sup. Fig 5-18**). This was also observed for 8 and 12 h p.i. PAA treatment of WT-F in our independent ATAC-seq experiment with little differences between the two time-points (**Sup. Fig 24**). Control experiments with mock ± PAA at 8 and 12 h showed no changes in chromatin accessibility at promoters (**Sup. Fig 25**). We previously found that PAA alleviates the depletion of Pol II from host genomes by inhibiting viral DNA replication (29). This provides further evidence that Pol II depletion leads to the observed shifts in chromatin accessibility at promoters. The original ATAC-seq experiment with null mutant infections also included infection with an ICP4 null mutant at 8 h p.i., which had not been previously published. Since ICP4 facilitates recruitment of Pol II to viral replication compartments (13, 14) and Pol II depletion from host promoters is not observed in ΔICP4 infection (13, 41), we now also investigated changes in chromatin accessibility in ΔICP4 infection. Strikingly, while we still observed the extension of accessible chromatin in down-(**Fig 5a**, **Sup. Figs 26a-g,k,l**) or upstream direction (**Fig 5b**, **Sup Fig. 26h,i**), it was similarly reduced in ΔICP4 infection compared to WT infection as upon PAA treatment. The fraction of viral reads in ATAC-seq experiments was comparably low at 3-4% in both ΔICP4 and WT+PAA infection (**Sup. Table 1**). This is consistent with previous reports that there is no viral DNA replication in the absence of ICP4 (42, 43). Moreover, high chromatin accessibility in HSV-1 infection has been shown to be independent of ICP4 (13) and ATAC-seq read coverage remained essentially uniform (**Sup. Fig 2a,b**), thus low ATAC-seq read numbers in ΔICP4 infection are not due to reduced chromatin accessibility of viral genomes.

**Fig 5:**
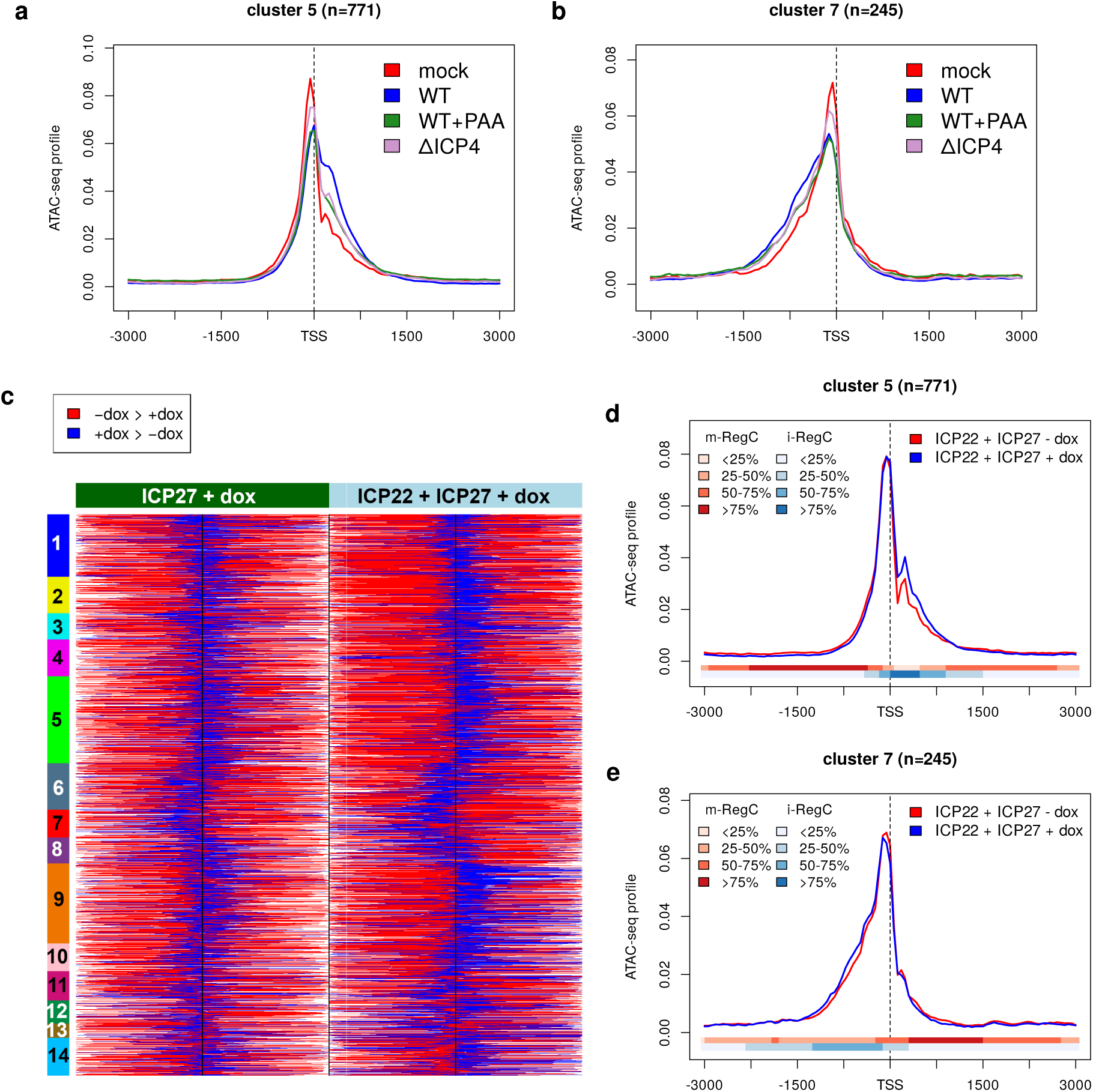
Chromatin accessibility changes upon ICP4 knockout and dox-induced combined ICP22 and ICP27 expression. **(a-b)** Metagene curves showing average ATAC-seq profiles in promoter windows for mock (red), WT (blue), WT+PAA (green) and ΔICP4 (violet) infection for clusters 5 **(a)** and 7 **(b)**. Metagene plots for all other clusters are shown in **Sup. Fig 26**. **(c)** Heatmap showing the location of differential regions (m- and i-RegC) identified by RegCFinder for T-HF-ICP27 cells and T-HF-ICP22/ICP27 cells upon dox exposure. Each row represents the results for one of the input windows included in Fig 1a. Promoter windows are ordered as in Fig 1a and clusters are annotated as colored and numbered rectangles on the left. Log2 fold-changes for differential regions are shown in **Sup. Fig 27b**. **(d-e)** Metagene curves showing average ATAC-seq profiles in promoter windows for T-HF-ICP22/ICP27 cells with (blue) and without (red) dox exposure for clusters 5 **(d)** and 7 **(e)**. The colored bands below the metagene curves indicate the percentage of genes having an m-RegC (red, decreased upon dox exposure) or i-RegC (blue, increased upon dox exposure) or at that position. Metagene plots for all clusters are shown in **Sup. Fig 27**.

We next tested for enrichment in promoter regions of ICP4 binding sites on the human genome previously identified with ChIP-seq by Dremel *et al.* (41). This showed a significant enrichment (adj. p. = 1.1 × 10^−5^) of ICP4 binding sites for genes with significant RegC (compared to the NA group of genes without any significant changes) and among these, a weakly significant enrichment in clusters 9 (pattern I, adj. p = 0.0046) and 11 (pattern II, adj. p.=0.0018). Notably, the only cluster with less frequent ICP4 binding than the NA group was cluster 13 (pattern III), which did not show shifts in chromatin accessibility. Thus, promoters of genes with significant shifts in chromatin accessibility are more frequently bound by ICP4 than genes without shifts, providing evidence that Pol II depletion from host genomes by ICP4 contributes to this phenomenon. However, since PAA does not inhibit ICP4 synthesis (27), the strongly reduced chromatin changes at promoters in WT + PAA infection indicate that in absence of viral DNA replication ICP4 activity is not sufficient to induce the full extent of chromatin accessibility changes.

Nevertheless, the question remains why reduced changes in chromatin accessibility are observed even in absence of ICP4. A possible explanation is that activities of other immediate-early proteins, which are still expressed and active in the absence of ICP4 (42–44), lead to an effective depletion of Pol II from host gene bodies. Two candidates for this are ICP27, which redirects Pol II transcription from gene bodies into intergenic regions by disrupting transcription termination (20), and ICP22, which inhibits host transcription elongation (15). In particular, HSV-1-induced disruption of transcription termination has a massive effect on remaining host transcriptional activity, with ∼50% of newly transcribed RNA reads originating from intergenic regions (20). To investigate whether ICP27 and/or ICP22 expression alone can induce such changes, we re-analyzed our recently published Omni-ATAC-seq (a recent improvement of ATAC-seq (45)) data for telomerase-immortalized human foreskin fibroblasts (T-HFs) that express either ICP22 (T-HF-ICP22 cells) or ICP27 (T-HF-ICP27 cells) in isolation or combination (T-HF-ICP22/ICP27 cells) upon doxycycline (dox) exposure. While we did not identify any significant RegC upon ICP22 expression alone, ICP27 expression alone led to significant changes for >2,000 genes (**Fig 5c**, **Sup. Fig 27a,b**). Combined ICP22 and ICP27 expression led to more pronounced results, with the same patterns observed as for WT infection but in an attenuated manner (**Fig 5c-e**, **Fig 27**). Thus, the combined activity of ICP22 and ICP27 is sufficient to induce a moderate extension of chromatin accessibility at promoters. RNA-seq analysis performed in parallel to ATAC-seq in the same experiment showed that ICP27 was not as strongly expressed in the T-HF-ICP22/ICP27 cells upon dox exposure as in the T-HF-ICP27 cells (∼3-fold less). In contrast, ICP22 was much more strongly expressed upon dox exposure in T-HF-ICP22/ICP27 cells than in the T-HF-ICP22 cells (>5-fold more). This indicates that direct effects mediated by ICP22 are important for further extending chromatin accessibility at promoters rather than any indirect effects via enhancement of ICP27 expression and thus read-through.

Nevertheless, our results showed that the pronounced changes observed in WT infection require viral DNA replication but neither ICP22 nor ICP27 expression. Notably, although ICP27 is required for optimal viral DNA replication, knockout of ICP27 does not completely abolish viral DNA replication (46). Consistent with this, ∼23% of ATAC-seq reads have a viral origin in ΔICP27 infection in contrast to only 3-4% in ΔICP4 and WT + PAA infection (**Sup. Table 1, Sup. Fig 2a,b**). This is comparable to the 8 h p.i. time-point in the time-course experiment. Furthermore, depletion of Pol II from host promoters is still observed in ΔICP22 and ΔICP27 infection (41). We conclude that the common feature between the different experimental conditions that exhibit promoter chromatin changes is the depletion of RNA Pol II from host promoter regions either by a generalized loss of Pol II or by Pol II translocation into downstream genomic regions upon disruption of transcription termination.

### HSV-1 infection induces a downstream shift of +1 nucleosomes

The HSV-1-induced changes in chromatin accessibility at promoters indicate a broadening of the nucleosome-free region around promoters in either sense (pattern I) or antisense (pattern II) direction. The extension of nucleosome-free regions requires shifts in +1 or −1 nucleosome positions, and depletion of Pol II has been shown to induce a downstream shift of +1 nucleosomes in yeast (30). We, thus, performed ChIPmentation in HFF for mock and WT infection at 8 h p.i. with an antibody recognizing the C-terminal part of the non-canonical histone H2A.Z (n=3 replicates). H2A.Z is highly enriched at gene promoters at −1 and +1 nucleosome positions (32, 33) and is encoded by two genes, whose protein products H2A.Z.1 and H2A.Z.2 differ by only three amino acids. While their genomic occupancy patterns are similar, there are quantitative differences, with H2A.Z.1 being more abundant at active promoters than H2A.Z.2 (47). Since the C-terminal regions of H2A.Z.1 and H2A.Z.2 differ only by the last amino acid, the antibody used for ChIPmentation recognizes both isoforms. A metagene analysis of all analyzed promoter windows showed the expected distribution of H2A.Z occupancy in mock infection with two peaks on both sides of the TSS, corresponding to −1 and +1 nucleosome positions (**Sup. Fig 28a**).

Application of RegCFinder to H2A.Z ChIPmentation data generally identified a relative increase in H2A.Z downstream of the TSS during infection and a relative decrease upstream of the TSS for pattern I and combined pattern I+II clusters (**Fig 6a**, log2 fold-changes for differential regions shown in **Sup. Fig 28b**, examples in **Fig 6b**, **Sup. Fig 29a-g,k,l**). Metagene analyses showed that this reflected a downstream shift and broadening of +1 nucleosome peaks as well as a relative increase of +1 nucleosome peaks compared to −1 nucleosome peaks (**Fig 6d**, **Sup. Fig 28c-i,m,n**). In contrast, cluster 7 (most pronounced pattern II cluster) showed the opposite trend with relative increases in H2A.Z upstream of the TSS and decreases downstream of the TSS (example in **Fig 6c**). Consistent with pattern II representing pattern I in antisense direction, the metagene analysis showed an upstream shift and broadening of the −1 nucleosome peak and a relative increase of the −1 nucleosome peak compared to the +1 nucleosome peak (**Fig 6e**). However, this was only observed for a few genes in cluster 6 and cluster 11 (**Fig 6a**, **Sup. Fig 28j,k**, examples in **Sup. Fig 29h,i**), which fits with the observation that pattern II in the ATAC-seq data was also much less pronounced for these clusters. In contrast to pattern I and II, pattern III in the ATAC-seq data (cluster 13) was not associated with distribution changes in H2A.Z (**Sup. Fig 25l**, example in **Sup. Fig 29j**), thus it is not shaped by shifts in +1 and/or −1 nucleosome positioning. This is consistent with pattern III being at least partly associated with dOCR induction for upstream genes, which we previously linked to impaired nucleosome repositioning following Pol II transcription downstream of genes (29).

**Fig 6:**
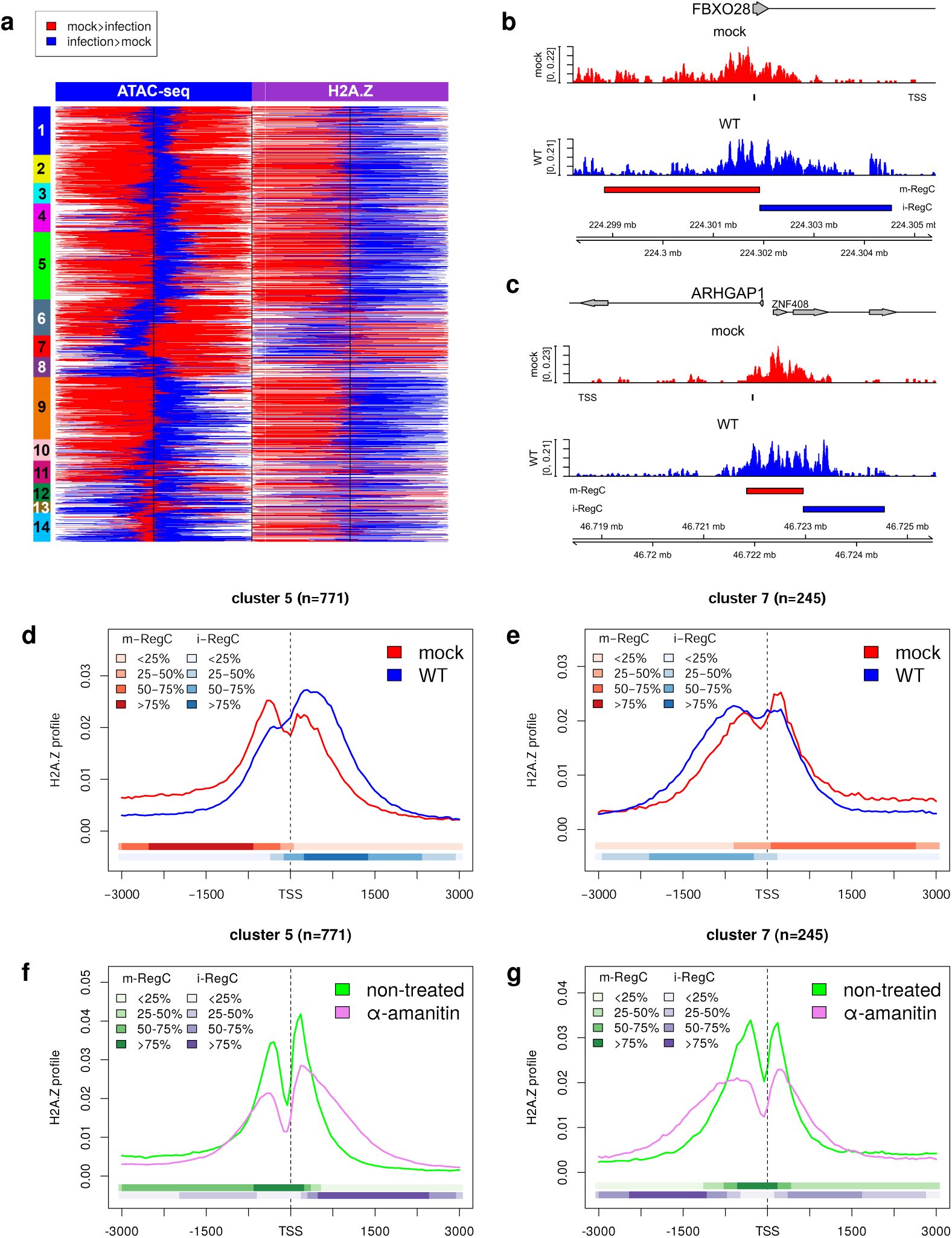
HSV-1 infection and depletion of Pol II from human genes lead to a downstream shift of H2A.Z-containing +1 nucleosomes. **(a)** Heatmap showing the location for differential regions (m- and i-RegC) identified in the WT vs. mock comparison on the ATAC-seq (left half, identical to left-most part of Fig 1a) and H2A.Z ChIPmentation data (right half) for the promoter windows included in Fig 1a. Differential regions with mock>infection (m-RegC) are marked in red, and differential regions with infection>mock (i-RegC) in blue. The order of promoter windows is the same as in Fig 1a, and clusters are indicated by colored and numbered rectangles on the left. Log2 fold-changes for differential regions are shown in **Sup. Fig 28b**. **(b,c)** Read coverage in a ±3.6 kb window around the TSS in H2A.Z ChIPmentation data for mock (red) and WT (blue) infection, for example, genes with **(b)** pattern I (FBXO28) and **(c)** pattern II (ARHGAP1). For descriptions of read coverage plots, see caption to Fig 1. **(d-g)** Metagene plots of H2A.Z profiles for mock and WT infection **(d,e)** and untreated and α-amanitin-treated HCT116 cells **(f,g)** for clusters 5 (pattern I, **d,f**) and 7 (pattern II, **e,g**). Metagene plots for other clusters can be found in **Sup. Fig 28** and **30**, respectively. The colored bands below the metagene curves in each panel indicate the percentage of genes having m- or i-RegC at that position. For **(f,g)**, m-RegC (green) are differential regions with relative read coverage higher in untreated cells and i-RegC (violet) differential regions with relative read coverage higher upon α-amanitin-treatment.

In summary, our data confirm that the broadening of chromatin accessibility around promoters results from down- and upstream shifts of +1 or −1 nucleosomes, respectively. Here, the shift direction depends on whether sense or antisense transcription represents the dominant direction of transcription at this promoter. Notably, a recent study in yeast proposed that −1 nucleosomes should be considered +1 nucleosomes for antisense transcription (48). This suggests that the loss of Pol II from host promoters during HSV-1 infection induces a shift of +1 and −1 nucleosome positions similar to what was previously observed for yeast. To confirm that inhibition of transcription alone can lead to shifts of +1 and −1 nucleosomes in human cells, we re-analyzed published H2A.Z ChIP-seq data of HCT116 cells with and without α-amanitin treatment from the recent study by Lashgari *et al*. (49). They showed that α-amanitin treatment leads to reduced Pol II levels at gene TSSs and increased incorporation of H2A.Z at TSSs of transcribed genes. Here, Pol II ChIP and RT-qPCR of control genes showed that α-amanitin treatment reduced Pol II signal at the TSS to 10-20% of untreated cells and transcriptional activity to less than 40%. This is not much lower than what has previously been reported for HSV-1 infection: Abrisch *et al.* found that Pol II occupancy on gene bodies upon 4 h HSV-1 infection was reduced on average to around 20% (9), and we previously estimated that transcriptional activity was reduced to 20-40% between 4 and 8 h p.i. (19). Notably, in contrast to DRB and flavopiridol, which inhibit transcription via arresting Pol II in a paused state at the promoter, α-amanitin induces a loss of Pol II from the host genome, thus making it a better model of HSV-1-induced Pol II depletion.

Lashgari *et al*. previously only analyzed changes in absolute H2A.Z levels at promoters and did not focus on changes in +1 and −1 nucleosome positioning. Metagene and RegCFinder analyses of the H2A.Z ChIP-seq data indeed showed a similar trend upon α-amanitin treatment as during HSV-1 infection (**Fig 6f,g, Sup. Fig 30**, examples are shown in **Sup. Fig 31**). Pattern I and pattern I+II clusters showed a massive broadening of the +1 nucleosome peak into downstream regions, even more pronounced than what is observed upon HSV-1 infection. In addition, they also exhibited a (less pronounced) broadening of −1 nucleosome peaks. Similarly, pattern II clusters showed a strong broadening of both −1 and +1 nucleosome peaks into upstream regions. In the case of cluster 7 (strongest pattern II), the broadening was much more pronounced for the −1 nucleosome (i.e., the +1 nucleosome in antisense direction) than for the +1 nucleosome (i.e., the −1 nucleosome in antisense direction). This analysis confirms that the depletion of Pol II from promoters leads to a downstream shift of +1 nucleosomes in human cells. In summary, our results demonstrate that the shift of +1 and −1 nucleosomes in the direction of transcription in HSV-1 infection is a consequence of Poll II depletion from host chromatin.

## Discussion

Previously, we showed that HSV-1 infection disrupts chromatin architecture downstream of genes with strong read-through transcription. In this study, we reveal that chromatin architecture is also substantially altered at host gene promoters during HSV-1 infection. Here, 57% of genes showed a statistically significant change in the distribution of chromatin accessibility at promoters in WT HSV-1 infection, and metagene analyses indicated similar but less pronounced changes even for genes without statistically significant changes. In essence, we identified three types of changes: Most genes showed a shift and/or broadening of accessible chromatin into regions downstream of the TSS (pattern I). In contrast, ∼900 genes showed the opposite trend with a shift and/or broadening of accessible chromatin into regions upstream of the TSS (pattern II). A further ∼460 genes exhibited a combination of patterns I and II, with the broadening of accessible chromatin in the downstream direction being more pronounced than in the upstream direction. Only a small set of genes (126 genes) showed relative increases in chromatin accessibility up- and downstream of the TSS, which could partly be attributed to overlaps with dOCRs from upstream genes and partly to an enrichment for short non-coding RNAs not transcribed by Pol II, for which promoter windows were longer than the actual gene. We thus did not further investigate this pattern.

As patterns I and II were still observed in the absence of ICP22, which is necessary for dOCR induction, these are not artifacts of dOCRs extending into downstream genes. On the contrary, PAA treatment, which increases dOCR induction, substantially reduced – though not completely abolished – patterns I and II. Furthermore, knockout of neither ICP0, ICP22, ICP27, nor *vhs* substantially affected patterns I and II. In contrast, ICP4 knockout, similar to PAA treatment, also alleviated the down- and upstream broadening of chromatin accessibility. Both PAA treatment and ICP4 knockout largely abolish viral DNA replication and consequently alleviate the depletion of Pol II from host genes, leading us to hypothesize that this HSV-1-induced loss of Pol II causes the widespread extension of open chromatin at host gene promoters. This hypothesis was further supported by the observation that combined dox-induced expression of ICP22 and ICP27 (and to a lesser degree dox-induced ICP27 expression alone) led to some broadening of chromatin accessibility at promoters despite knockout of neither of these proteins affecting the HSV-1-induced changes. Knockout of ICP22 or even ICP27 does not sufficiently abolish viral DNA replication and thus has only a minor effect on the depletion of Pol II from host genomes and, consequently, on chromatin accessibility changes. In contrast, in the absence of viral DNA replication, ICP22 and ICP27 (and potentially other viral factors) sufficiently reduce Pol II levels on gene bodies to induce an alleviated form of those chromatin accessibility changes. Notably, we determined that the more pronounced changes upon co-expression of ICP27 with ICP22 compared to expression of ICP27 alone are likely due to the direct effects of ICP22 on transcription. ICP22 interacts with P-TEFb, several transcriptional kinases, as well as elongation factors such as the FACT complex to inhibit transcription elongation of cellular genes (15). Moreover, both the relatively late onset of chromatin accessibility changes between 4 and 8 h p.i. in our time-course ATAC-seq analysis and the generally reduced levels of changes in the time-course, which had a lower fraction of viral reads at 8 h p.i., confirm that significant depletion of Pol II is necessary to observe substantial effects on chromatin accessibility. On the other hand, the little differences observed between 8 and 12 h of infection and the null mutants of ICP0, ICP22, ICP27 and *vhs* also suggest that there is an upper limit to the extent of chromatin accessibility changes at promoters in HSV-1 infection.

Analysis of transcriptional activity using RNA-seq of chromatin-associated RNA revealed that pattern I and II are indeed linked to transcription, with pattern I associated with more highly expressed genes and pattern II associated with bidirectional promoters with strong antisense transcription on the opposite strand. In the case of cluster 7, for which pattern II was most pronounced, antisense transcription was much higher than sense transcription. This was not due to the widespread induction of antisense transcription in HSV-1 infection, which we previously reported on (50), but these genes already exhibited strong transcription in antisense direction in uninfected cells originating from bidirectional promoters. Consistently, cluster 7 was enriched for genes annotated as antisense, and more than a third of these promoters contained an annotated gene starting on the opposite strand within 1 kb upstream of the TSS of the target gene. Thus, pattern II is essentially just pattern I for the genes on the opposite strand in bidirectional promoters.

Since the broadening of chromatin accessibility in promoter regions suggested an extension of nucleosome-free regions at promoters, we mapped +1 and −1 nucleosome positions by ChIPmentation of the H2A.Z histone variant enriched at these nucleosomes. Analysis of H2A.Z occupancy indeed showed a downstream shift of +1 nucleosomes for pattern I genes in HSV-1 infection and an upstream shift of −1 nucleosomes for genes with the most pronounced pattern II (cluster 7). This further confirmed that pattern II just represents pattern I for genes on the antisense strand. It is also consistent with recent results from yeast by Bagchi *et al.* that −1 H2A.Z-containing nucleosomes should be considered as +1 nucleosomes for antisense transcription (48). Downstream shifts of the +1 nucleosome to more thermodynamically favorable sites have previously been reported upon Pol II degradation in yeast by Weiner *et al*. (30), in particular for highly expressed genes. Re-analysis of published H2A.Z ChIP-seq data for α-amanitin-induced depletion of Pol II from gene promoters indicates that depletion of Pol II alone is sufficient to induce downstream shifts of +1 nucleosomes in human cells. Moreover, for genes with dominant antisense transcription (cluster 7), it resulted in upstream shifts of −1 nucleosomes. Although 24 h α-amanitin treatment – as used for ChIP-seq – potentially leads to other effects beyond losing genome-bound Pol II that may explain chromatin changes, rapid degradation of Pol II with an inducible degron system has also been shown to increase chromatin dynamics similar to α-amanitin (51). The broadening of H2A.Z peaks in both HSV-1 infection and upon α-amanitin treatment, respectively, indicate a less precise positioning of corresponding nucleosomes. This was more pronounced upon α-amanitin treatment and also observed for −1 nucleosomes, suggesting that loss of Pol II upon α-amanitin treatment is more pronounced than in HSV-1 infection.

One open question remaining concerns the biological significance of the observed changes in nucleosome positioning. Our analysis of gene expression changes did not suggest an effect on differential gene expression, rather the opposite with strong down-regulation leading to the same effect for less strongly expressed genes (cluster 1) as for highly expressed genes that are not more down-regulated than other genes. However, the downstream shifts in +1 nucleosomes may provide an explanation for the downstream shift of Pol II pausing we recently reported for HSV-1 infection (38). The +1 nucleosome was shown to play a role in promoter-proximal Pol II pausing between the promoter and the +1 nucleosome (52). In particular, the +1 nucleosome represents a 2^nd^ barrier to Pol II pause release independent of the main pausing factor negative elongation factor (NELF). Upon NELF depletion, Pol II stops at a 2^nd^ pausing region around the +1 nucleosomal dyad-associated region (53). We previously observed that Pol II pausing in HSV-1 infection is shifted to more downstream and less well-positioned sites for the majority of genes (38), consistent with +1 nucleosome positioning also appearing less well-positioned upon HSV-1 infection in our H2A.Z ChIPmentation data. Alternatively, the shifts in +1 nucleosomes might also be linked to the mobilization of H1, H2 (including H2A.Z), and H4 histones during HSV-1 infection (54–56), which has been proposed to serve as a source of histones for viral chromatin assembly. Nevertheless, even if the changes in nucleosome positioning upon Pol II depletion are of no further functional consequence for HSV-1 infection, they are highly relevant for any functional genomics studies on chromatin architecture in HSV-1 infection or other conditions that deplete Pol II from the genome. If not properly taken into account, the broadening of accessible chromatin at promoters may be mistaken, e.g., for differential transcription factor binding, which could lead to wrong conclusions. Together with read-through transcription and dOCRs extending into downstream genes, this represents one more example of how HSV-1 infection confounds standard functional genomics analyses. In any case, our study highlights that HSV-1 infection impacts chromatin architecture at promoters independently of the widespread changes downstream of genes mediated by ICP22.

## Materials and Methods

### Previously published sequencing data analyzed in this study

ATAC-seq for mock and WT HSV-1 infection (strain 17, 8 h p.i.), WT infection with 8 h PAA treatment, infection with ICP0-null mutant (ΔICP0, strain 17, (57), 12 h p.i.), ICP22-null mutant (ΔICP22, R325, strain F, (58), 12 h p.i.), ICP27-null mutant (ΔICP27, strain KOS, (59), 8 h p.i.) and *vhs*-null mutant (Δ*vhs*, strain 17, (60)) in of HFF were taken from our recent publication (29) (n=2 apart from ΔICP22 infection with n=4, GEO accession: GSE185234). This experiment also included infection with an ICP4-null mutant (ΔICP4, n12, strain KOS, (42), 8 h p.i.) which had not previously been published. ATAC-seq data for mock and WT HSV-1 infection of HFF at 1, 2, 4, 6 and 8 h p.i. WT infection (n=2 replicates, GEO accession: GSE100611) and chromatin-associated RNA-seq data for mock and 8 h p.i. WT infection (GEO accession: GSE100576) were taken from our previous publication (21). H2A.Z ChIP-seq data of untreated and α-amanitin-treated HCT116 cells were taken from the study by Lashgari *et al*. (49) (n=2, GEO accession: GSE101427).

### ChIPmentation, library preparation and sequencing

HFF were purchased from ECACC and cultured in Dulbecco’s Modified Eagle Medium (DMEM, ThermoFisher #41966052) supplemented with 10% (v/v) Fetal Bovine Serum (FBS, Biochrom #S0115), 1× MEM Non-Essential Amino Acids (ThermoFisher #11140050) and 1% penicillin/streptomycin. Two days prior to infection, two million HFF cells were seeded in 15 cm dishes. On the day of infection, cells had expanded to ∼80% confluency. Cells were infected with the respective viruses as described in the results section (n=3). At 8 p.i., cells were fixed for 10 minutes at room temperature by adding 1% formaldehyde (final) directly to the medium. Cells were scraped in 1mL of ice-cold 1x PBS containing protease inhibitor cocktail (1x) (Roche #11836153001) with an additional 1mM phenylmethylsulfonyl fluoride (PMSF). Cells were pelleted at 1500 rpm for 20 min at 4 °C. Supernatant was aspirated and cell pellets were frozen in liquid N_2_.

Cell pellets were resuspended in 1.5 mL 0.25% [w/v] SDS sonication buffer (10 mM Tris pH=8.0, 0.25% [w/v] SDS, 2 mM EDTA) with 1x protease inhibitors and 1 mM additional PMSF and incubated on ice for 10 minutes. Cells were sonicated in fifteen 1 minute intervals, 25% amplitude, with Branson Ultrasonics SonifierTM S-450 until most fragments were in the range of 200-700 bp as determined by agarose gel electrophoresis. Two million cells used for the preparation of the ChIPmentation libraries were diluted 1:1.5 with equilibration buffer (10 mM Tris, 233 mM NaCl, 1.66% [v/v] Triton X-100, 0.166% [w/v] sodium deoxycholate, 1 mM EDTA, protease inhibitors) and spun at 14,000x g for 10 minutes at 4 °C to pellet insoluble material. Supernatant was transferred to a new 1.5 mL screw-cap tube and topped up with RIPA-LS (10 mM Tris-HCl pH 8.0, 140 mM NaCl, 1 mM EDTA pH 8.0, 0.1% [w/v] SDS, 0.1% [w/v] sodium deoxycholate, 1% [v/v] Triton X-100, protease inhibitors) to 200 μL. Input and gel samples were preserved. Lysates were incubated with 1μg/IP of anti-H2A.Z antibody (Diagenode, #C15410201) on a rotator overnight at 4 °C.

Dependent on the added amount of antibody, the amount of Protein A magnetic beads (ThermoFisher Scientific #10001D) was adjusted (e.g., for 1-2 μg of antibody/IP = 15 μL of beads) and blocked overnight with 0.1% [w/v] bovine serum albumin in RIPA buffer. On the following day, beads were added to the IP samples for 2 h on a rotator at 4 °C to capture the antibody-bound fragments. The immunoprecipitated chromatin was subsequently washed twice with 150 μL each of ice-cold buffers RIPA-LS, RIPA-HS (10 mM Tris-HCl pH 8.0, 50 0mM NaCl, 1 mM EDTA pH 8.0, 0.1% [w/v] SDS, 0.1% [v/v] sodium deoxycholate, 1% [v/v] Triton X-100), RIPA-LiCl (10 mM Tris-HCl pH 8.0, 250 mM LiCl, 1 mM EDTA pH 8.0, 0.5% [w/v] sodium deoxycholate, 0.5% [v/v] Nonidet P-40) and 10 mM Tris pH 8.0 containing protease inhibitors. Beads were washed once more with ice-cold 10 mM Tris pH 8.0 lacking inhibitors and transferred into new tubes.

Beads were resuspended in 25 μL of the tagmentation reaction mix (Nextera DNA Sample Prep Kit, Illumina) containing 5 μL of 5x Tagmentation buffer, 1 μL of Tagment DNA enzyme, topped up with H_2_O to the final volume and incubated at 37 °C for 10 minutes in a thermocycler. Beads were mixed after 5 minutes by gentle pipetting. To inactivate the Tn5 enzyme, 150 μL of ice-cold RIPA-LS was added to the tagmentation reaction. Beads were washed twice with 150 μL of RIPA-LS and 1x Tris-EDTA and subjected to de-crosslinking by adding 100 μL ChIPmentation elution buffer (160 mM NaCl, 40 μg/mL Rnase A (Sigma-Aldrich #R4642), 1x Tris-EDTA (Sigma #T9285) and incubating for 1h at 37 °C followed by overnight shaking at 65 °C. The next day, 4 mM EDTA and 200 μg/mL Proteinase K (Roche, #03115828001) were added, and samples incubated for another 2h at 45 °C with 1000 rpm shaking. Supernatant was transferred into a new tube and another 100 μL of ChIPmentation elution buffer was added for another hour at 45 °C with 1000 rpm shaking. DNA was isolated with MinElute PCR Purification Kit (Qiagen #28004) and eluted in 21 μL of H_2_O.

DNA for the final library was prepared with 25 μL NEBNext Ultra II Q5 Master Mix, 3.75 μL IDT custom primer i5_n_x (10 μM); 3.75 μL IDT custom primer i7_n_x (10 μM); 3.75 μL H_2_O and 13.75 μL ChIPmentation DNA. The Cq value obtained from the library quantification, rounded up to the nearest integer plus one additional cycle, was used to amplify the rest of the ChIPmentation DNA. Library qualities were verified by High Sensitivity DNA Analysis on the Bioanalyzer 2100 (Agilent) before performing sequencing on NextSeq 500 (paired-end 35bp reads) at the Core Unit Systemmedizin, Würzburg, Germany. All samples were sequenced at equimolar ratios.

### Read alignment

The read alignment pipeline was implemented and run in the workflow management system Watchdog (61, 62) as already previously described (38). Public sequencing data were downloaded from SRA using the sratoolkit version 2.10.8. Sequencing reads were aligned against the human genome (GRCh37/hg19), the HSV-1 genome (Human herpesvirus 1 strain 17, GenBank accession code: JN555585) and human rRNA sequences using ContextMap2 version 2.7.9 (63) (using BWA as short read aligner (64) and allowing a maximum indel size of 3 and at most 5 mismatches). For the two repeat regions in the HSV-1 genome, only one copy was retained each, excluding nucleotides 1–9,213 and 145,590–152,222 from the alignment. SAM output files of ContextMap2 were converted to BAM files using samtools (65). Read coverage in bedGraph format was calculated from BAM files using BEDTools (66). Subregions of promoter windows (TSS ± 3 kb) with differential read coverage in ATAC-seq and H2A.Z ChIPmentation/-seq data were determined with RegCFinder (36). For this purpose, RegCFinder was applied to ATAC-seq data for all pairwise comparisons of mock to WT and null mutant infections as well as mock to each timepoint of infection for the time-course data and to H2A.Z ChIPmentation and ChIP-seq data for the comparison of mock and WT infection as well as untreated and α-amanitin-treated HCT116 cells.

### Quality control

Statistics on numbers of mapped reads and reads mapped to human and HSV-1 genomes were determined with samtools (65). Promoter/Transcript body (PT) scores were determined with ATACseqQC (66). For peak calling, BAM files with mapped reads were converted to BED format using BEDvTools (67) and peaks were determined from these BED files using F-Seq with default parameters (68). The fraction of reads in peaks (FRiP) was calculated with featureCounts (69) using identified peaks as annotation. Annotation of peaks relative to genes was performed using ChIPseeker (70).

### Data plotting and statistical analysis

All figures were created in R and all statistical analyses were performed in R (67). Read coverage plots were created using the R Bioconductor package Gviz (68).

### Metagene and clustering analysis

Metagene analyses were performed as previously described (69) using the R program developed for this previous publication (available with the Watchdog *binGenome* module in the Watchdog module repository (https://github.com/watchdog-wms/watchdog-wms-modules/)). For promoter region analyses, the regions −3 kb to +3 kb of the TSS were divided into 101 equal-sized bins for each gene. For each bin, the average coverage per genome position was calculated and bin read coverages were then normalized by dividing by the total sum of all bins. Metagene curves for each replicate were created by averaging results for corresponding bins across all genes in a cluster/group and then averaged across replicates. For hierarchical clustering analysis, RegCFinder profiles of differential regions were calculated for each promoter window and comparison by setting each position in an m-RegC to 1, in an i-RegC to −1 and all other positions to 0. RegCFinder profiles of each promoter window for each comparison were concatenated into one row in the matrix. Hierarchical clustering of the resulting matrix was then performed using the hclust function in R according to Euclidean distances and Ward’s clustering criterion.

### Analysis of downstream open chromatin regions (dOCRs)

Open chromatin regions (OCRs) were determined from ATAC-seq data by first converting BAM files with mapped reads to BED format using BEDTools (70) and then determining enriched regions from these BED files using F-Seq with default parameters (71). dOCRs for individual genes were calculated from OCRs as previously described (21, 29). In brief, dOCRs are determined for genes by first assigning all OCRs overlapping with the 10 kb downstream of a gene to this gene. Second, OCRs starting at most 5 kb downstream of the so far most downstream OCR of a gene are also assigned to this gene. In both steps, individual OCRs can be assigned to multiple genes. The second step is iterated until no more OCRs can be assigned. The dOCR of a gene is then defined as the region from the gene 3’end to the end of the most downstream OCR assigned to this gene.

### Gene expression analysis

Number of fragments (=read pairs) per gene were determined from mapped paired-end RNA-seq reads in a strand-specific manner using featureCounts (72) and gene annotations from Ensembl (version 87 for GRCh37). For genes, all fragments overlapping exonic regions on the corresponding strand by ≥ 25bp were counted for the corresponding gene. Fold-changes in gene expression and statistical significance of changes were determined using DESeq2 (73) and p-values were adjusted for multiple testing using the method by Benjamini and Hochberg (74). Gene expression was quantified in terms of fragments per kilobase of exons per million mapped reads (FPKM). Only reads mapped to the human genome were counted for the total number of mapped reads for FPKM calculation.

### Motif discovery and enrichment analysis

Motif discovery was performed for each cluster separately for the i- and m-RegC regions using findMotifsGenome.pl script of the Homer suite (40), with our 7,649 input promoter windows as background. For this purpose, we used the hg19 annotation provided by Homer and automated trimming of input windows was disabled. Significant motifs were identified at a q-value (=False Discovery Rate (FDR) calculated with the Benjamini-Hochberg method (74)) cutoff of 0.01. Over-representation of Gene Ontology (GO) terms was performed separately for each cluster using the g:Profiler webserver (75) and the R package gprofiler2 (76), which provides an R interface to the webserver. As background gene list, the genes corresponding to our 7,649 input promoter windows were provided. P-values were corrected for multiple testing using the Benjamini-Hochberg method (74) and significantly over-represented GO terms were identified at an multiple testing adjusted p-value cutoff of 0.001. Enrichment of gene types (obtained from the Ensembl annotation (version 87 for GRCh37)) within clusters was determined using one-sided Fisher’s exact tests (with alternative = greater). Enrichment (odds-ratio >1) or depletion (odds-ratio < 1) of ICP4 binding sites from the study of Dremel *et al.* (41) within clusters as well as enrichment and depletion of clusters within gene groups with different expression levels were determined with two-sided Fisher’s exact tests. P-values were always corrected for multiple testing using the Benjamini-Hochberg method.

## Acknowledgements

This work was funded by the Deutsche Forschungsgemeinschaft (DFG, German Research Foundation, www.dfg.de) in the framework of the Research Unit FOR5200 DEEP-DV (443644894) project FR 2938/11-1 to C.C.F. and by the European Research Council (ERC, https://erc.europa.eu) project ERC-2021-CoG 101041177 – DecipherHSV to L.D.

## Notes

### Competing Interest Statement

The authors have declared no competing interest.

### Summary of Updates

We included additional controls, included additional relevant references and provided more details and figures on ATAC-seq coverage of viral genomes.

