## Supplemental Figures for "HSV-1 infection induces a downstream shift of the +1 nucleosome"

### Supplementary Figures

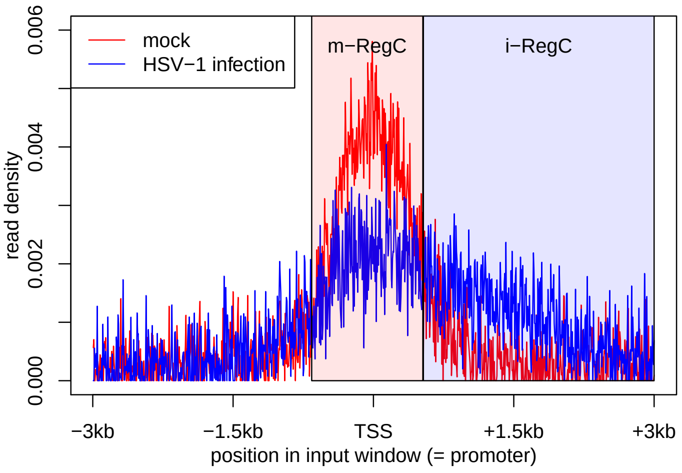


(a)

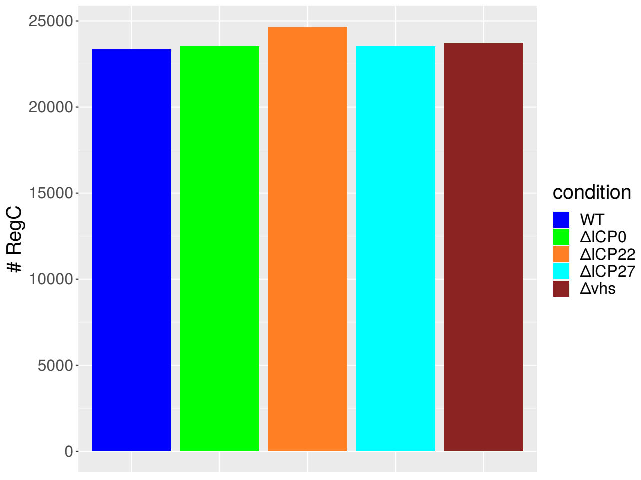


(b)

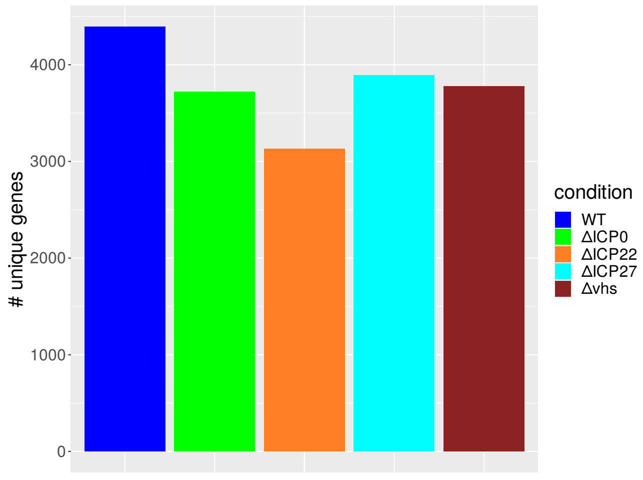


(c)

**(Continued on next page)**

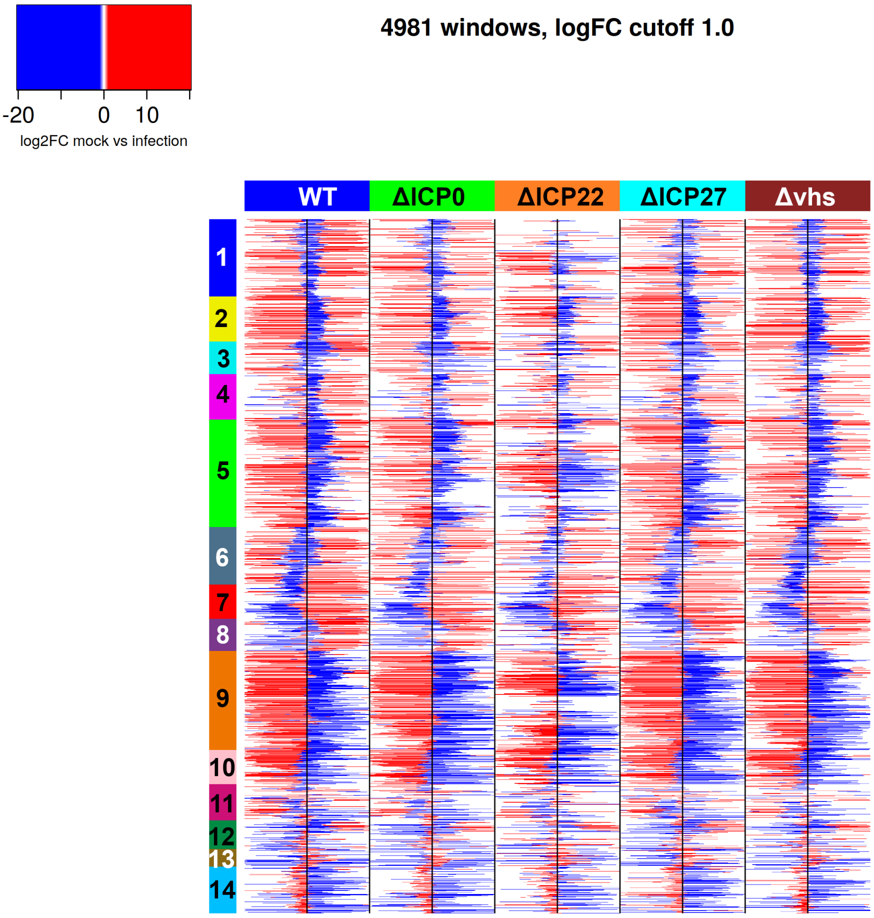


**(d)**

**Sup. Fig 1** **(a)** Overview on the RegCFinder approach (Weiss and Friedel, 2023). The key objective of RegCFinder is to identify subregions of an input window that show a relative increase in read density within the input window in one condition compared to a second condition. In our application, this means identifying subregions of 6 kb promoter windows that exhibit a relative increase in read density in mock compared to HSV-1 infection (red shaded region, denoted as m-RegC) or vice versa (blue shaded region, denoted as i-RegC). In this example, an m-RegC (mock>infection) is identified around the TSS and an i-RegC (infection>mock) downstream of the TSS. This would reflect a relative decrease of read density at the TSS and a relative increase downstream of the TSS during infection. Statistical significance of changes in read density between identified subregions of input windows (including identified m- and i-RegC as well as filler regions between them) is determined with DEXSeq (Anders *et al.*, 2012). **(b)** Barplot showing the number of identified differential regions (= m- and i-RegC) for WT and null mutant infections in comparison to mock infection. **(c)** Barplot showing the number of unique genes with at least one statistically significant (multiple testing adjusted p-value (adj. p.) $\leq 0.01$) differential region for WT and null mutant infection in comparison to mock infection. **(d)** Heatmap visualizing log2 fold-changes for the identified differential regions shown in **Fig. 1a**. For this purpose, results for the same input window for the different comparisons of WT or null mutant virus infections to mock are concatenated in one row of the heatmap matrix. Colored rectangles on top of columns indicate which virus infection was compared to mock. Black vertical lines in the center of each comparison indicate the position of the TSS. Regions corresponding to statistically significant differential regions are colored according to the log2 fold-change in mock vs. infection determined by DEXSeq. Here, the color scale is continuous between -1 and 1 and log2 fold-changes $> 1$ are colored the same red and log2 fold-changes $< 1$ the same blue. Promoter windows are ordered as in **Fig. 1a** and clusters from **Fig. 1a** are annotated as colored and numbered rectangles on the left.

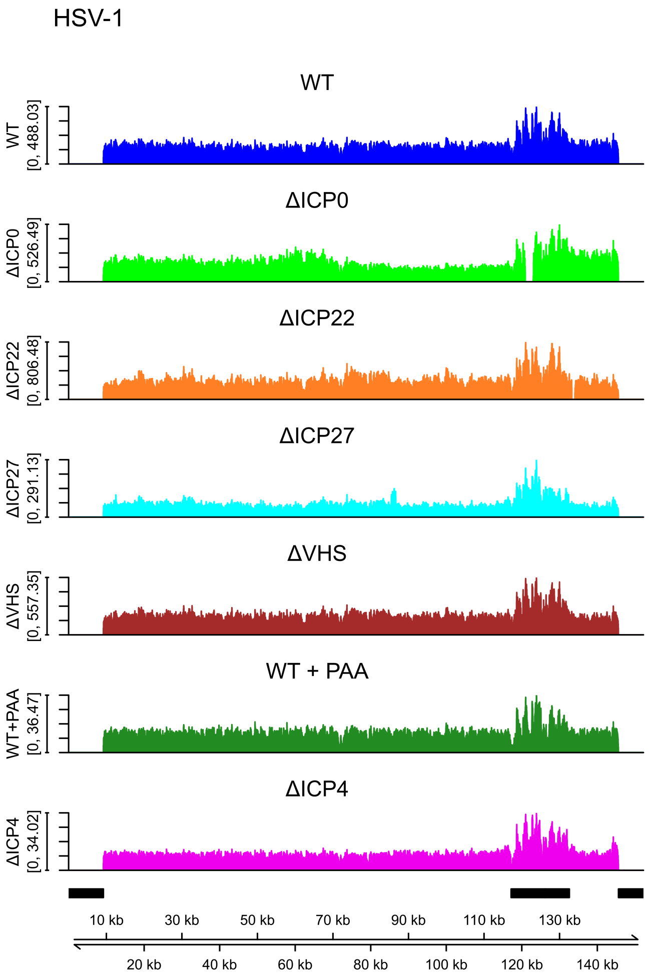


(a)

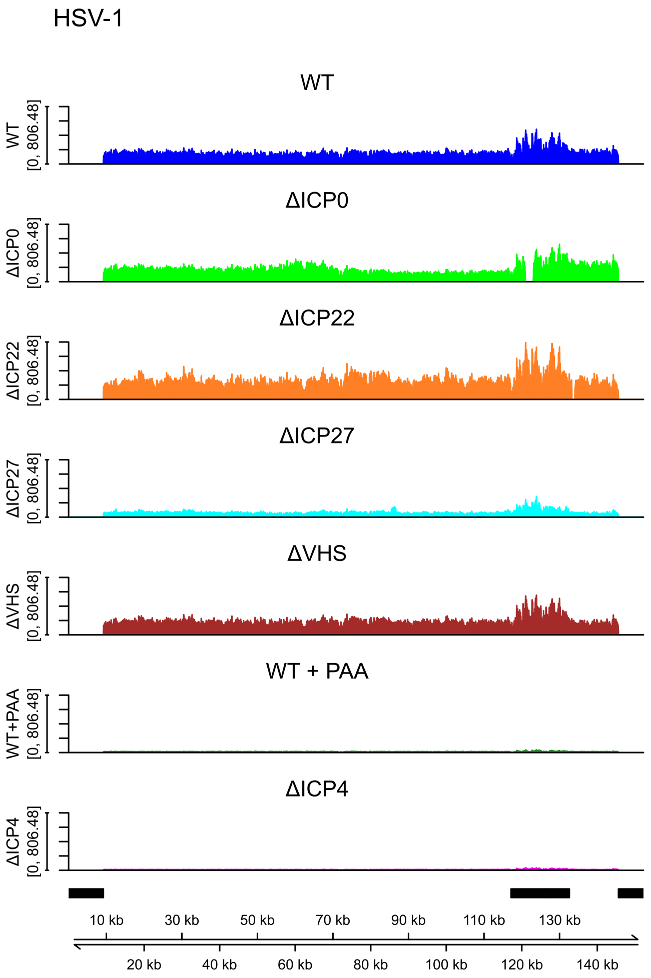


(b)

(Continued on next page)

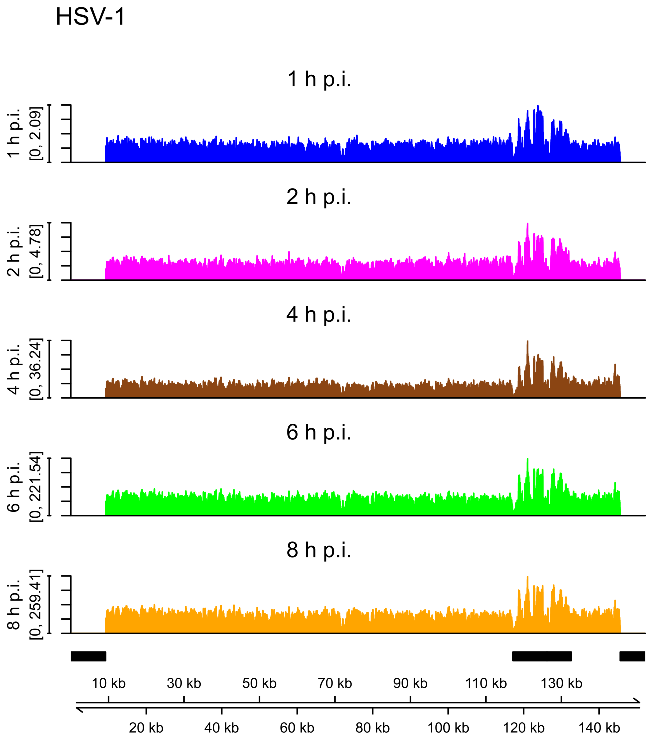


**(c)**

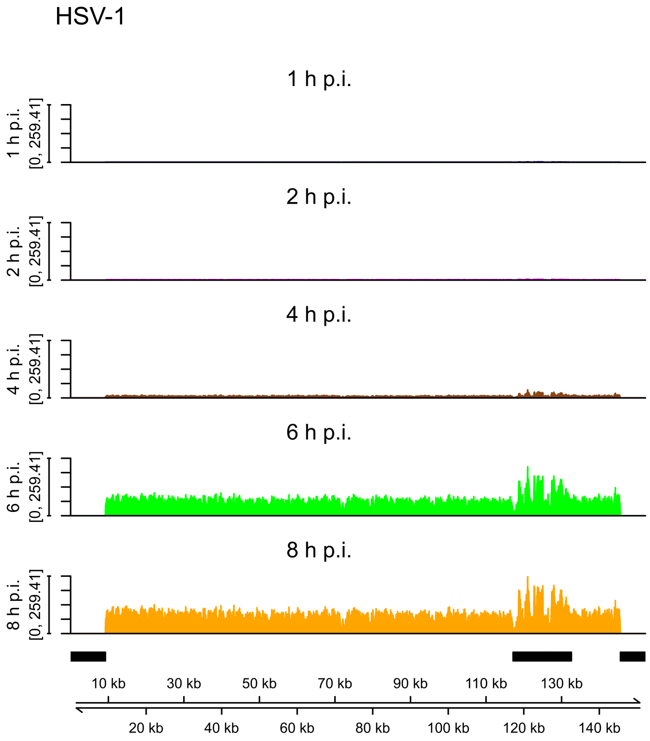


**(d)**

**Sup. Fig 2** Read coverage on the HSV-1 genome for **(a,b)** WT (blue), ΔICP0 (green), ΔICP22 (orange), ΔICP27 (cyan), Δ*vhs* (brown), WT+PAA infection (dark green) and ΔICP4 infection (magenta) from the first ATAC-seq experiment and **(c,d)** 1 h (blue), 2 h (magenta), 4 h (brown), 6 h (green), and 8 h p.i. (orange) HSV-1 infection from the ATAC-seq time-course experiment. Read coverage was normalized to total number of mapped reads for each sample and averaged between replicates. For **(a)** and **(c)**, the y-axis range was determined independently for each condition and for **(b)** and **(d)** the same y-range was chosen for all conditions based on the maximum observed coverage. The black rectangles above the genome coordinates depict the position of the inverted repeat regions in the HSV-1 genome. The terminal repeat copies at the start and end of the genome were masked from read alignment, resulting in all reads from the inverted repeats mapping to the internal repeat copies.

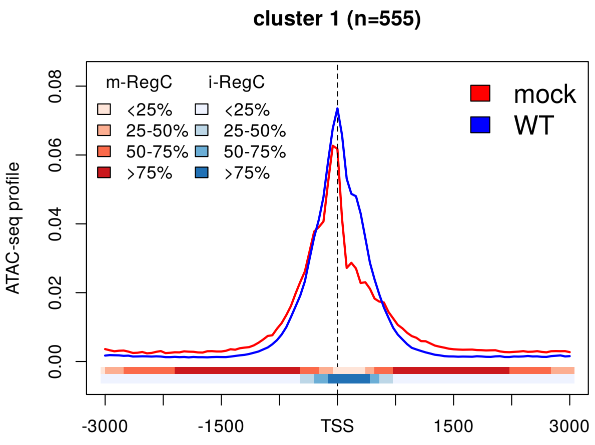


(a) pattern I

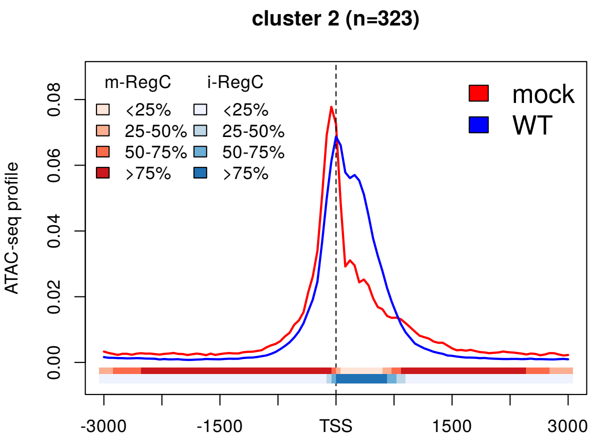


(b) pattern I

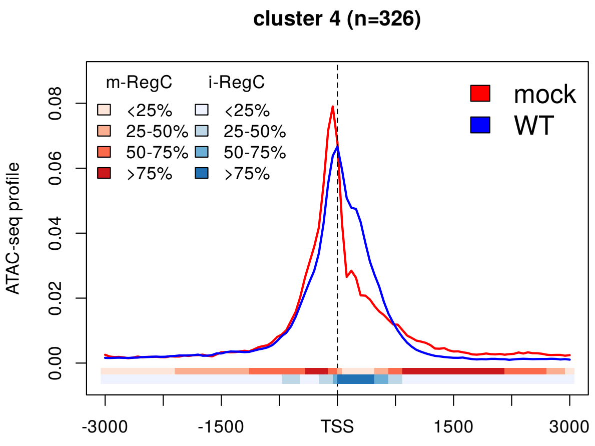


(c) pattern I

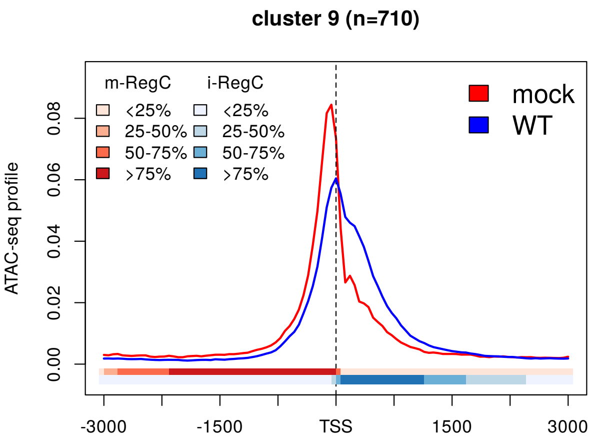


(d) pattern I

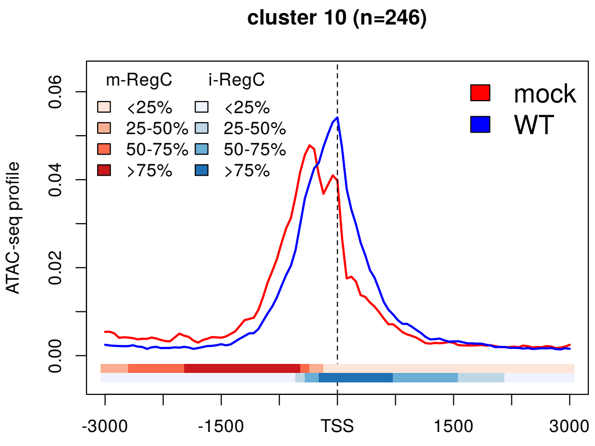


(e) pattern I

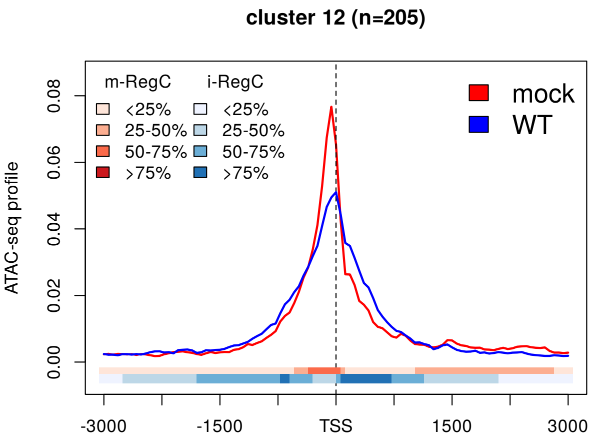


(f) pattern I

(Continued on next page)

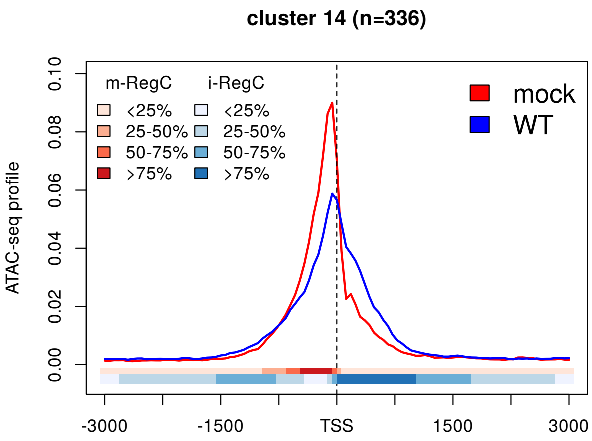


(g) pattern I

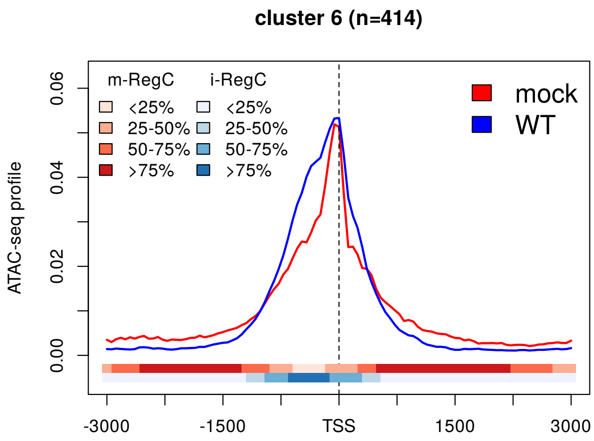


(h) pattern II

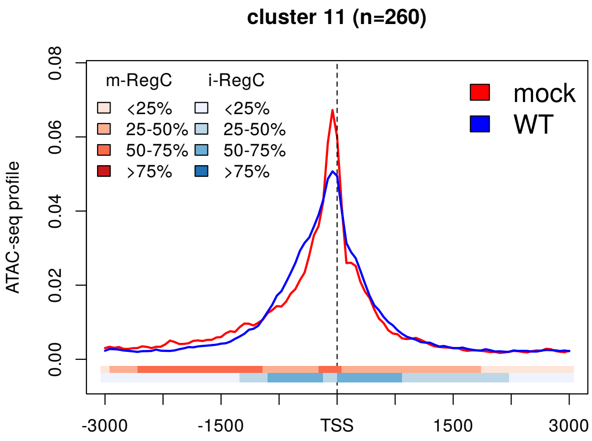


(i) pattern II

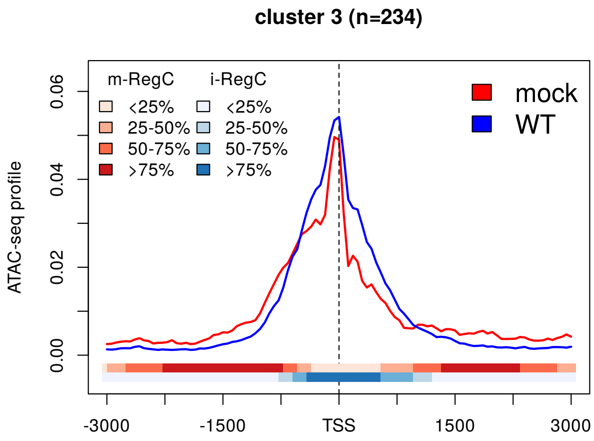


(j) pattern I + II

**Sup. Fig 3** Metagene plots showing ATAC-seq profiles around the TSS $\pm$ 3 kb in mock (red) and WT (blue) infection for all clusters apart from clusters 5, 7, 8, and 13, which are shown in **Fig. 1**. See Materials and Methods for a detailed description of metagene plots. The colored bands below the metagene curves in each subfigure indicate the percentage of genes having an m- or i-RegC (red or blue, respectively) at that position for the comparison of WT infection to mock. Subfigures **(a-g)** show clusters with pattern I with a shift and/or broadening of the TSS peak into downstream regions, while **(h-i)** show clusters that exhibit the mirror pattern II with a shift and/or broadening of the TSS peak into upstream regions. Subfigure **(j)** shows a cluster with combined patterns I and II.

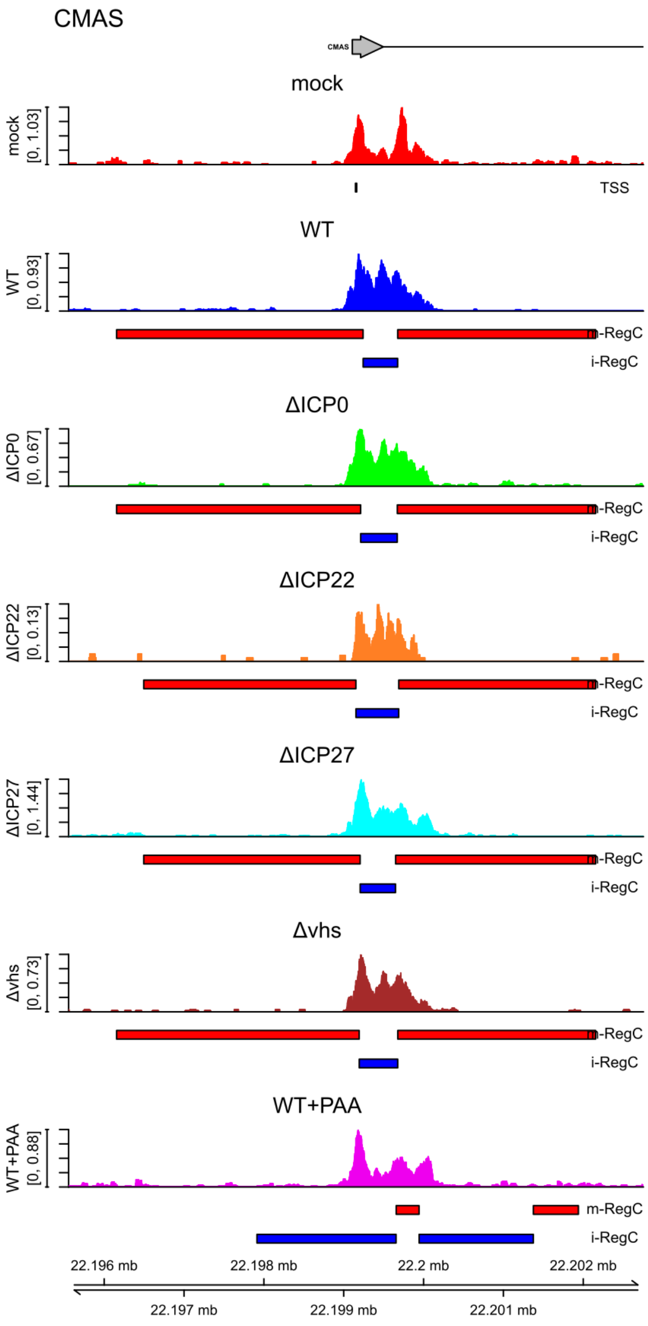


**(a)** pattern I, cluster 1

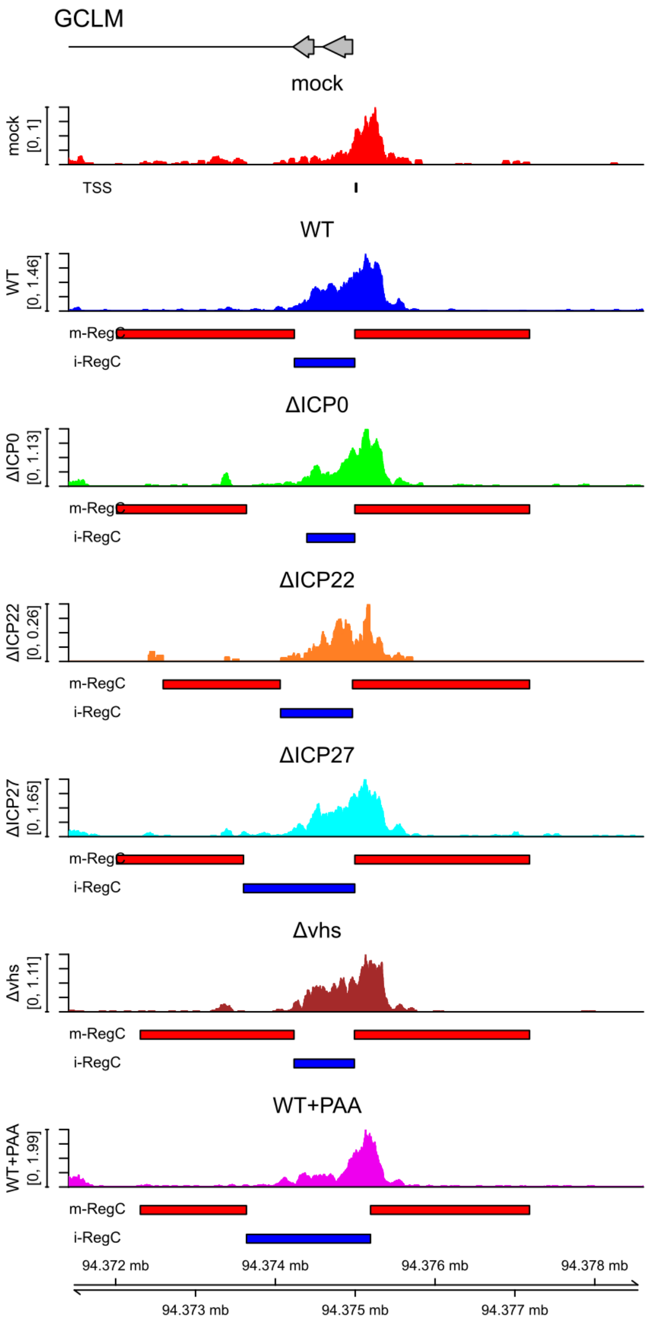


**(b)** pattern I, cluster 2

**(Continued on next page)**

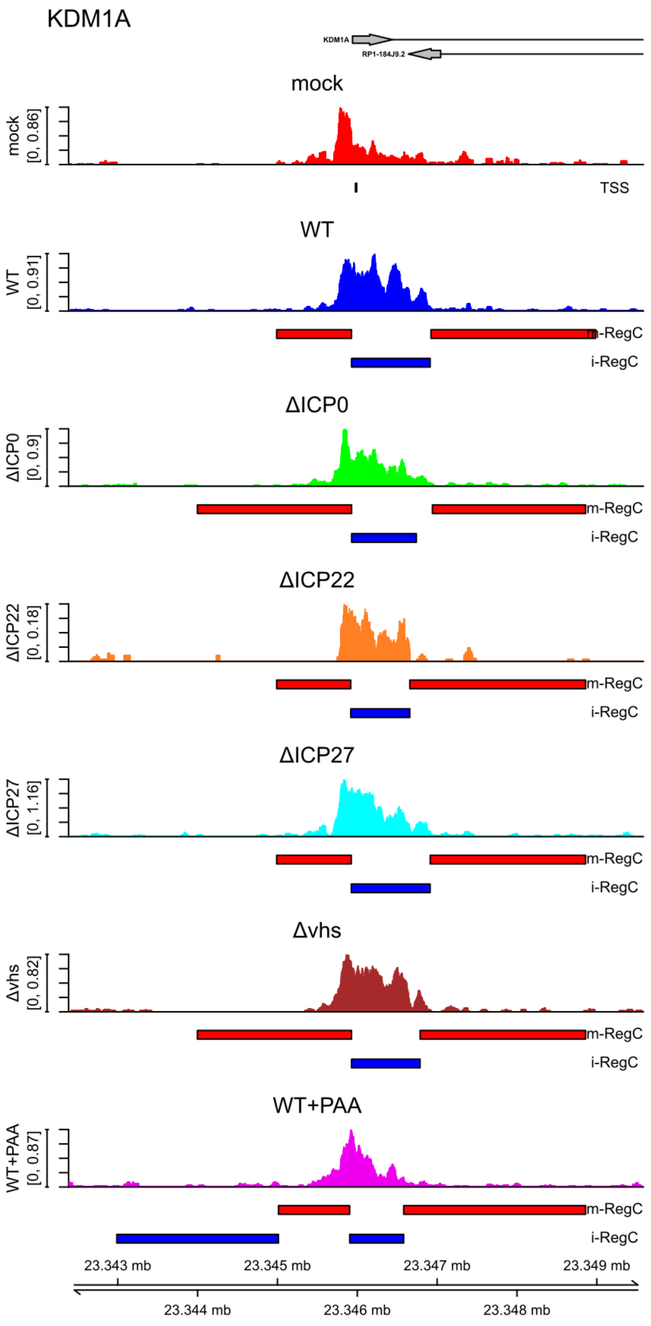


**(c)** pattern I, cluster 4

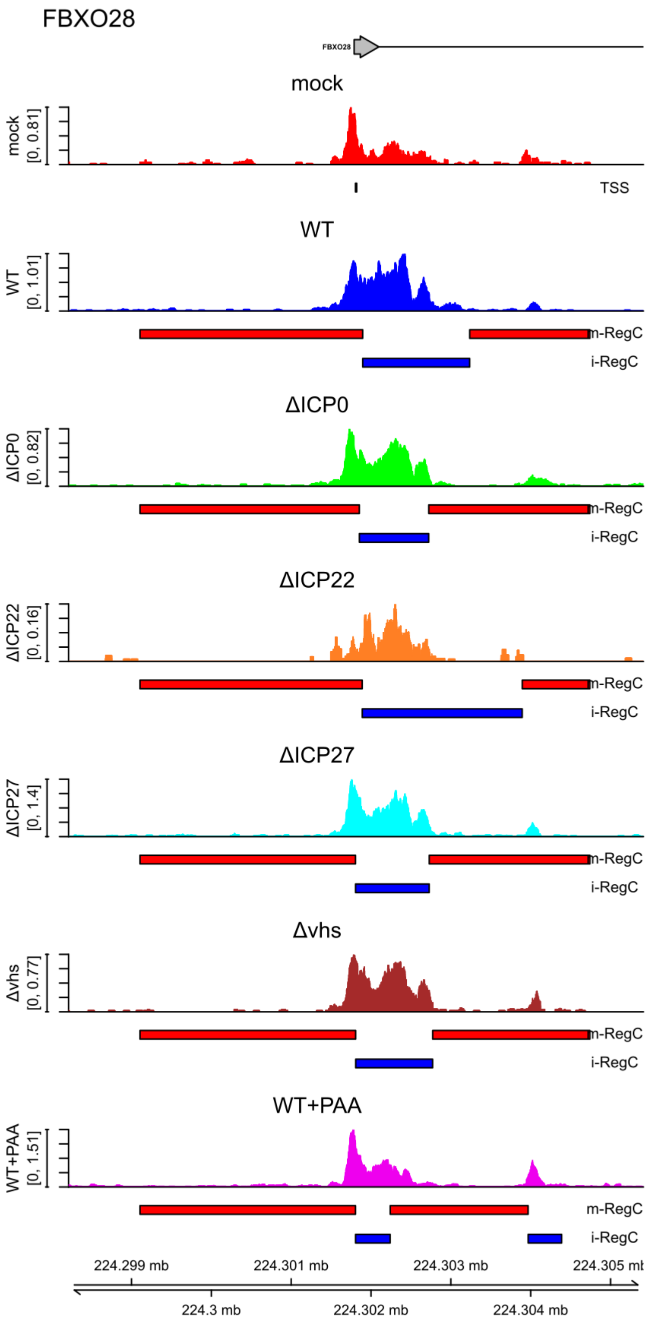


**(d)** pattern I, cluster 5

**(Continued on next page)**

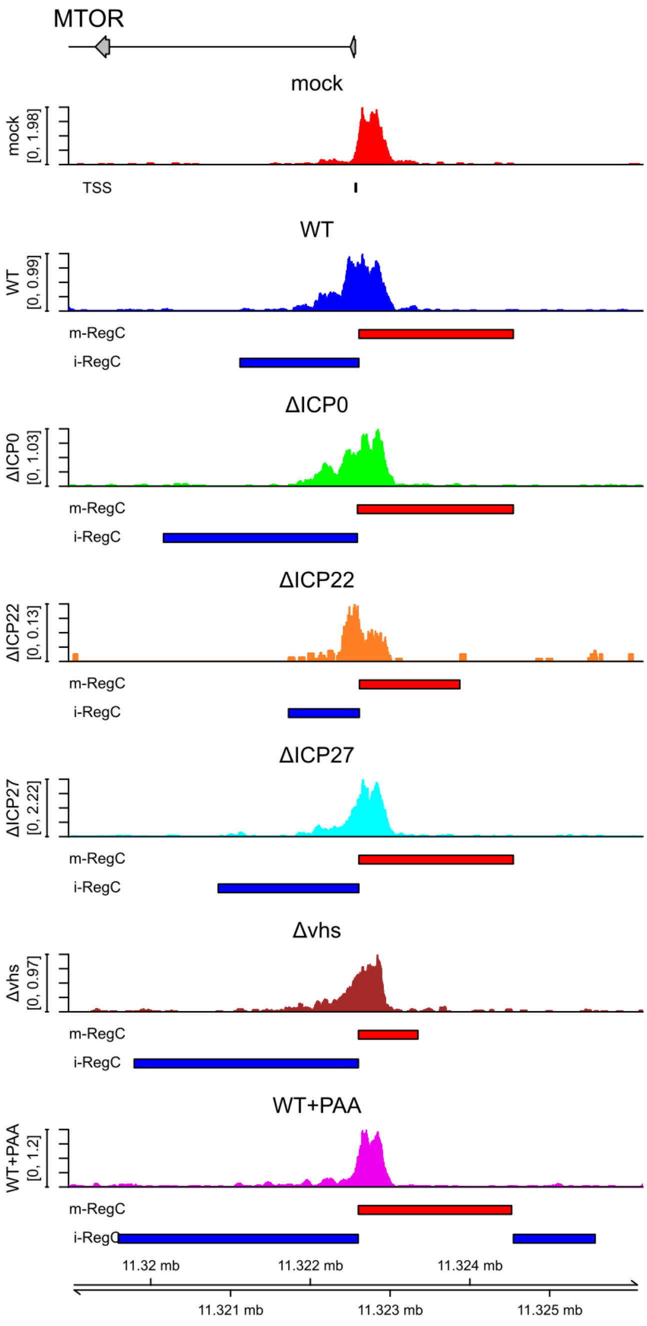


(e) pattern I, cluster 9

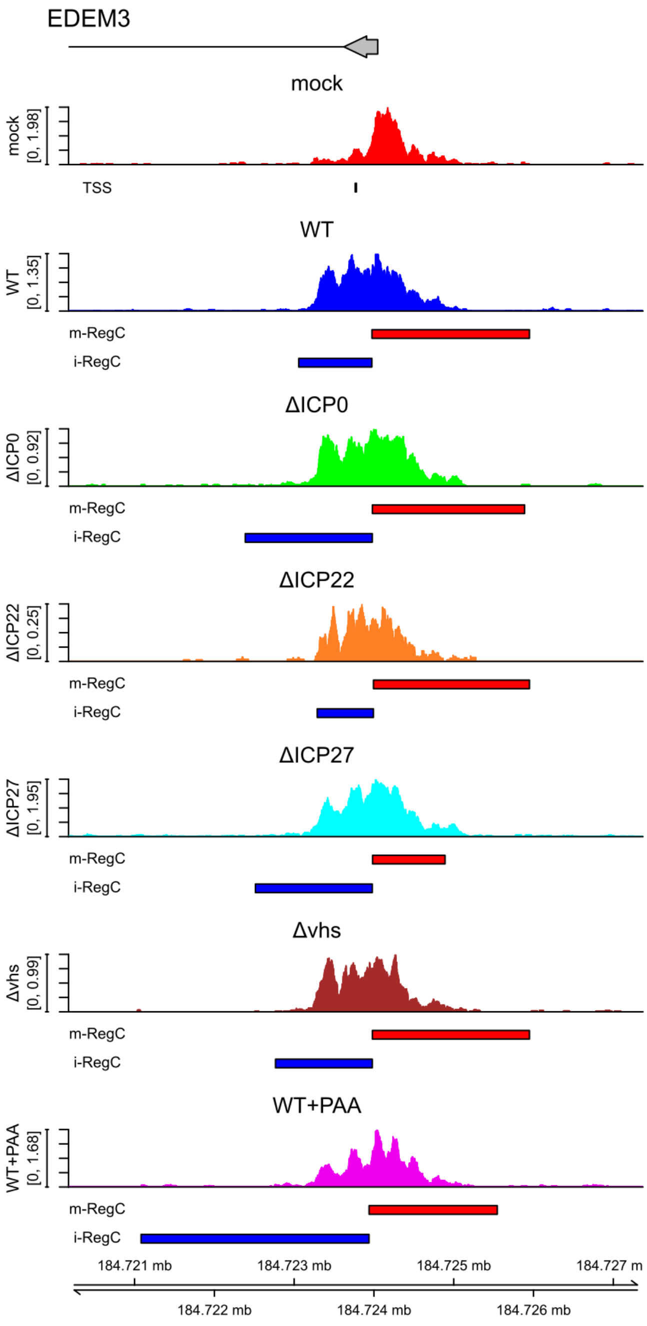


(f) pattern I, cluster 10

**(Continued on next page)**

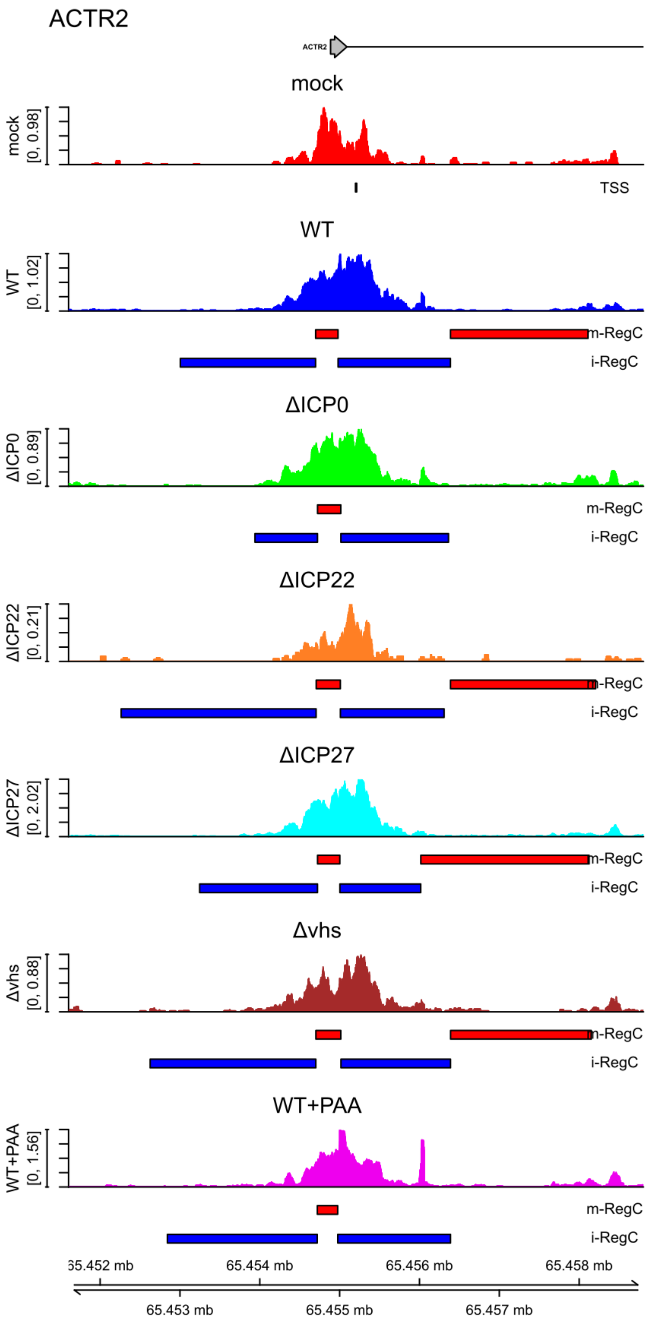


**(g)** pattern I, cluster 12

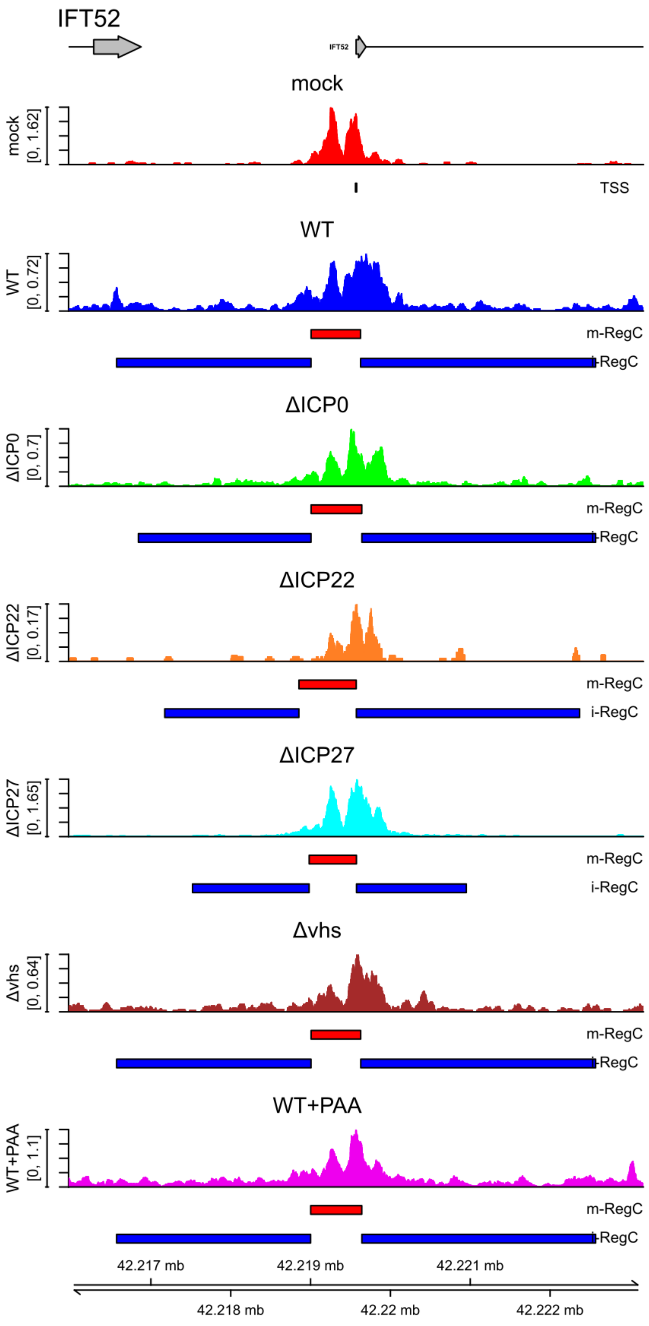


**(h)** pattern I, cluster 14

**(Continued on next page)**

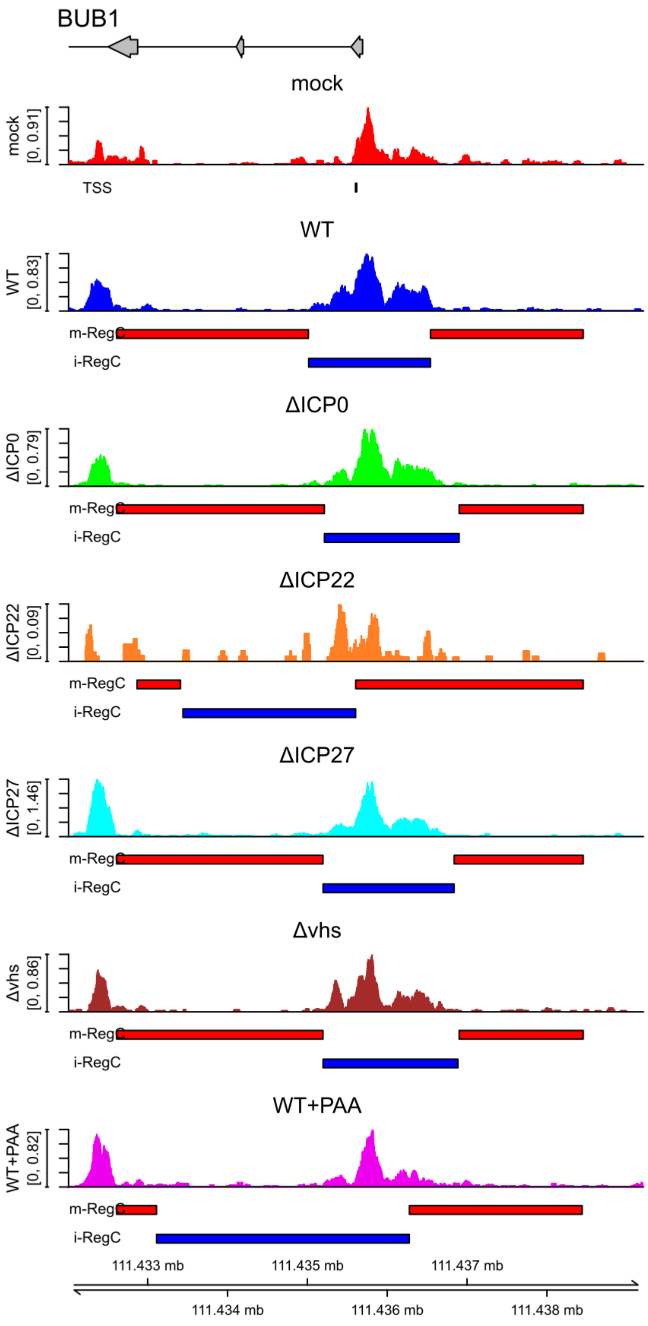


(i) pattern II, cluster 6

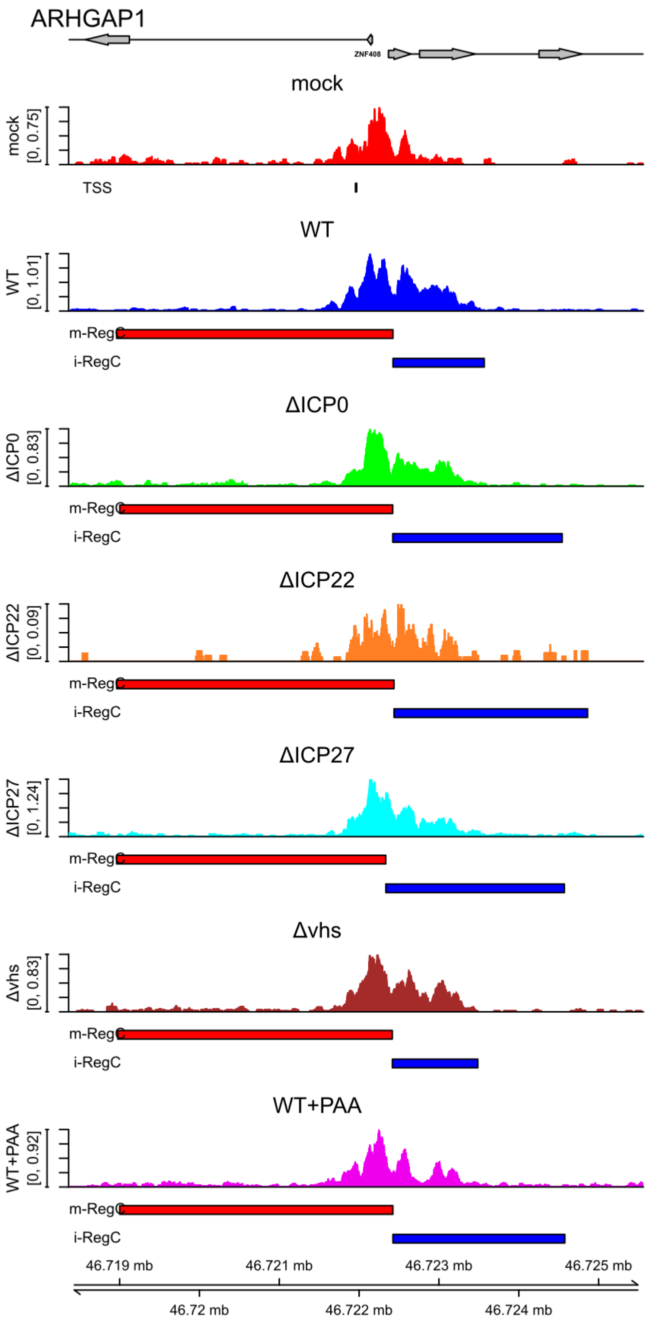


(j) pattern II, cluster 7

**(Continued on next page)**

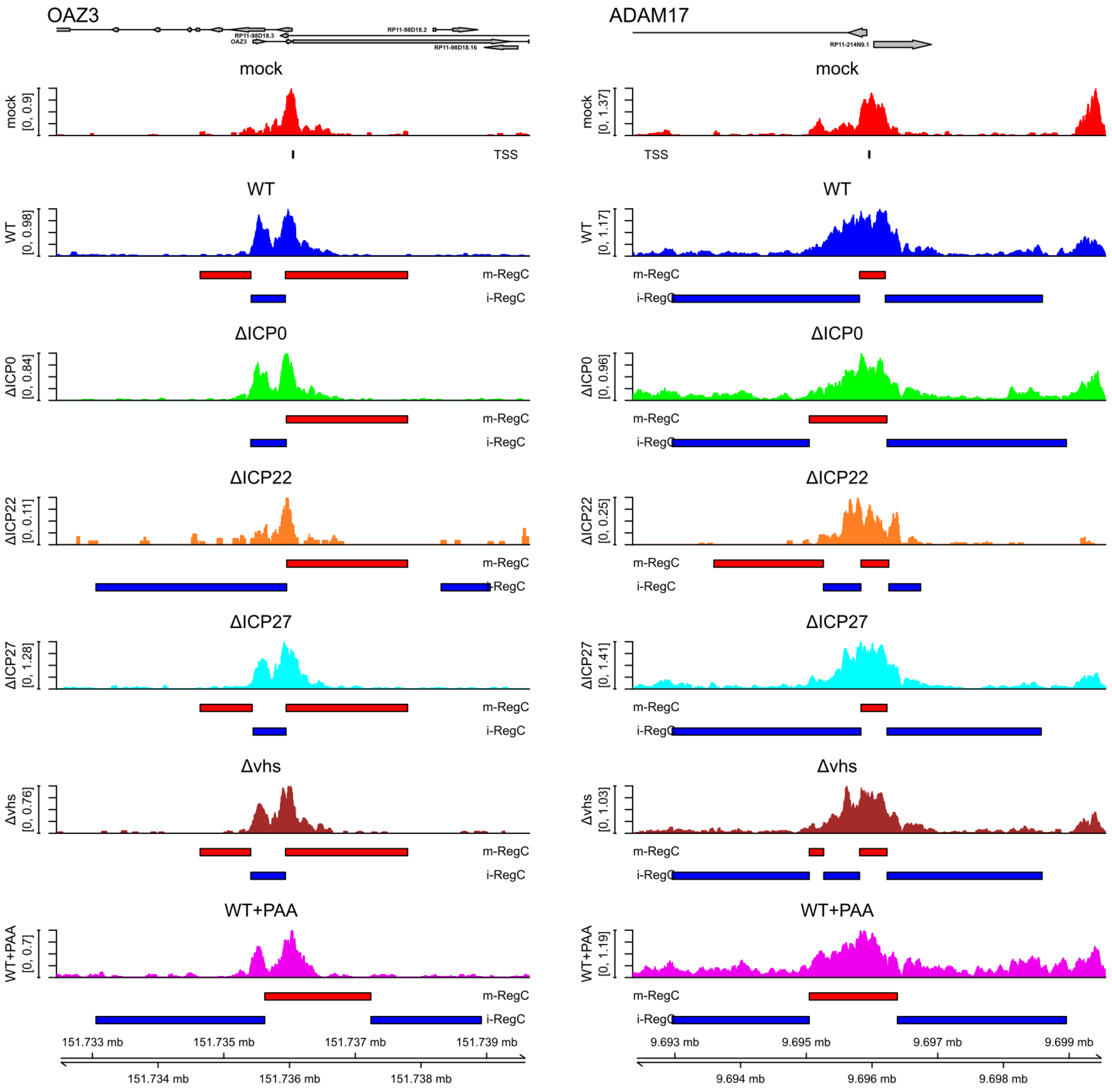


**(k)** pattern II, cluster 11

**(l)** pattern III, cluster 13

**(Continued on next page)**

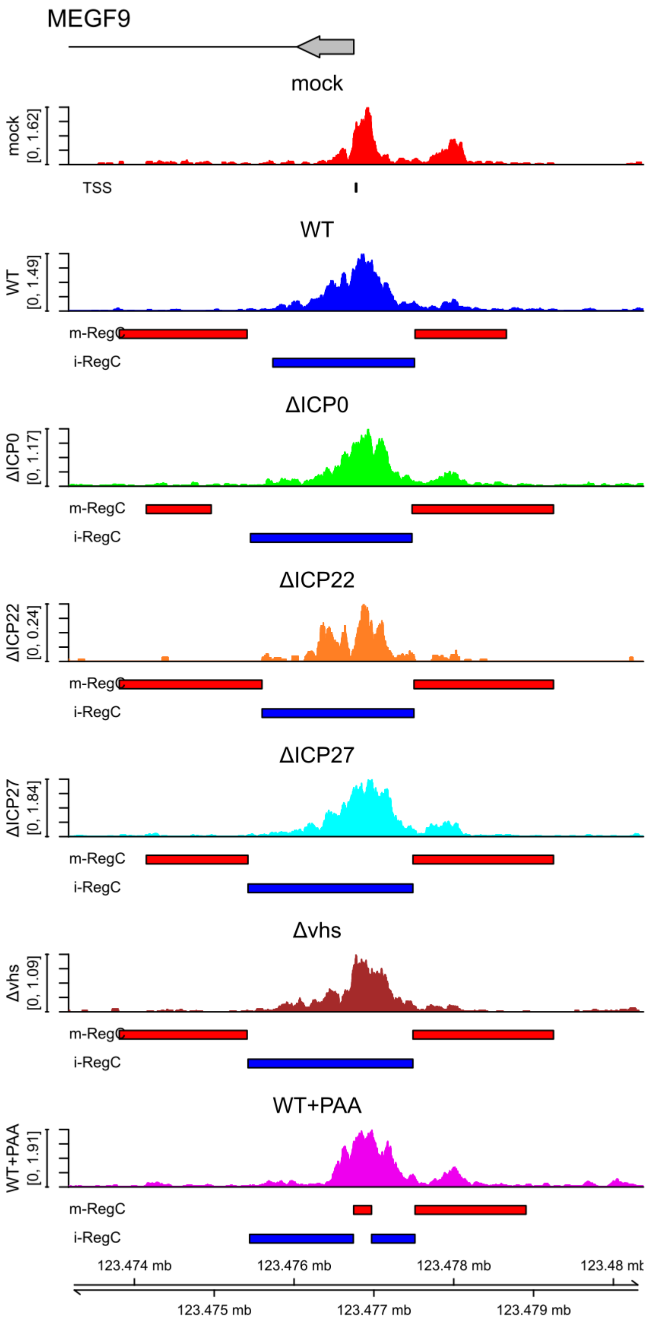


**(m)** pattern I + II, cluster 3

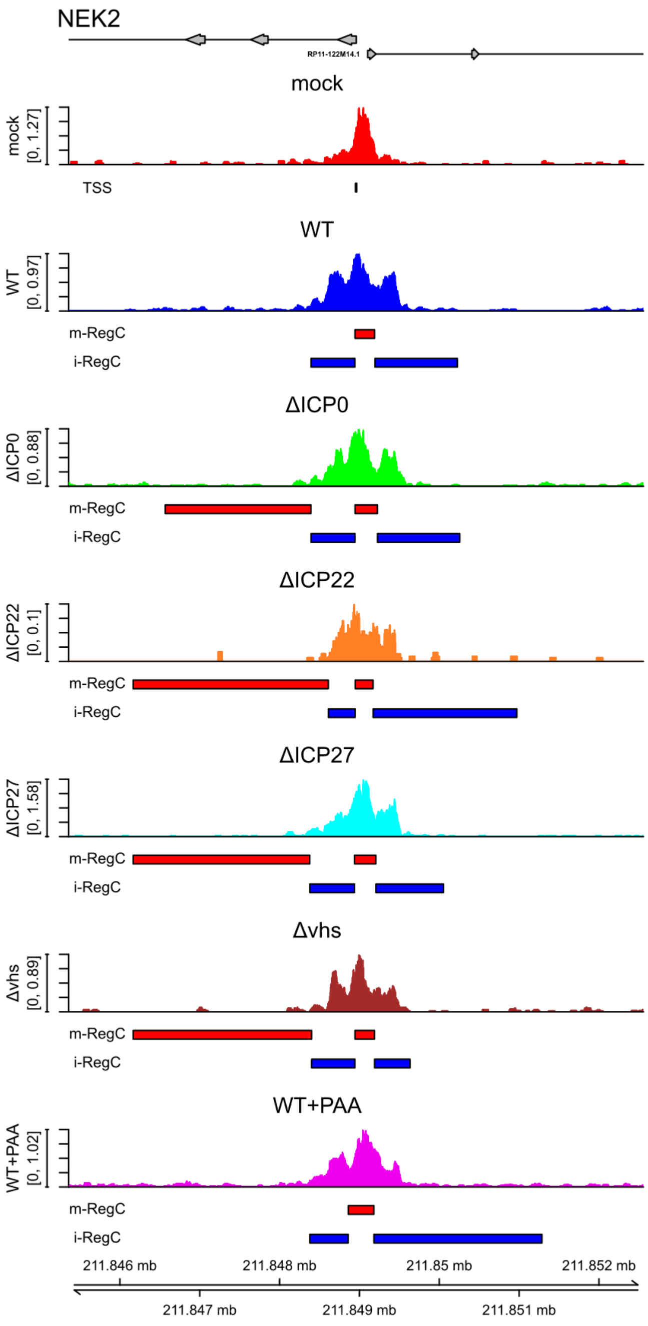


**(n)** pattern I + II, cluster 8

**Sup. Fig 4** Read coverage in a ± 3.6 kb window around the TSS in ATAC-seq data of mock (red), WT (blue), ΔICP0 (green), ΔICP22 (orange), ΔICP27 (cyan), Δ*vhs* (brown) and WT+PAA infection (magenta) for example genes for all patterns and clusters. Pattern and cluster are indicated below each subfigure. Read coverage was normalized to total number of mapped reads for each sample and averaged between replicates. The TSS used in the analysis is indicated by a short vertical line below the read coverage track for mock infection. Gene annotation is indicated at the top. Boxes represent exons, lines represent introns, and direction of transcription is indicated by arrowheads. The name of the gene whose promoter window was analyzed is indicated in larger font on the top left and – if not clear from the context – beside the gene annotation. Names for other genes overlapping the input window are indicated next to these genes if necessary. Below each read coverage track m- (red bars) and i-RegCs (blue bars) are indicated for the comparison of the corresponding virus infection to mock.

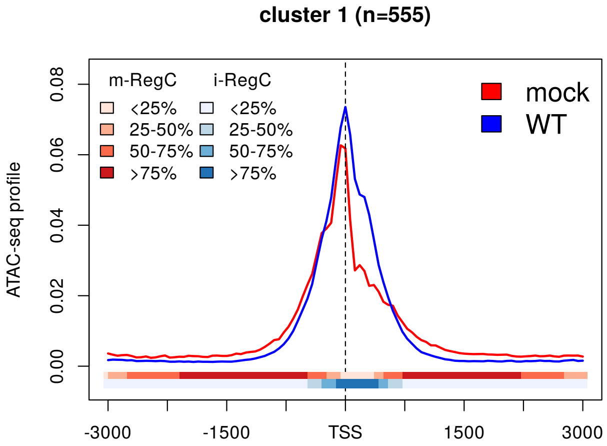


(a)

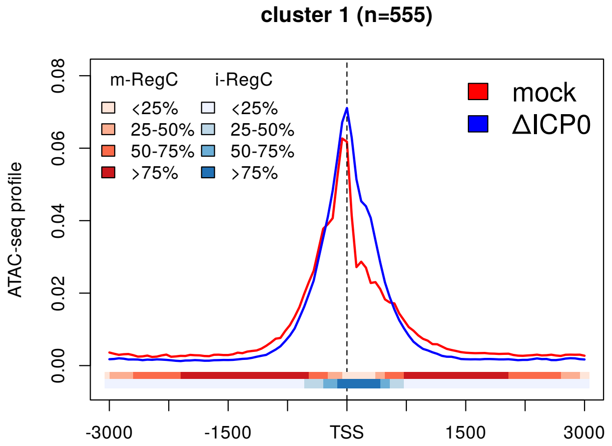


(b)

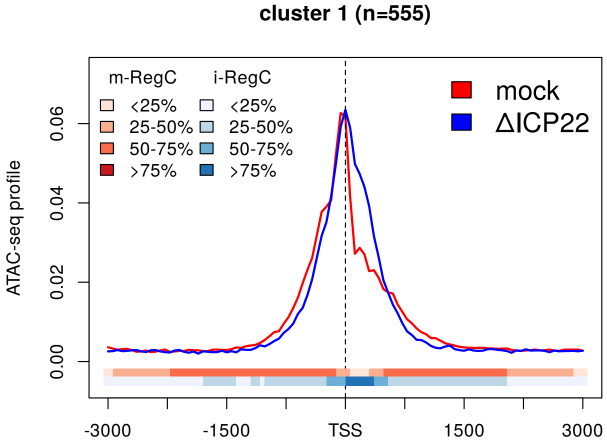


(c)

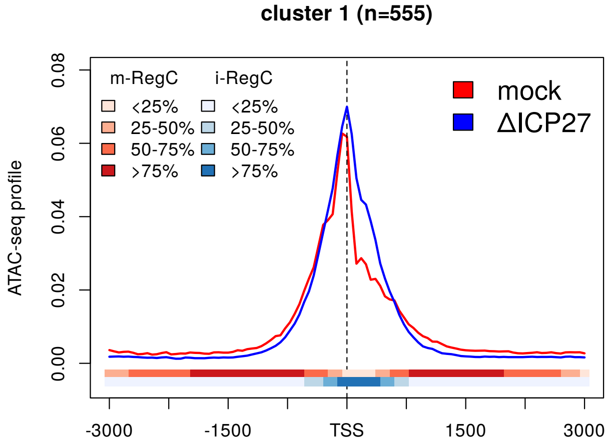


(d)

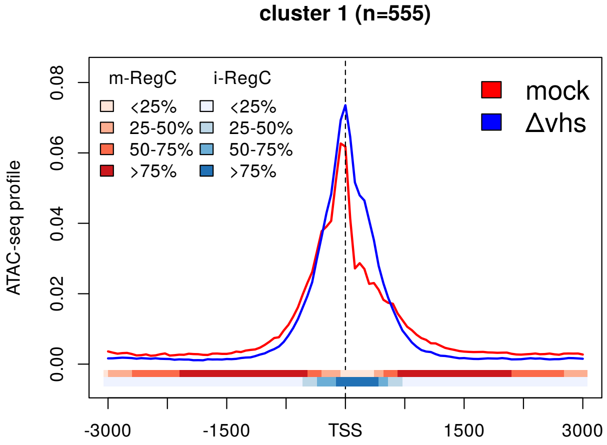


(e)

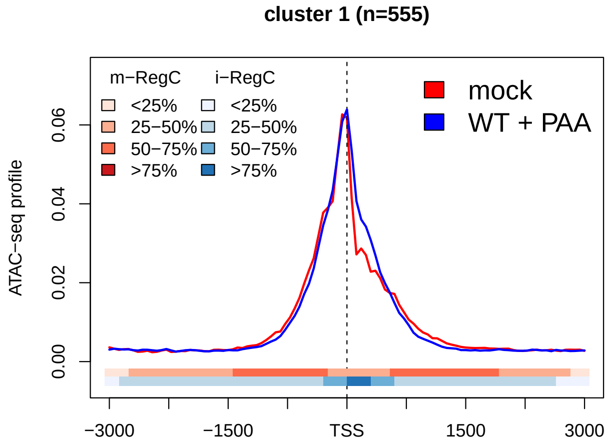


(f)

**Sup. Fig 5** Metagene plots of ATAC-seq profiles in mock as well as **(a)** WT infection, **(b-e)** null mutant infections and **(f)** WT+PAA infection for cluster 1 (pattern I). See Materials and Methods for a detailed description of metagene plots. The colored bands below the metagene curves in each subfigure indicate the percentage of genes having an m- or i-RegC (red or blue, respectively) at that position for the corresponding comparison of WT or null mutant virus infection to mock.

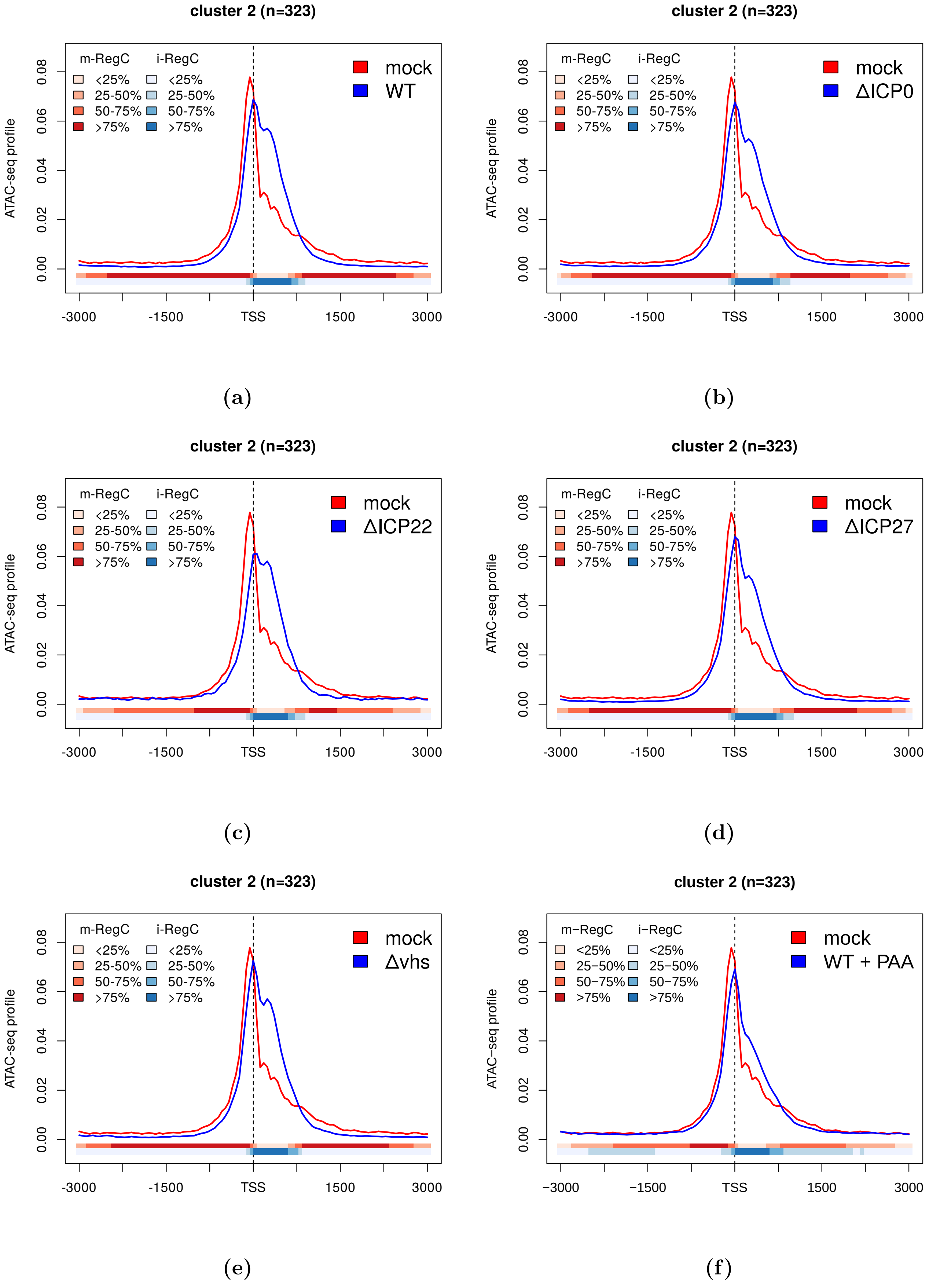


**Sup. Fig 6** Metagene plots of ATAC-seq profiles in mock as well as **(a)** WT infection, **(b-e)** null mutant infections and **(f)** WT+PAA infection for cluster 2 (pattern I). See Materials and Methods for a detailed description of metagene plots. The colored bands below the metagene curves in each subfigure indicate the percentage of genes having an m- or i-RegC (red or blue, respectively) at that position for the corresponding comparison of WT or null mutant virus infection to mock.

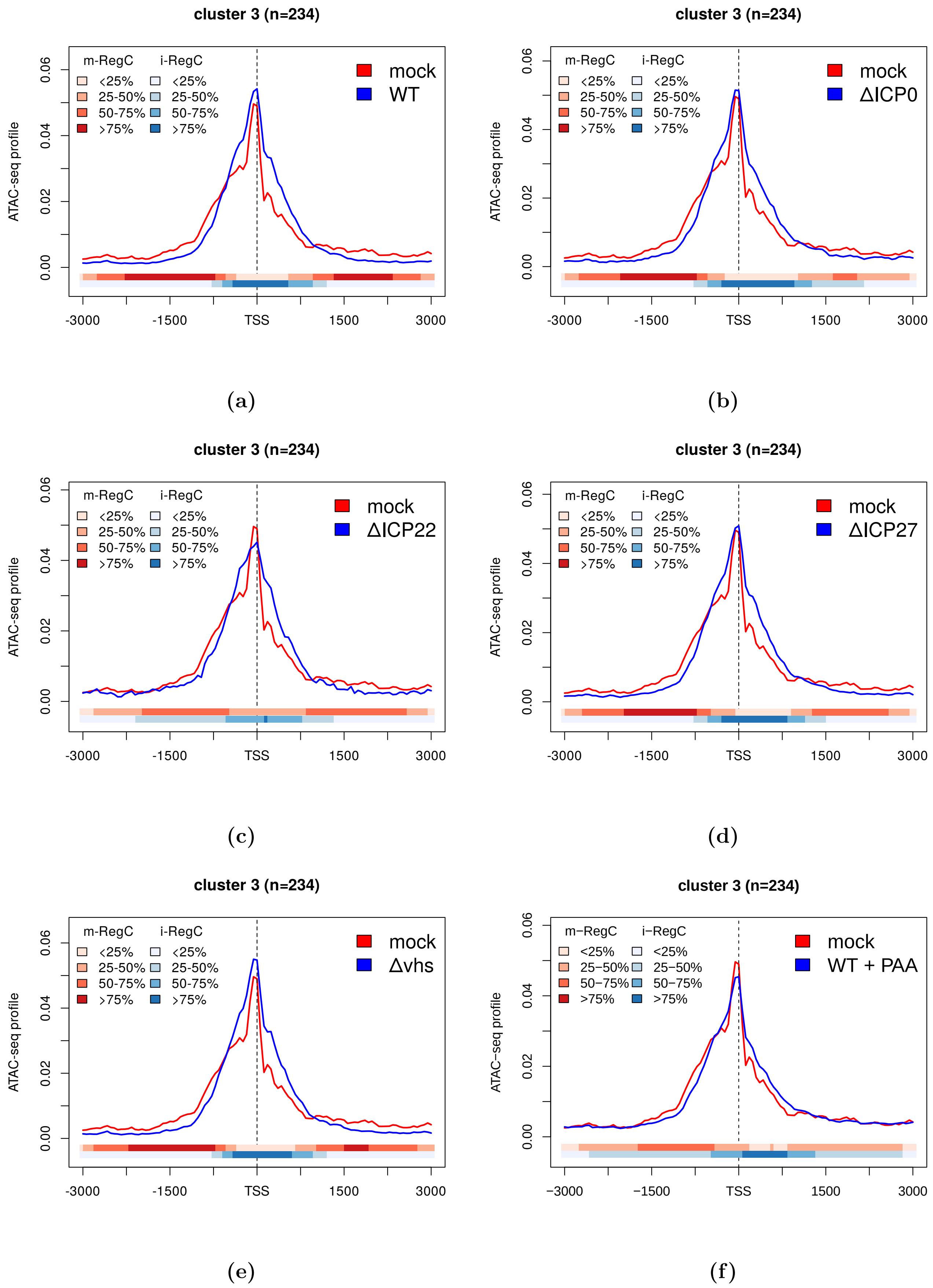


**Sup. Fig 7** Metagene plots of ATAC-seq profiles in mock as well as **(a)** WT infection, **(b-e)** null mutant infections and **(f)** WT+PAA infection for cluster 3 (combined pattern I + II). See Materials and Methods for a detailed description of metagene plots. The colored bands below the metagene curves in each subfigure indicate the percentage of genes having an m- or i-RegC (red or blue, respectively) at that position for the corresponding comparison of WT or null mutant virus infection to mock.

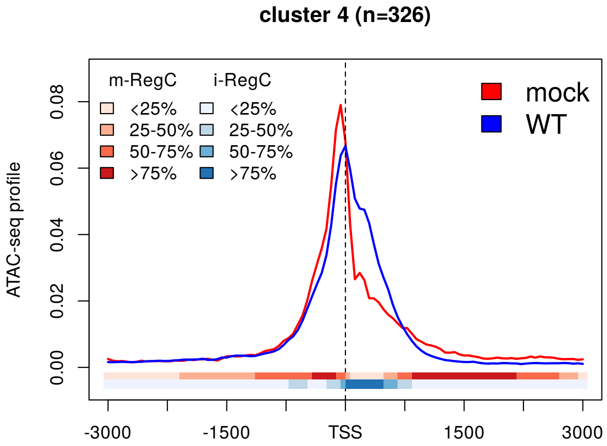


(a)

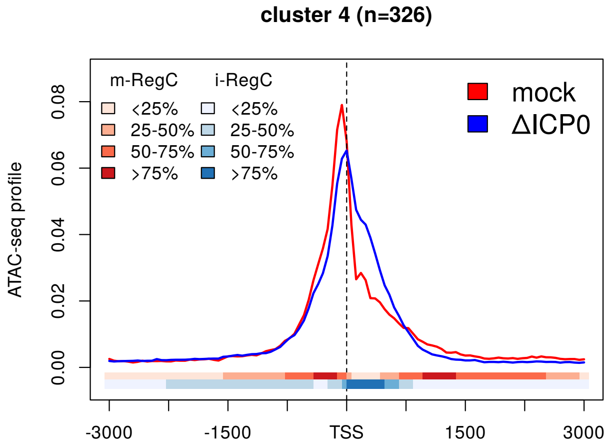


(b)

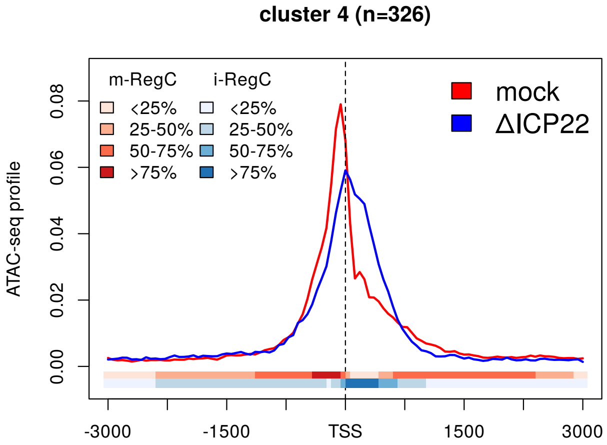


(c)

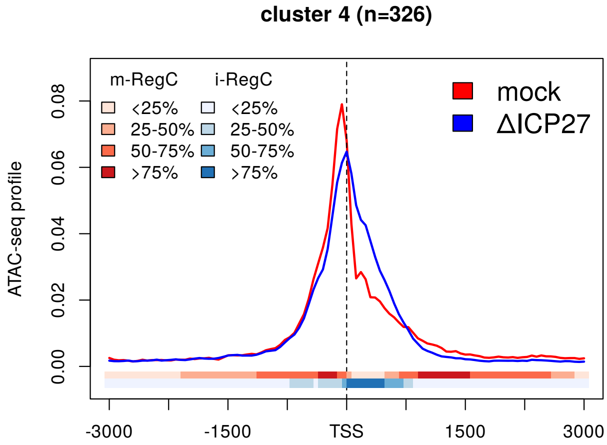


(d)

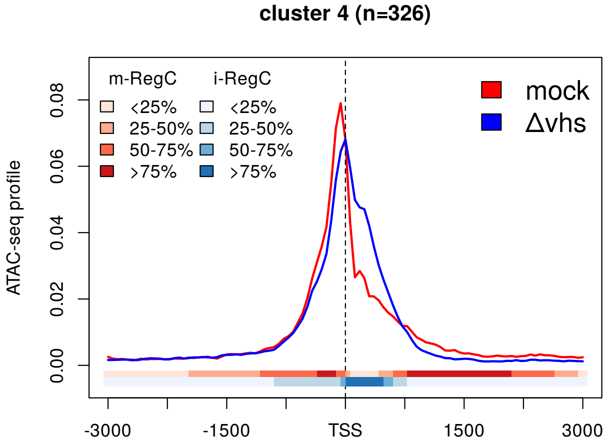


(e)

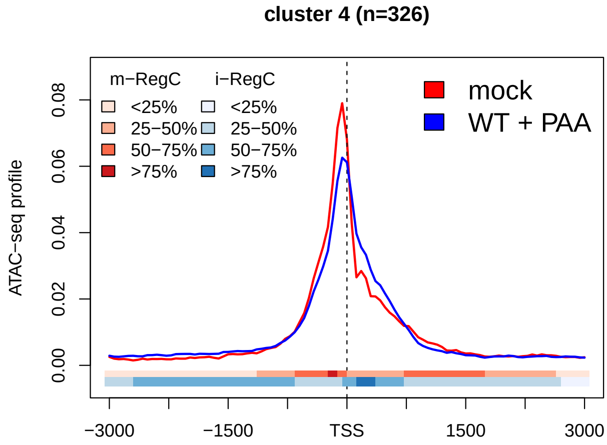


(f)

**Sup. Fig 8** Metagene plots of ATAC-seq profiles in mock as well as **(a)** WT infection, **(b-e)** null mutant infections and **(f)** WT+PAA infection for cluster 4 (pattern I). See Materials and Methods for a detailed description of metagene plots. The colored bands below the metagene curves in each subfigure indicate the percentage of genes having an m- or i-RegC (red or blue, respectively) at that position for the corresponding comparison of WT or null mutant virus infection to mock.

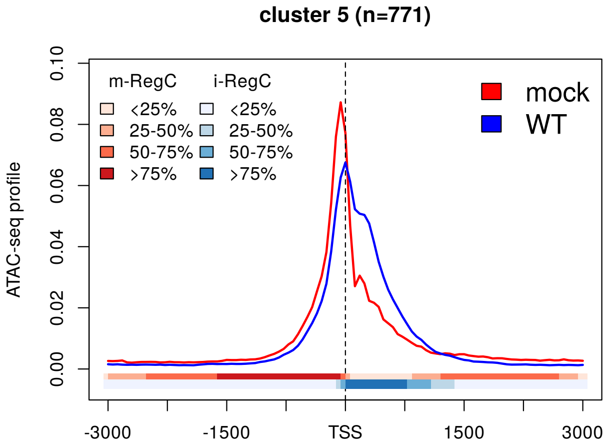


(a)

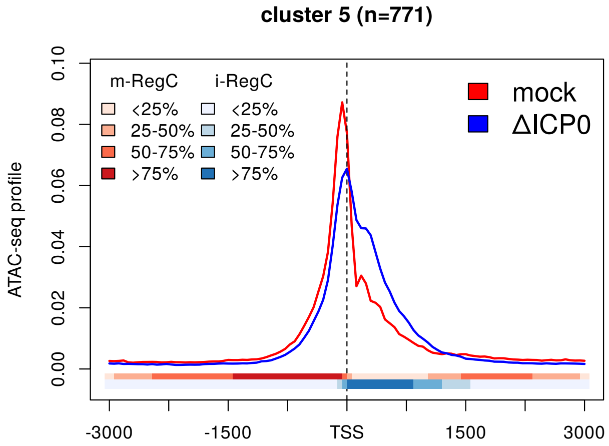


(b)

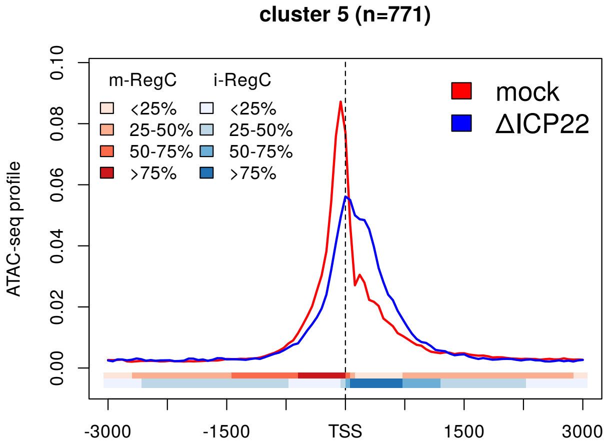


(c)

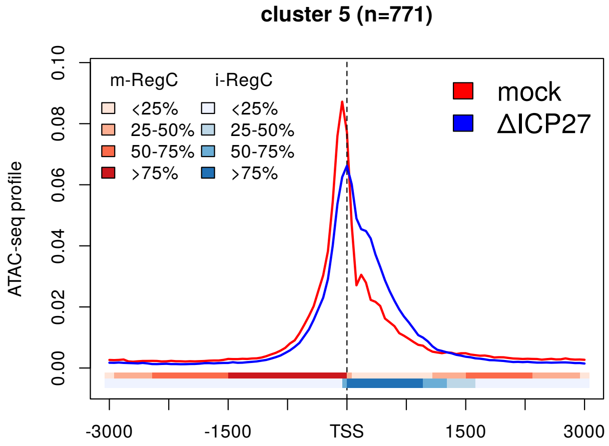


(d)

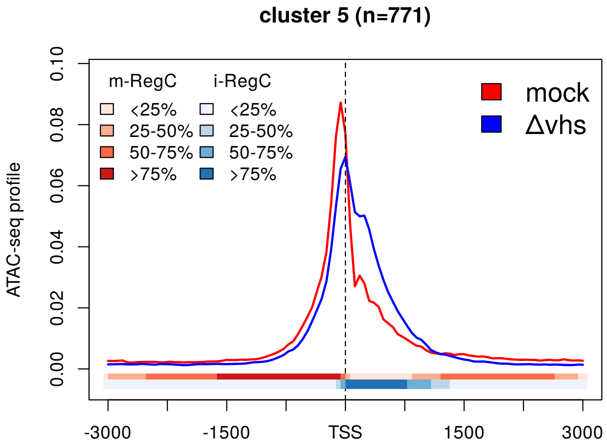


(e)

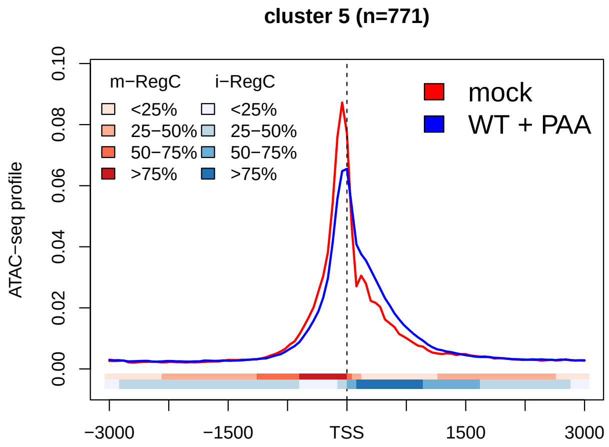


(f)

**Sup. Fig 9** Metagene plots of ATAC-seq profiles in mock as well as **(a)** WT infection, **(b-e)** null mutant infections and **(f)** WT+PAA infection for cluster 5 (pattern I). See Materials and Methods for a detailed description of metagene plots. The colored bands below the metagene curves in each subfigure indicate the percentage of genes having an m- or i-RegC (red or blue, respectively) at that position for the corresponding comparison of WT or null mutant virus infection to mock.

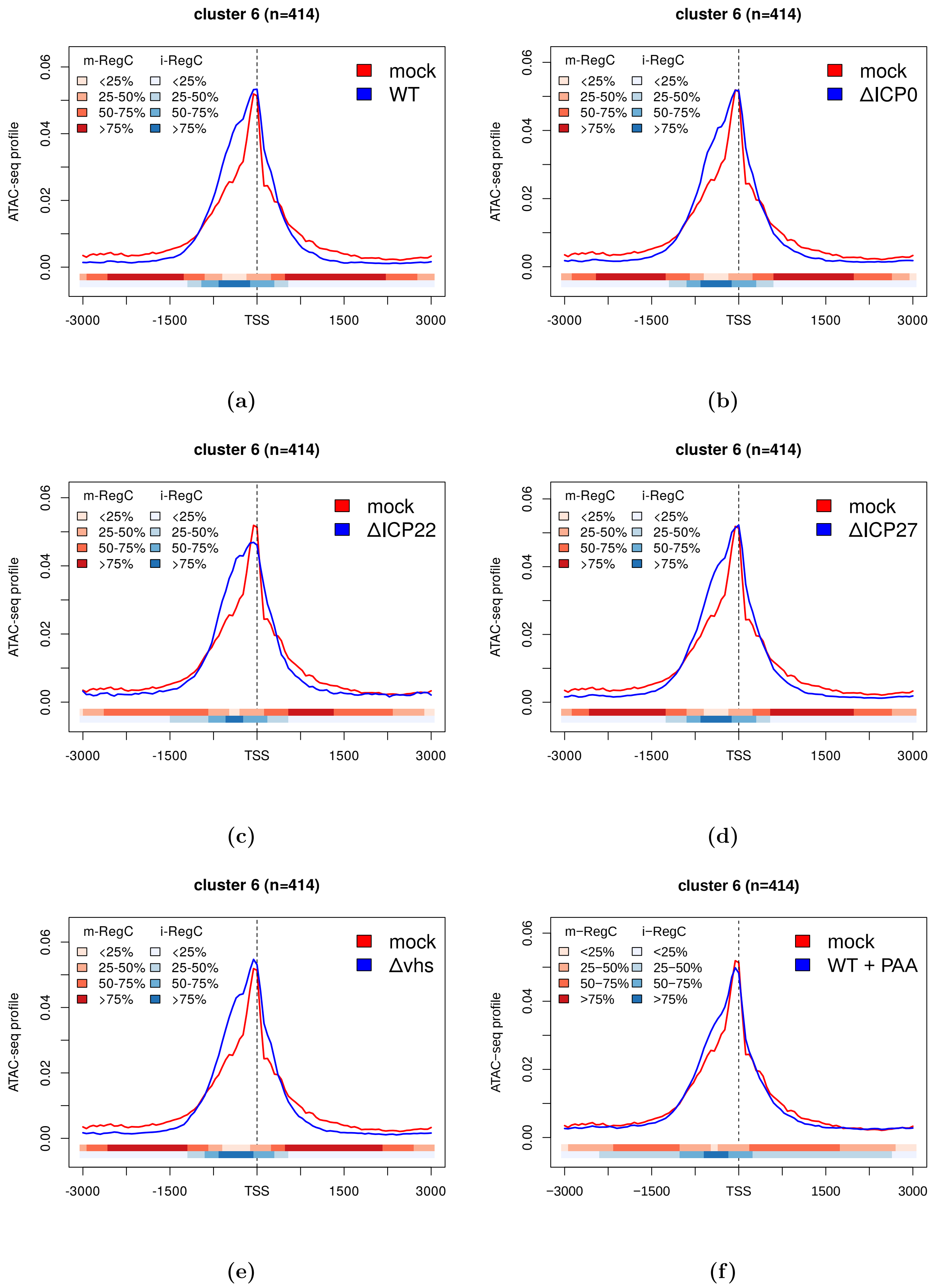


**Sup. Fig 10** Metagene plots of ATAC-seq profiles in mock as well as **(a)** WT infection, **(b-e)** null mutant infections and **(f)** WT+PAA infection for cluster 6 (pattern II). See Materials and Methods for a detailed description of metagene plots. The colored bands below the metagene curves in each subfigure indicate the percentage of genes having an m- or i-RegC (red or blue, respectively) at that position for the corresponding comparison of WT or null mutant virus infection to mock.

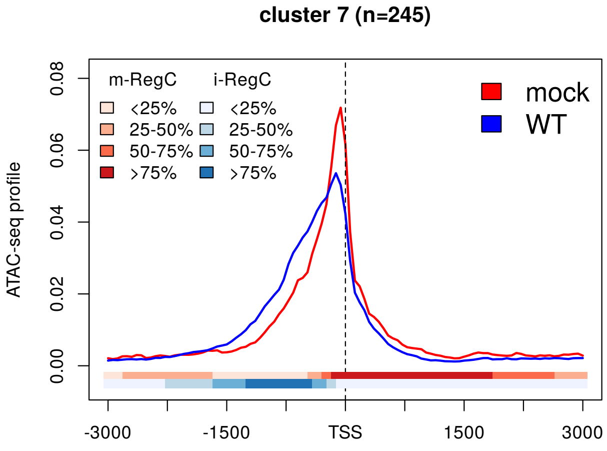


(a)

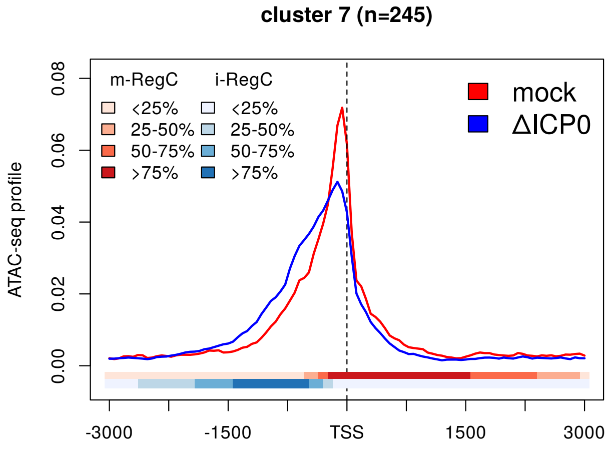


(b)

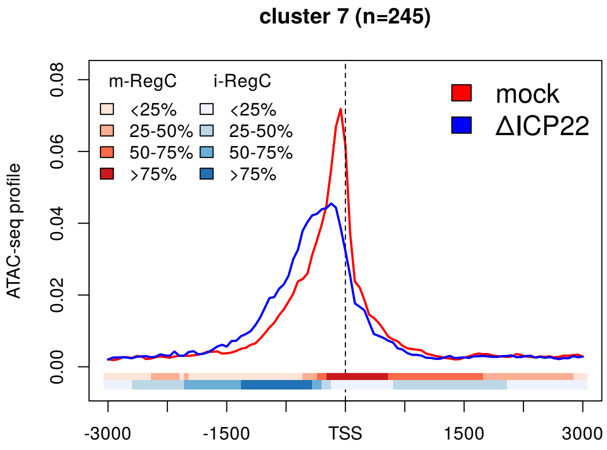


(c)

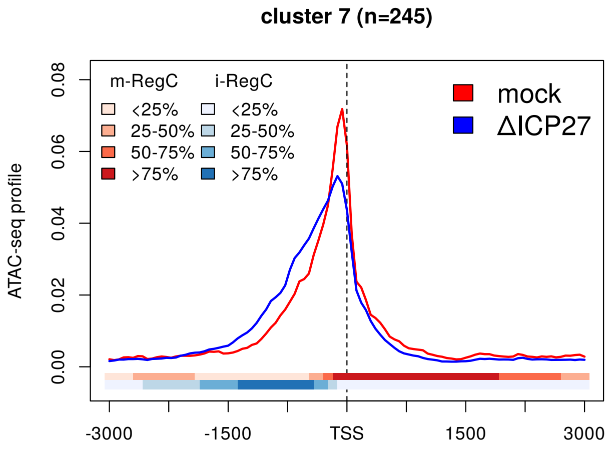


(d)

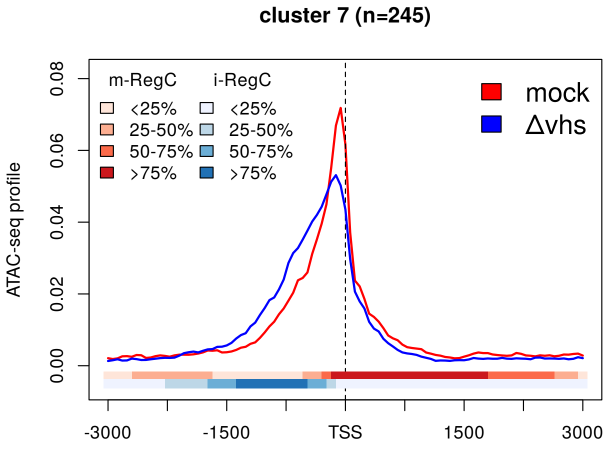


(e)

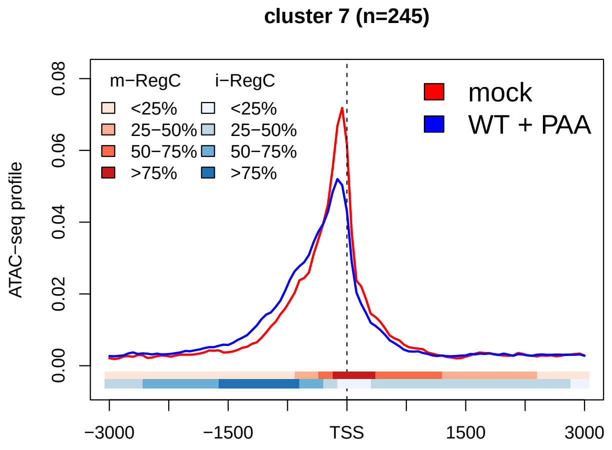


(f)

**Sup. Fig 11** Metagene plots of ATAC-seq profiles in mock as well as **(a)** WT infection, **(b-e)** null mutant infections and **(f)** WT+PAA infection for cluster 7 (pattern II). See Materials and Methods for a detailed description of metagene plots. The colored bands below the metagene curves in each subfigure indicate the percentage of genes having an m- or i-RegC (red or blue, respectively) at that position for the corresponding comparison of WT or null mutant virus infection to mock.

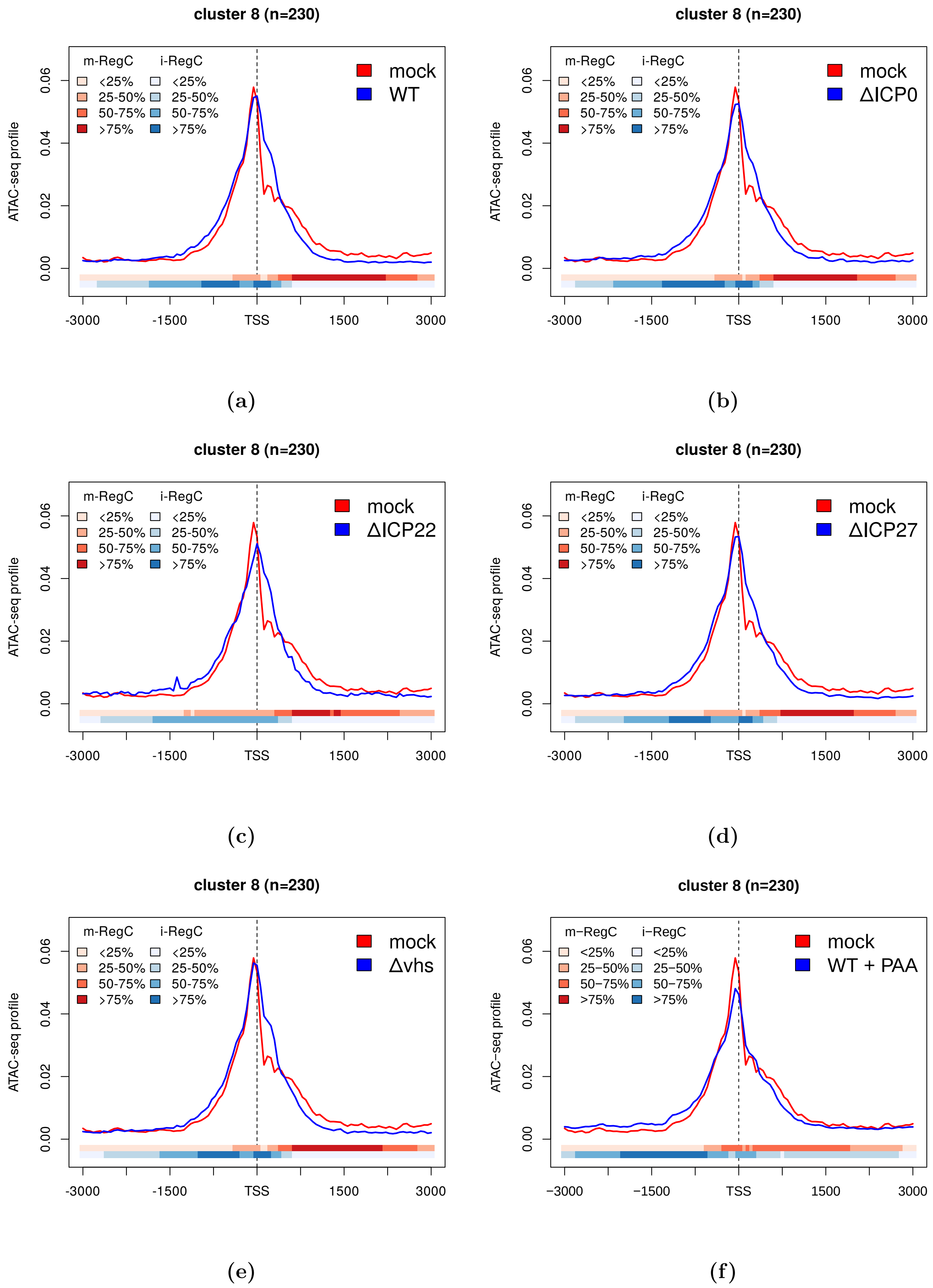


**Sup. Fig 12** Metagene plots of ATAC-seq profiles in mock as well as **(a)** WT infection, **(b-e)** null mutant infections and **(f)** WT+PAA infection for cluster 8 (combined pattern I + II). See Materials and Methods for a detailed description of metagene plots. The colored bands below the metagene curves in each subfigure indicate the percentage of genes having an m- or i-RegC (red or blue, respectively) at that position for the corresponding comparison of WT or null mutant virus infection to mock.

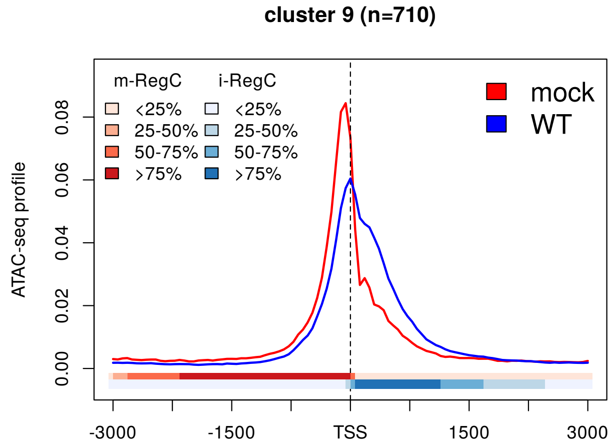


(a)

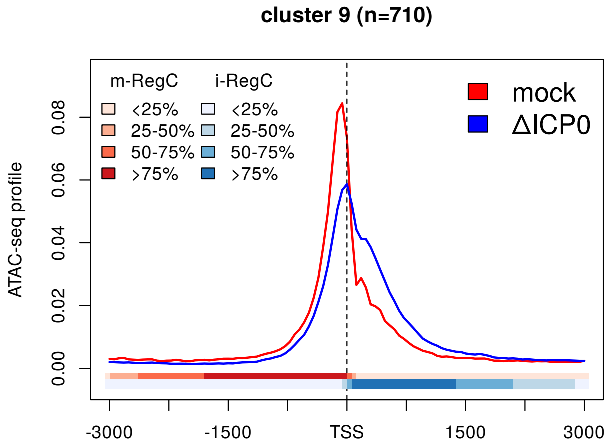


(b)

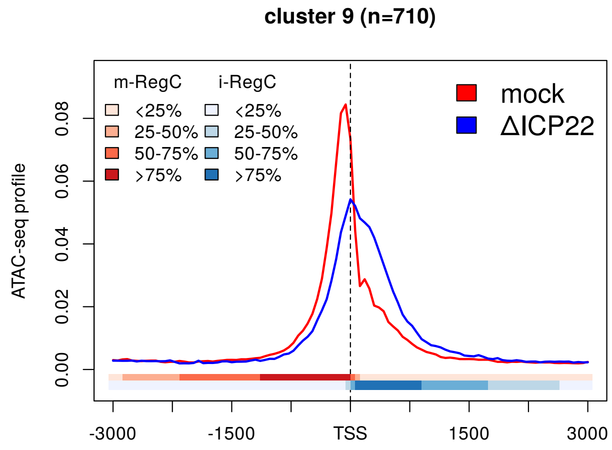


(c)

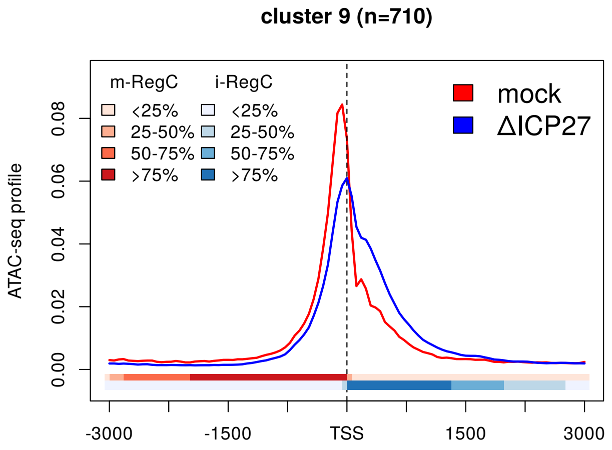


(d)

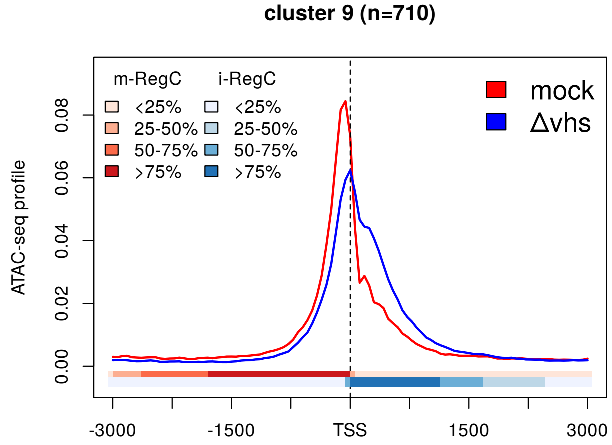


(e)

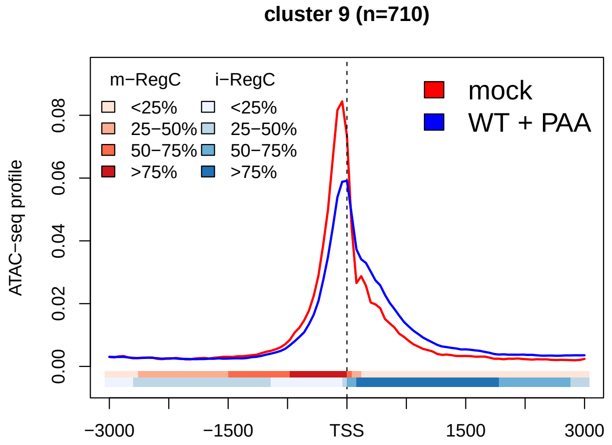


(f)

**Sup. Fig 13** Metagene plots of ATAC-seq profiles in mock as well as **(a)** WT infection, **(b-e)** null mutant infections and **(f)** WT+PAA infection for cluster 9 (pattern I). See Materials and Methods for a detailed description of metagene plots. The colored bands below the metagene curves in each subfigure indicate the percentage of genes having an m- or i-RegC (red or blue, respectively) at that position for the corresponding comparison of WT or null mutant virus infection to mock.

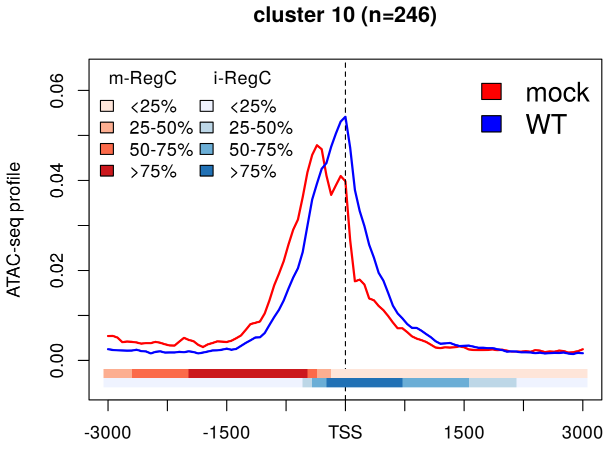


(a)

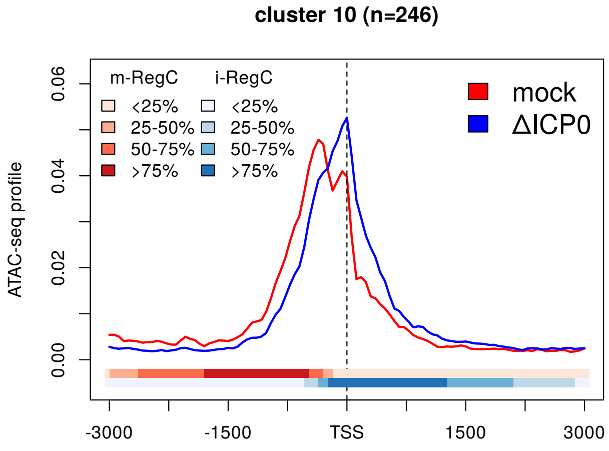


(b)

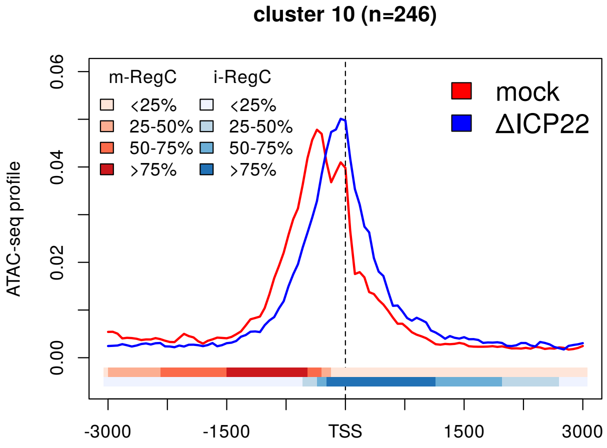


(c)

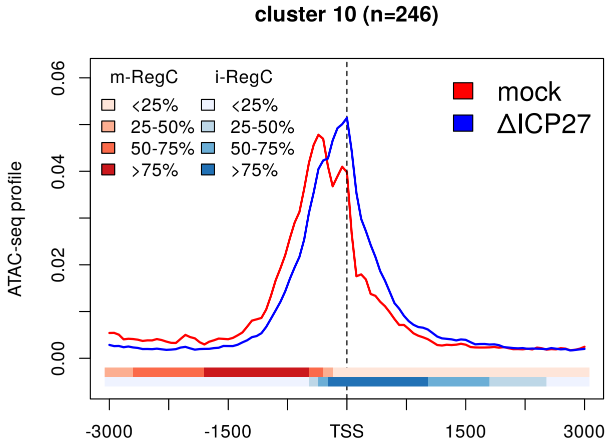


(d)

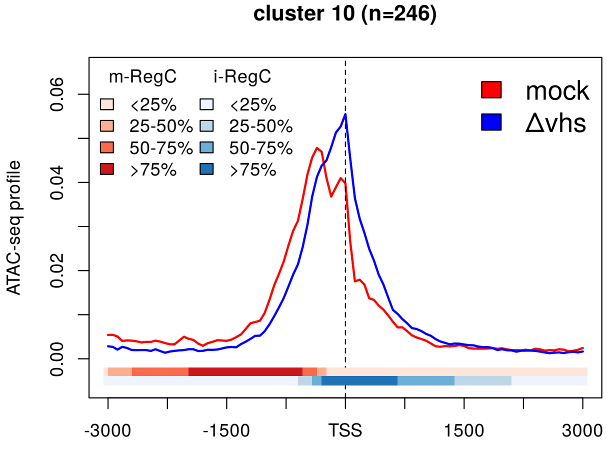


(e)

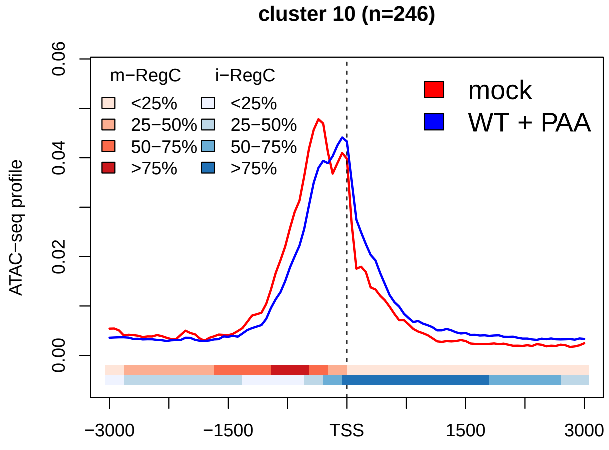


(f)

**Sup. Fig 14** Metagene plots of ATAC-seq profiles in mock as well as **(a)** WT infection, **(b-e)** null mutant infections and **(f)** WT+PAA infection for cluster 10 (pattern I). See Materials and Methods for a detailed description of metagene plots. The colored bands below the metagene curves in each subfigure indicate the percentage of genes having an m- or i-RegC (red or blue, respectively) at that position for the corresponding comparison of WT or null mutant virus infection to mock.

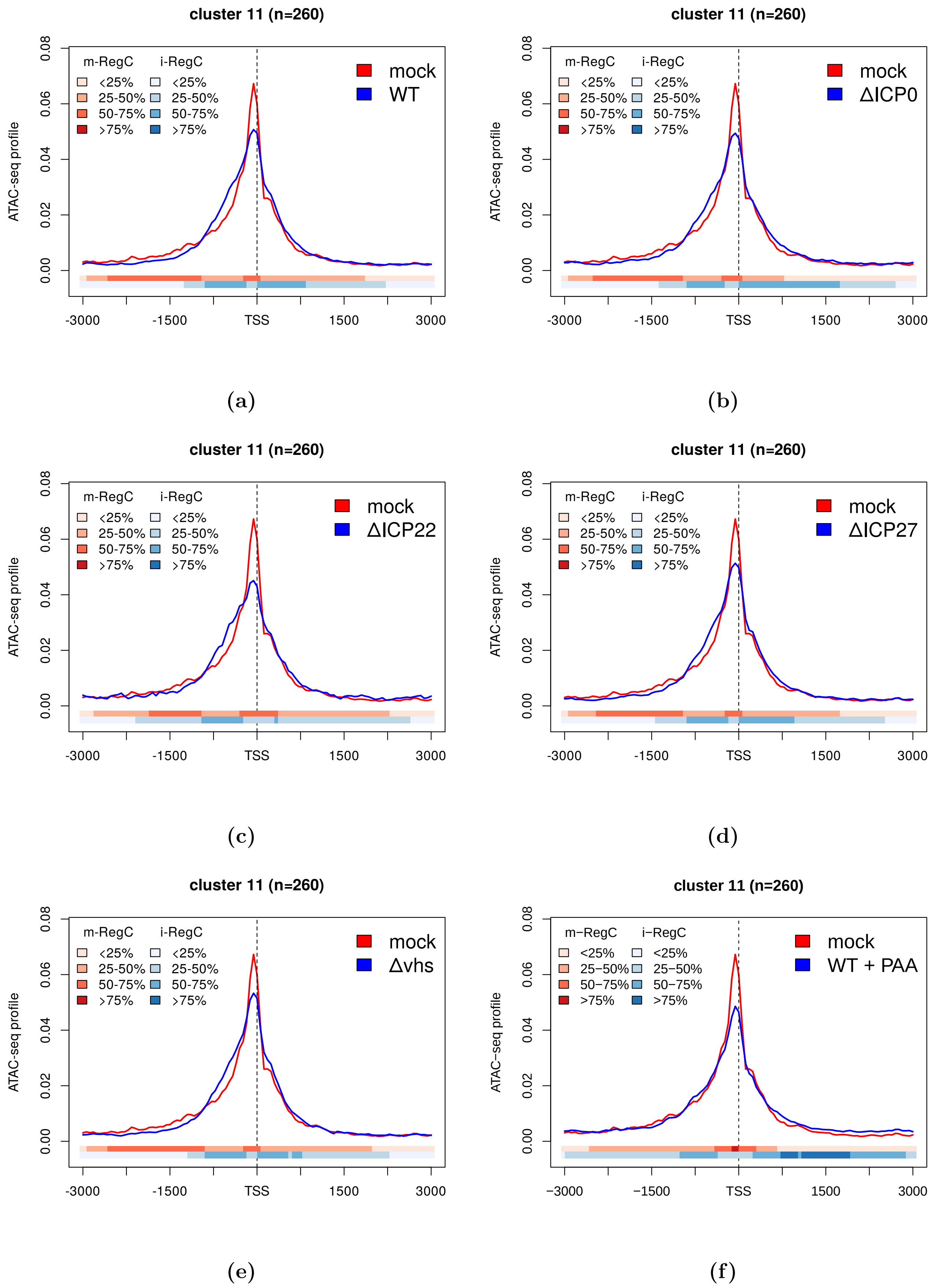


**Sup. Fig 15** Metagene plots of ATAC-seq profiles in mock as well as **(a)** WT infection, **(b-e)** null mutant infections and **(f)** WT+PAA infection for cluster 11 (pattern II). See Materials and Methods for a detailed description of metagene plots. The colored bands below the metagene curves in each subfigure indicate the percentage of genes having an m- or i-RegC (red or blue, respectively) at that position for the corresponding comparison of WT or null mutant virus infection to mock.

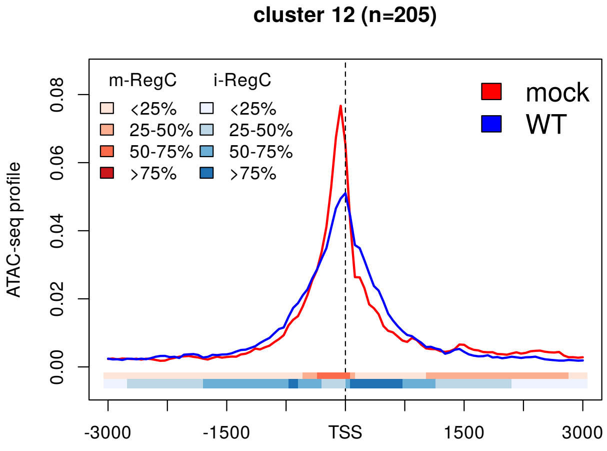


(a)

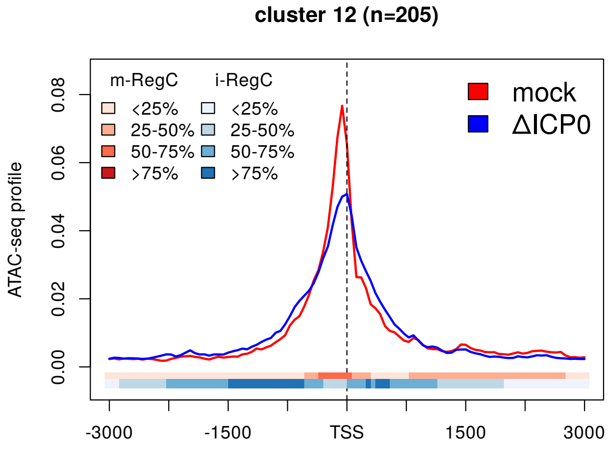


(b)

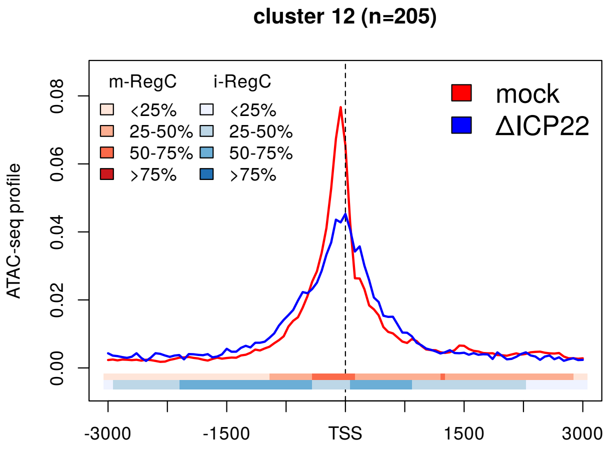


(c)

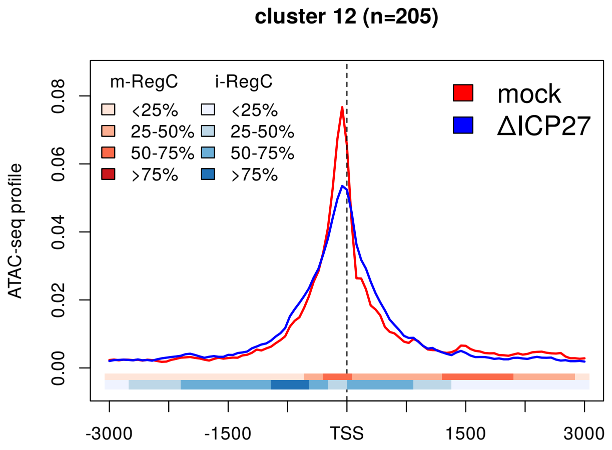


(d)

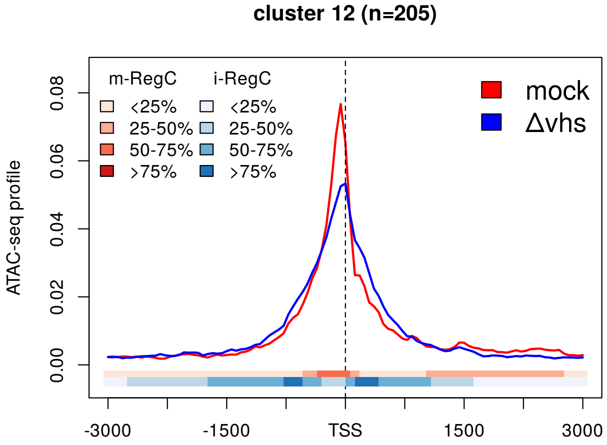


(e)

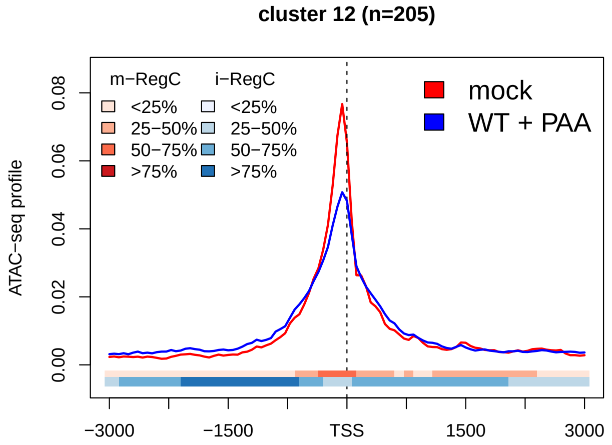


(f)

**Sup. Fig 16** Metagene plots of ATAC-seq profiles in mock as well as **(a)** WT infection, **(b-e)** null mutant infections and **(f)** WT+PAA infection for cluster 12 (pattern I). See Materials and Methods for a detailed description of metagene plots. The colored bands below the metagene curves in each subfigure indicate the percentage of genes having an m- or i-RegC (red or blue, respectively) at that position for the corresponding comparison of WT or null mutant virus infection to mock.

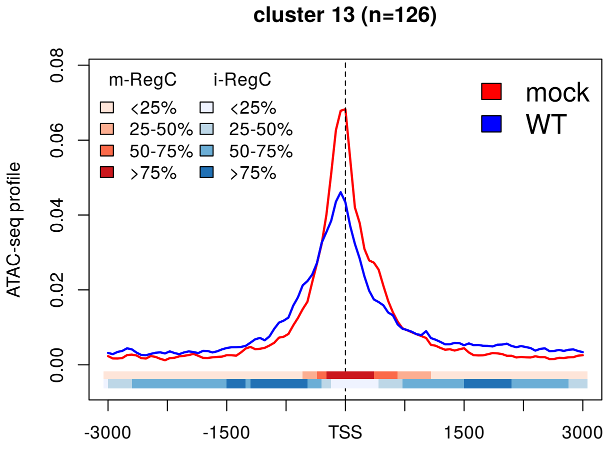


(a)

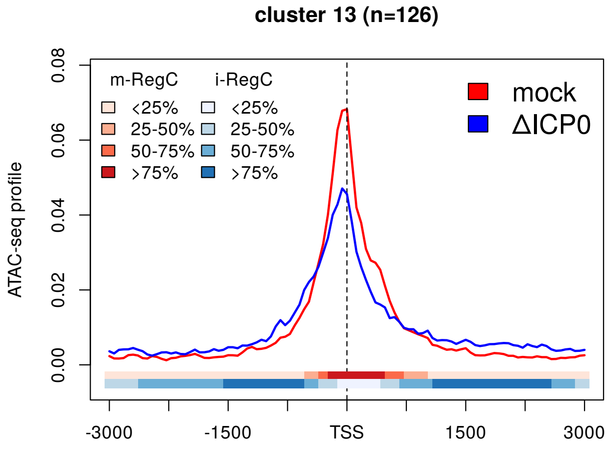


(b)

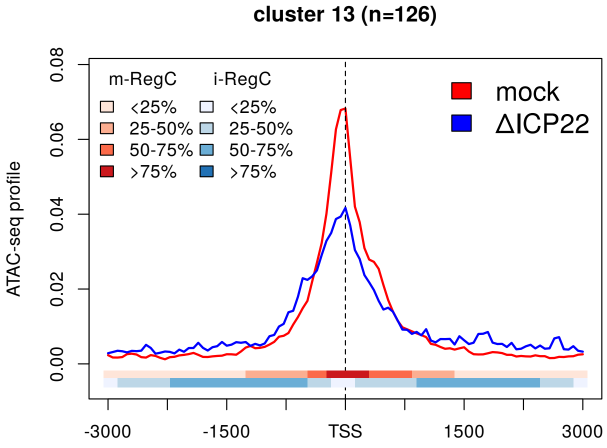


(c)

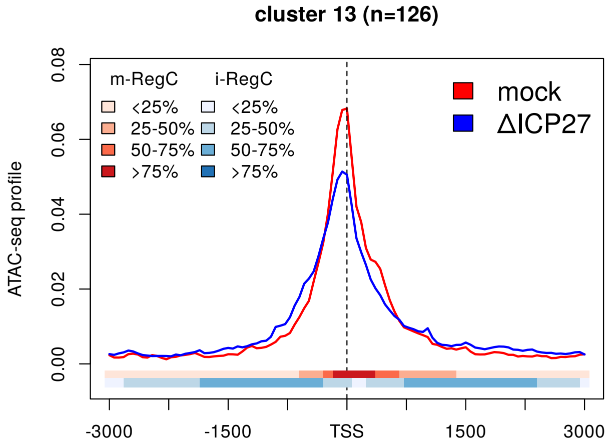


(d)

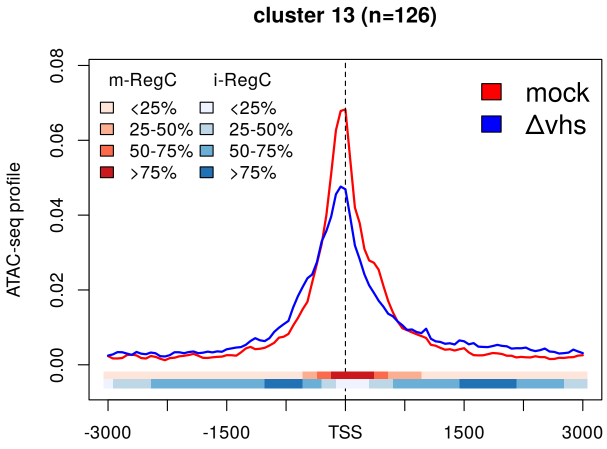


(e)

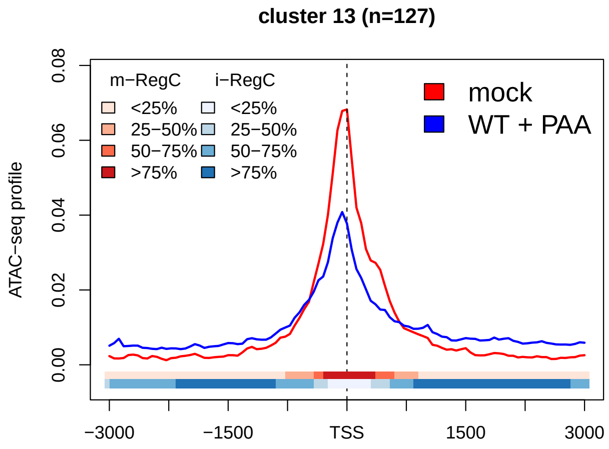


(f)

**Sup. Fig 17** Metagene plots of ATAC-seq profiles in mock as well as **(a)** WT infection, **(b-e)** null mutant infections and **(f)** WT+PAA infection for cluster 13 (pattern III). See Materials and Methods for a detailed description of metagene plots. The colored bands below the metagene curves in each subfigure indicate the percentage of genes having an m- or i-RegC (red or blue, respectively) at that position for the corresponding comparison of WT or null mutant virus infection to mock.

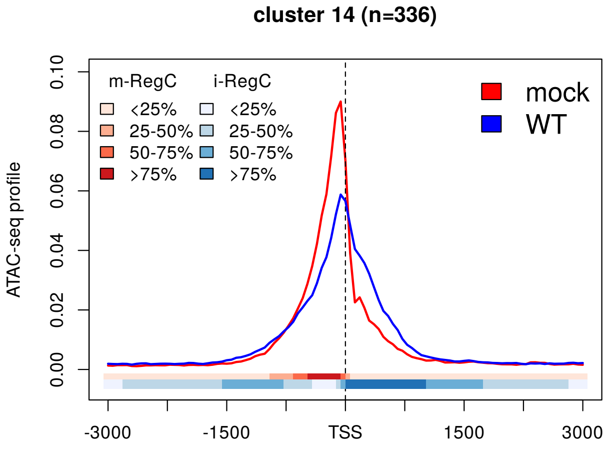


(a)

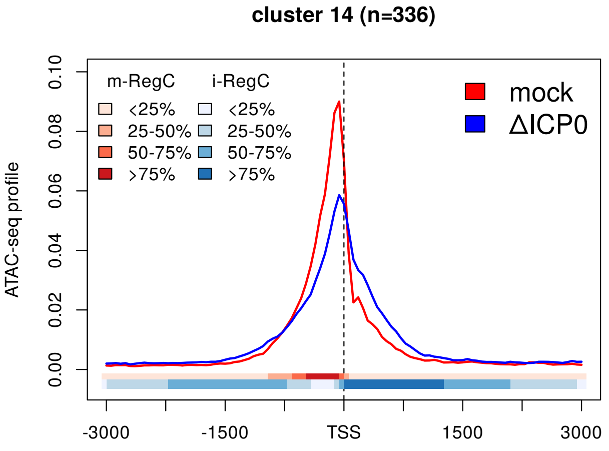


(b)

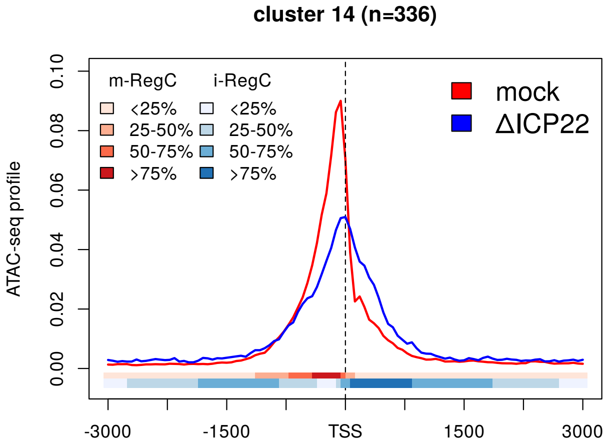


(c)

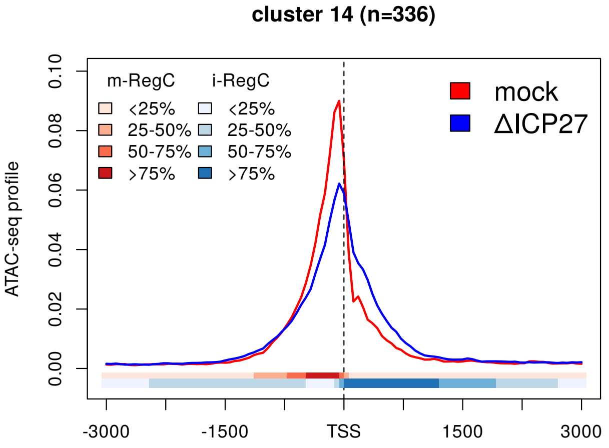


(d)

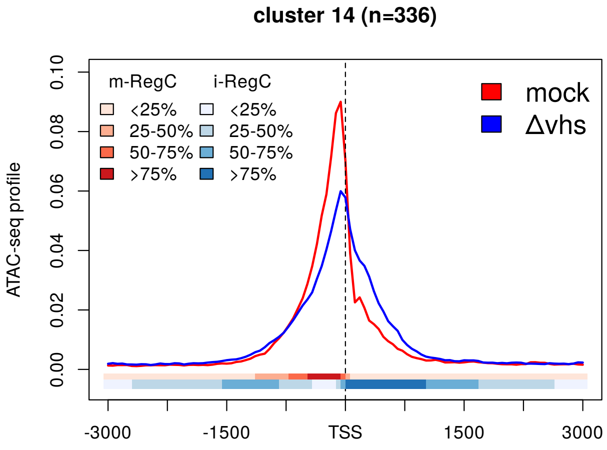


(e)

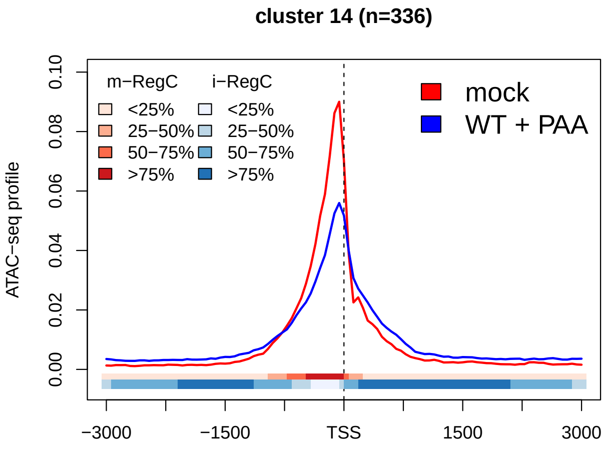


(f)

**Sup. Fig 18** Metagene plots of ATAC-seq profiles in mock as well as **(a)** WT infection, **(b-e)** null mutant infections and **(f)** WT+PAA infection for cluster 14 (pattern I). See Materials and Methods for a detailed description of metagene plots. The colored bands below the metagene curves in each subfigure indicate the percentage of genes having an m- or i-RegC (red or blue, respectively) at that position for the corresponding comparison of WT or null mutant virus infection to mock.

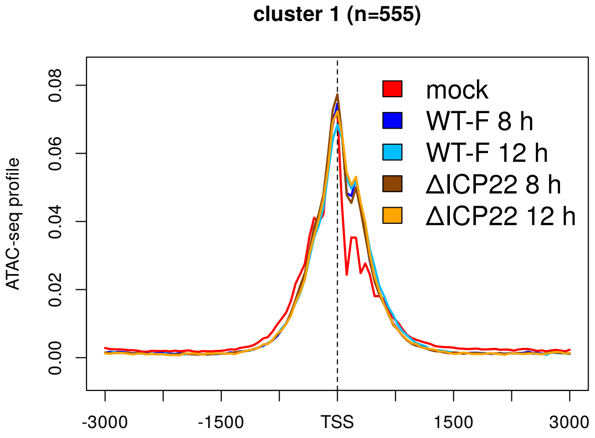


**(a)** pattern I

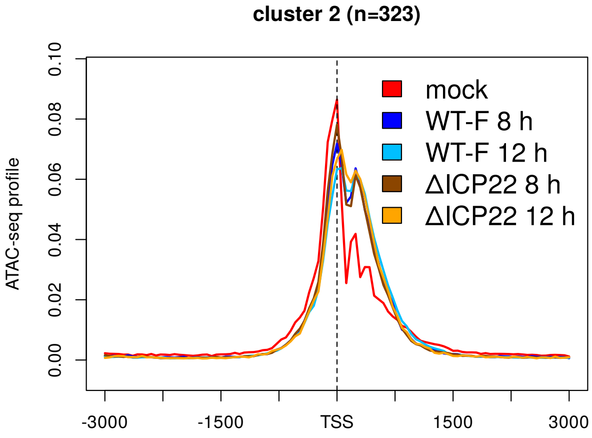


**(b)** pattern I

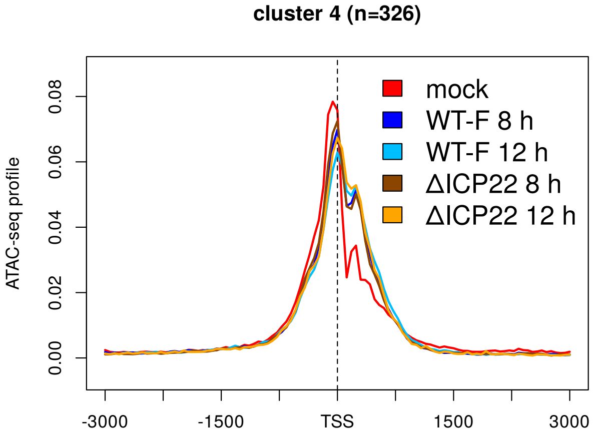


**(c)** pattern I

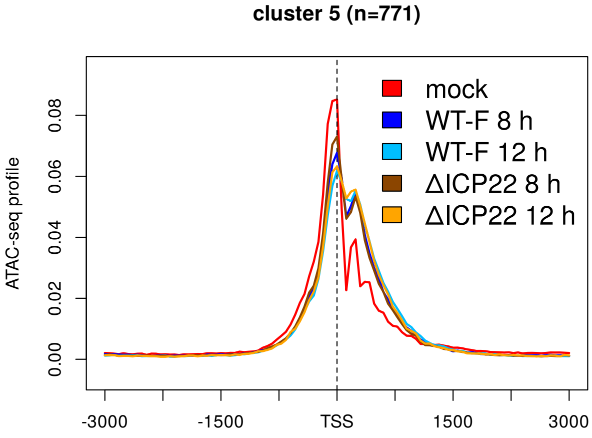


**(d)** pattern I

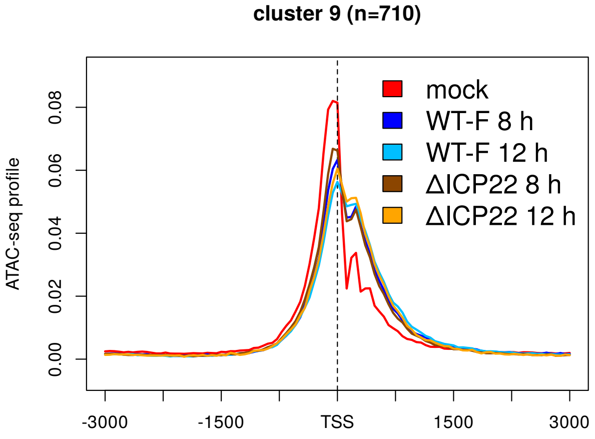


**(e)** pattern I

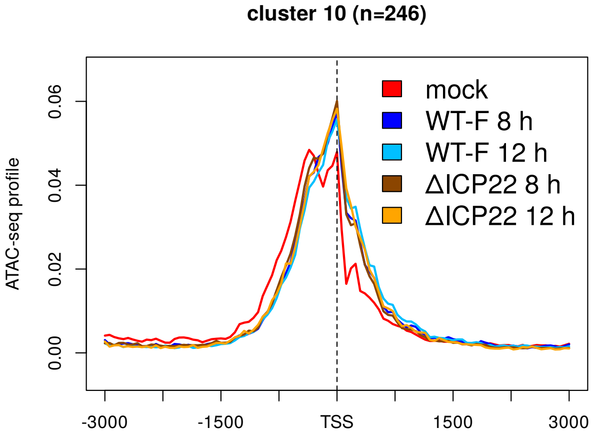


**(f)** pattern I

**(Continued on next page)**

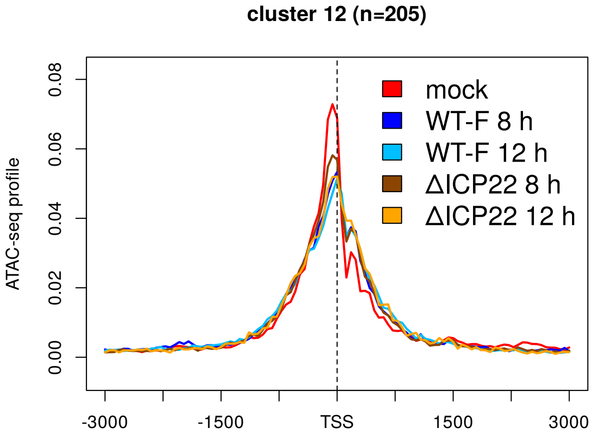


**(g)** pattern I

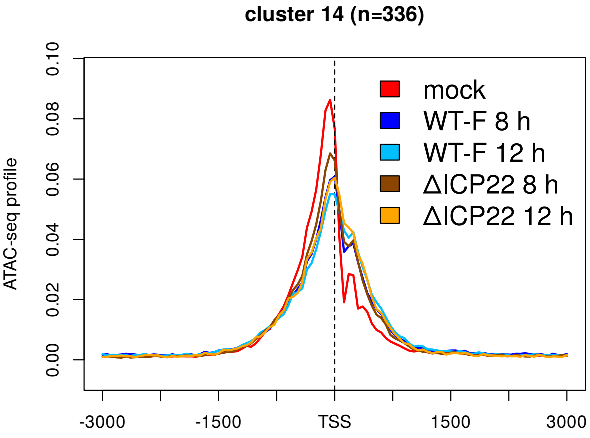


**(h)** pattern I

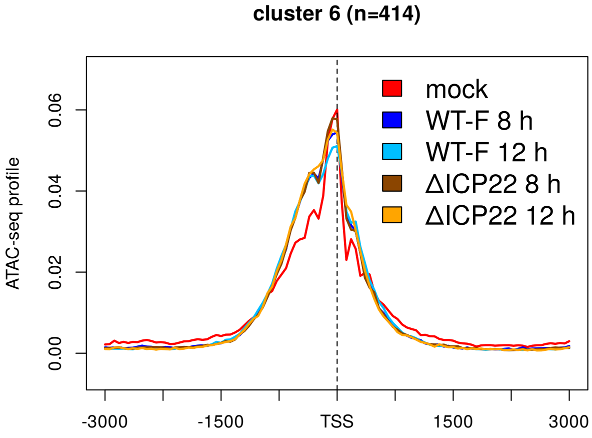


**(i)** pattern II

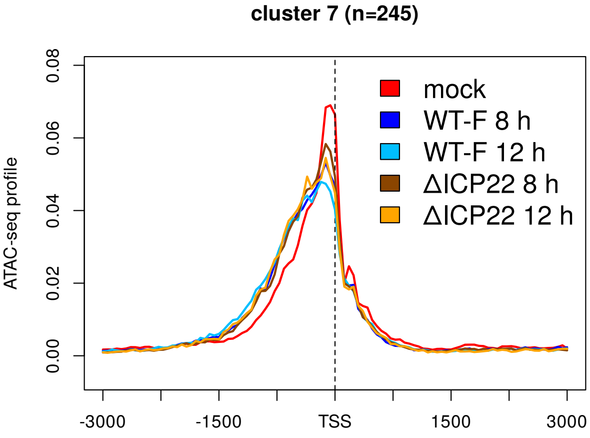


**(j)** pattern II

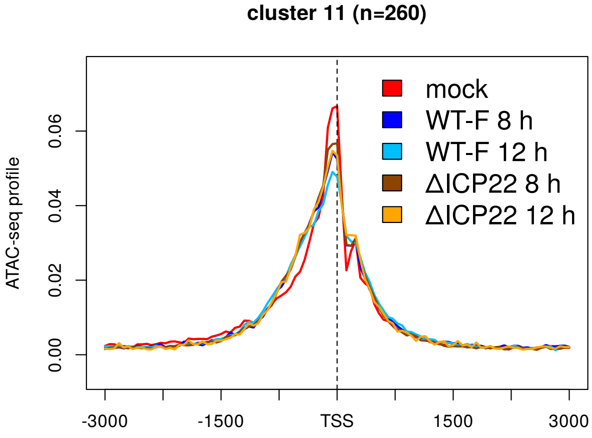


**(k)** pattern II

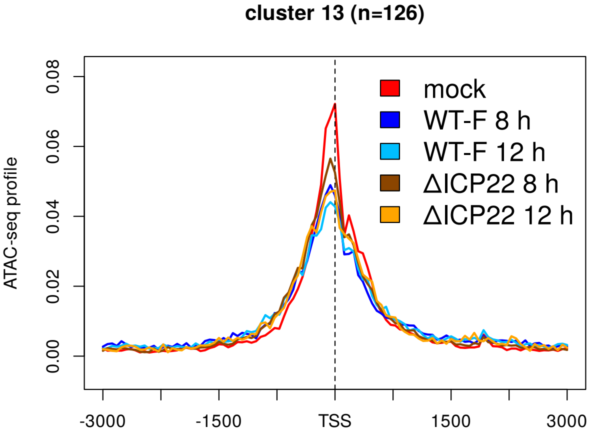


**(l)** pattern III

**(Continued on next page)**

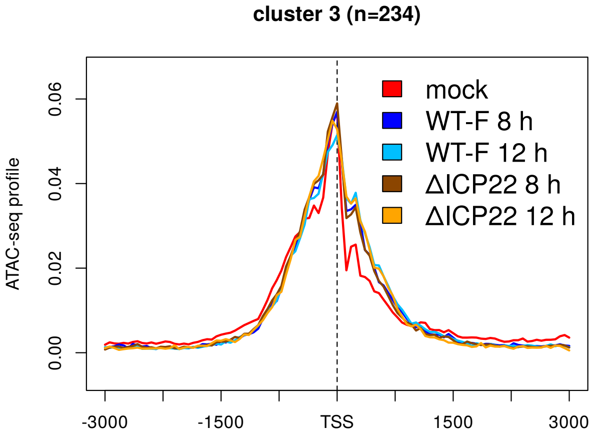


**(m)** pattern I + II

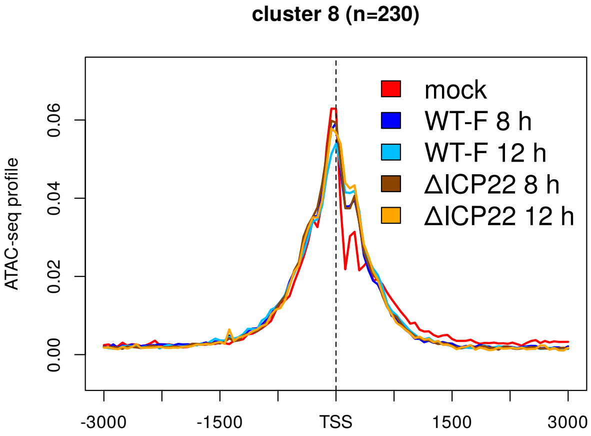


**(n)** pattern I + II

**Sup. Fig 19** Metagene plots of ATAC-seq profiles in mock, WT strain F (WT-F) and ΔICP22 infection at 8 and 12 h p.i. for all clusters (indicated on top of subfigures) ordered according to patterns (indicated below subfigures). See Materials and Methods for a detailed description of metagene plots.

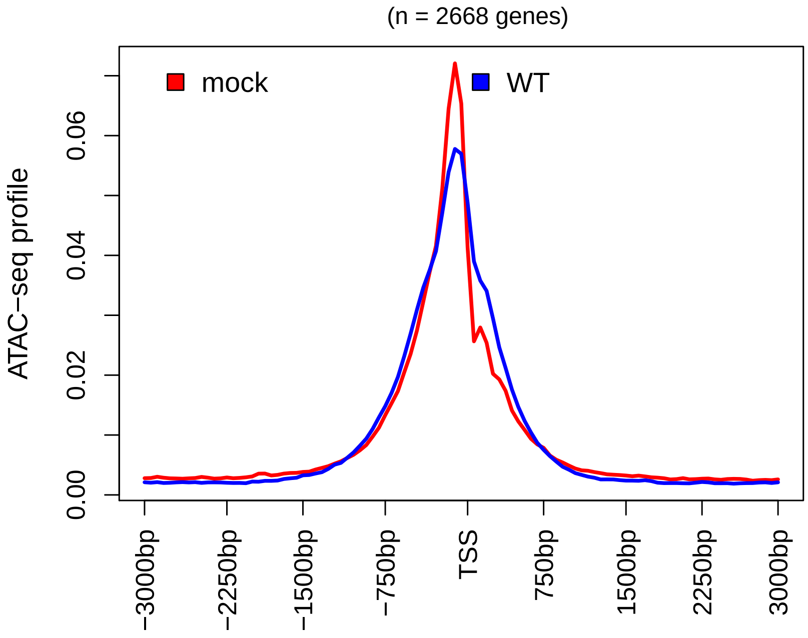


**(a)**

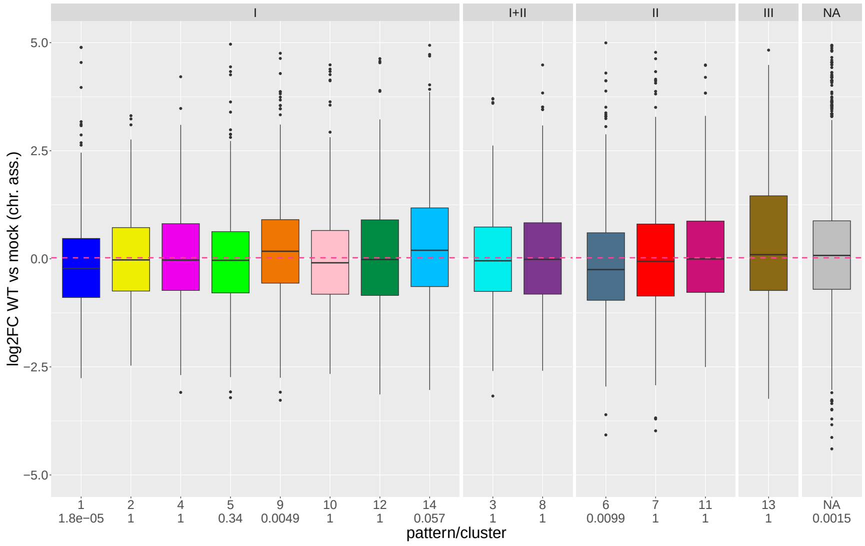


**(b)**

**Sup. Fig 20 (a)** Metagene plot of the ATAC-seq profiles for mock (red) and WT (blue) infection for all promoter windows for which no significant RegC was identified in the ATAC-seq data for any of the WT or null mutant infections compared to mock (denoted as NA group in **(b)**). These genes were not included in **Fig. 1a** and most analyses in this article. See Materials and Methods for an explanation of metagene plots. **(b)** Boxplot showing the distribution of log2 fold-changes in chromatin-associated RNA for 8 h p.i. WT infection compared to mock for all clusters (grouped by pattern) as well as genes without significant differential regions (NA group). P-values for Wilcoxon rank sum tests comparing log2 fold-changes for each group against all other analyzed genes were corrected for multiple testing using the Bonferroni method and are shown below each gene group.

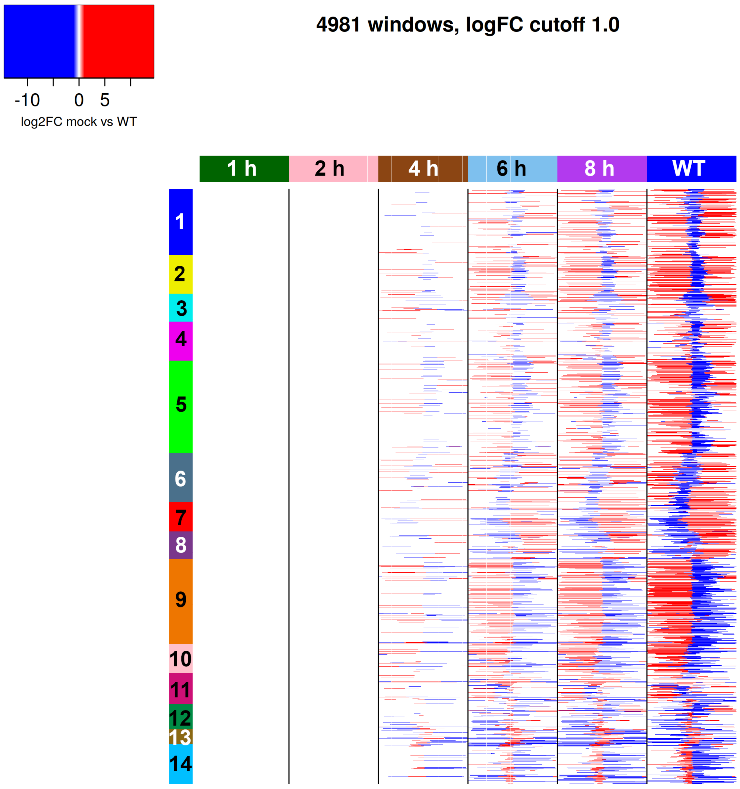


**Sup. Fig 21** Heatmap showing the log2 fold-changes determined with DEXSeq on the ATAC-seq time-course data (1, 2, 4, 6 and 8 h p.i. WT infection compared to mock) for the differential regions (m- and i-RegC) determined in the WT vs. mock comparison shown in **Fig. 1a**. Each row shows log2 fold-changes for the same m- and i-RegCs at the different time-points of the time-course compared to mock. For comparison, log2 fold-changes for the WT vs. mock comparison from **Fig. 1a**/**Sup. Fig. 1d** are also shown. Colored rectangles on top indicate the time-point or whether the WT vs. mock comparison is shown. Statistically significant differential regions are colored according to the log2 fold-change determined by DEXSeq. Here, the color scale is continuous between -1 and 1 and all log2 fold-changes > 1 are colored the same red and all log2 fold-changes < 1 the same blue. Promoter windows are ordered as in **Fig. 1a** and clusters from **Fig. 1a** are annotated as colored and numbered rectangles on the left.

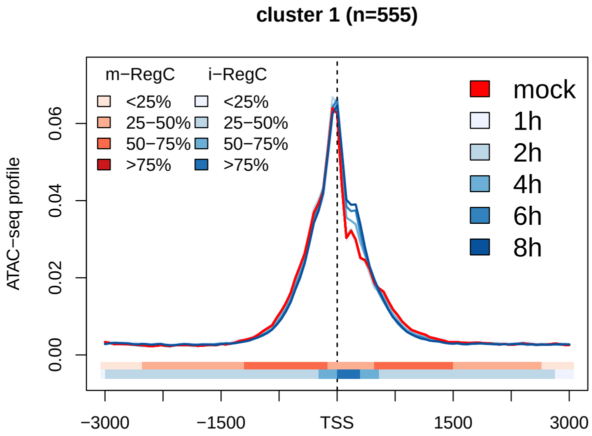


(a) pattern I

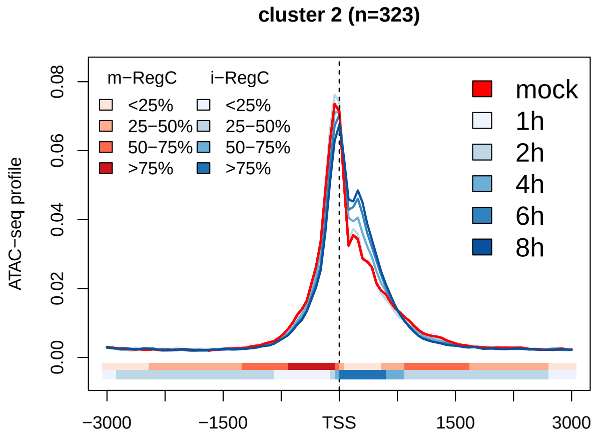


(b) pattern I

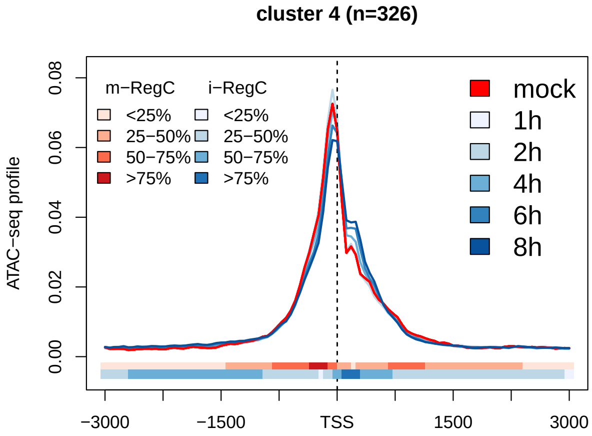


(c) pattern I

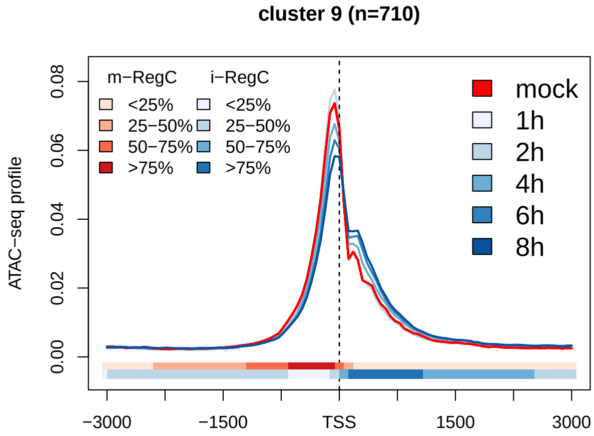


(d) pattern I

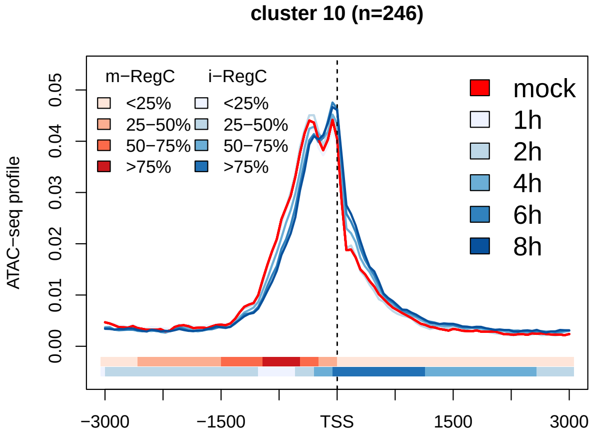


(e) pattern I

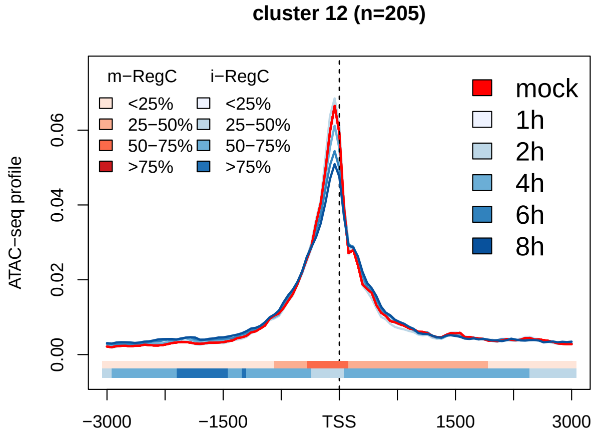


(f) pattern I

**(Continued on next page)**

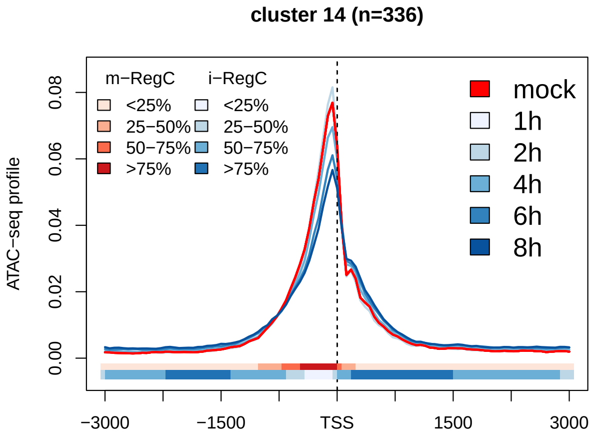


(g) pattern I

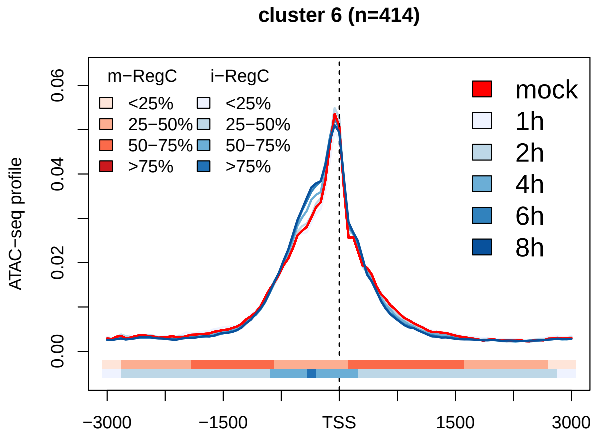


(h) pattern II

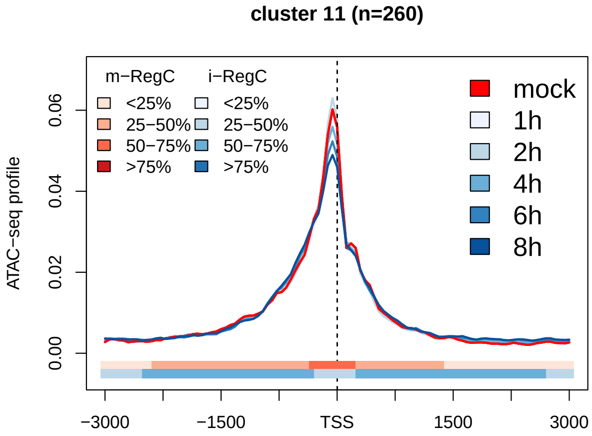


(i) pattern II

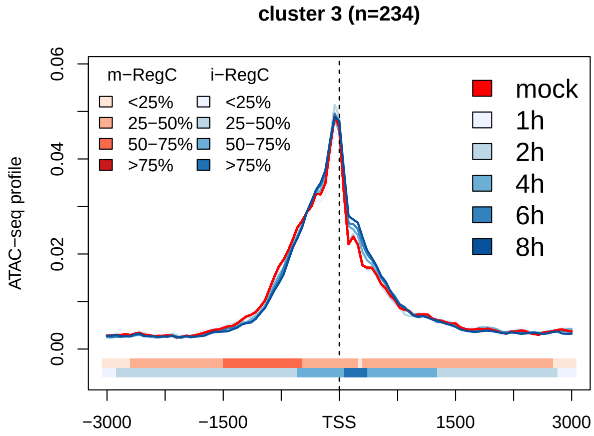


(j) pattern I + II

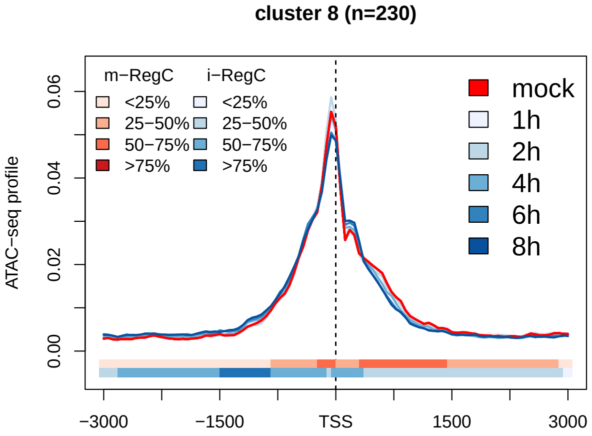


(k) pattern I + II

**Sup. Fig 22** Metagene plots of ATAC-seq profiles in mock infection (red) and all time-points of infection (blue shades) from the ATAC-seq time-course experiment for all clusters (indicated on top of subfigures) ordered according to patterns (indicated below subfigures). See Materials and Methods for a detailed description of metagene plots. The colored bands below the metagene curves in each subfigure indicate the percentage of genes having an m- or i-RegC (red or blue, respectively) at that position for the comparison of 8 h p.i. to mock.

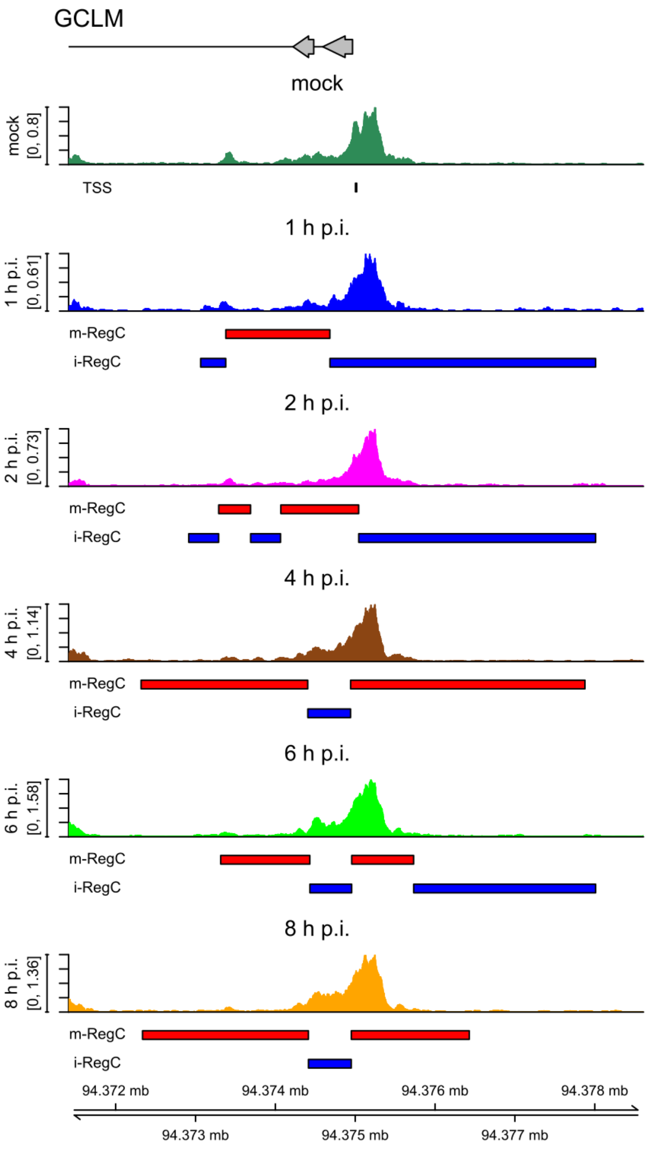


**(a)** pattern I, cluster 2

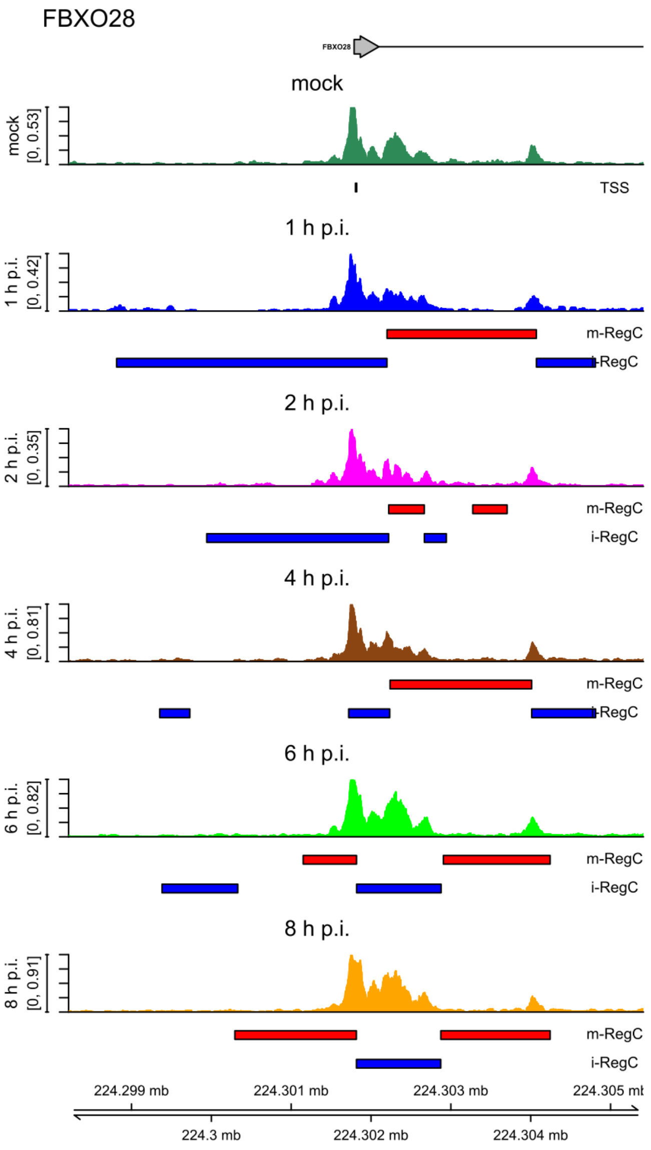


**(b)** pattern I, cluster 5

**(Continued on next page)**

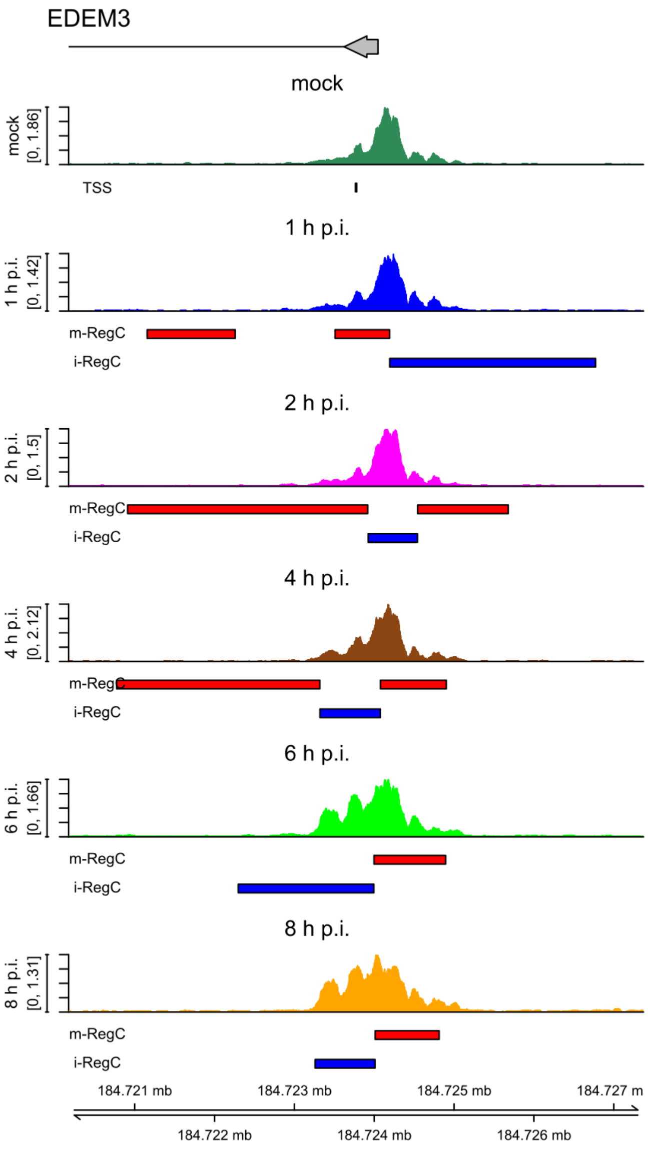


(c) pattern I, cluster 10

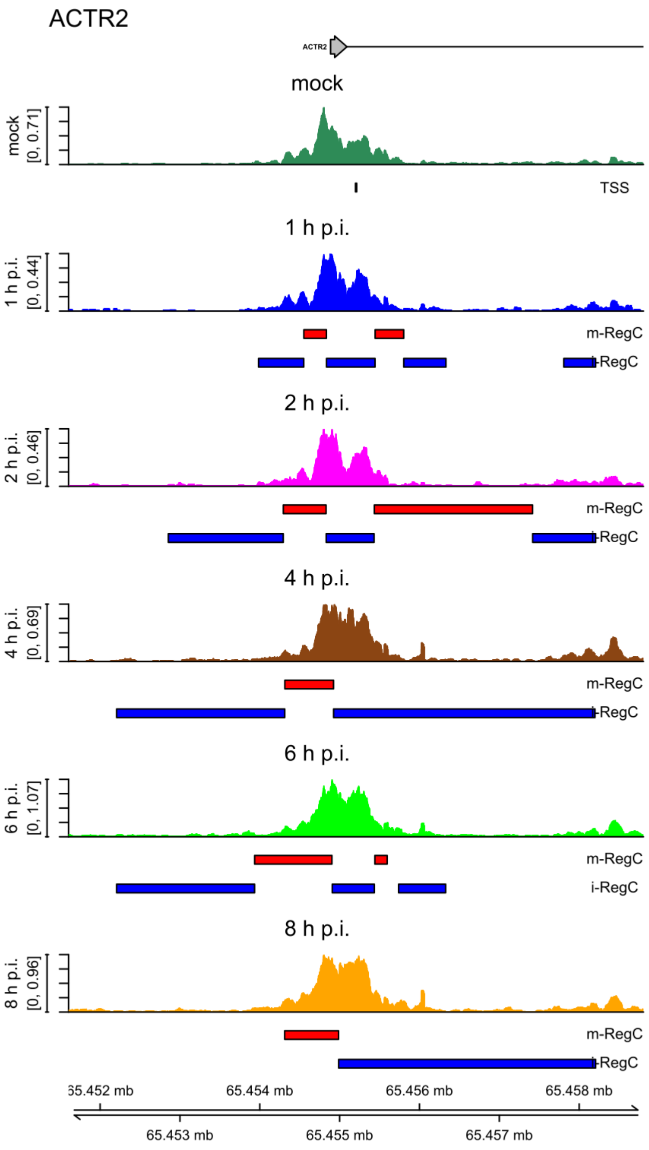


(d) pattern I, cluster 12

**(Continued on next page)**

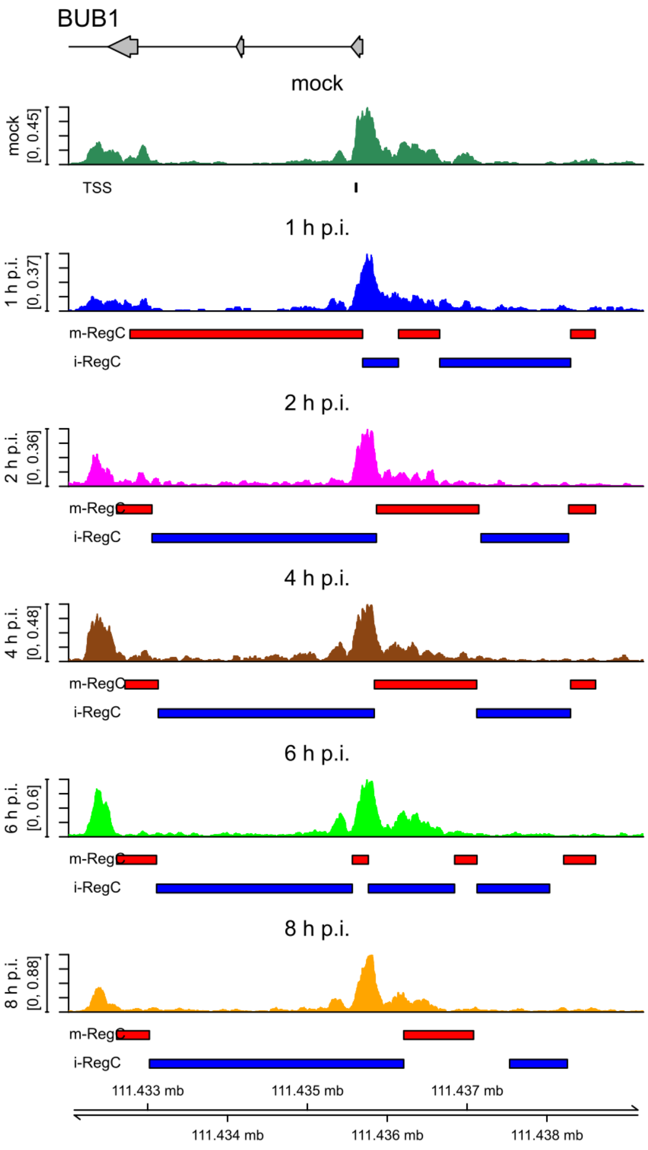


(e) pattern II, cluster 6

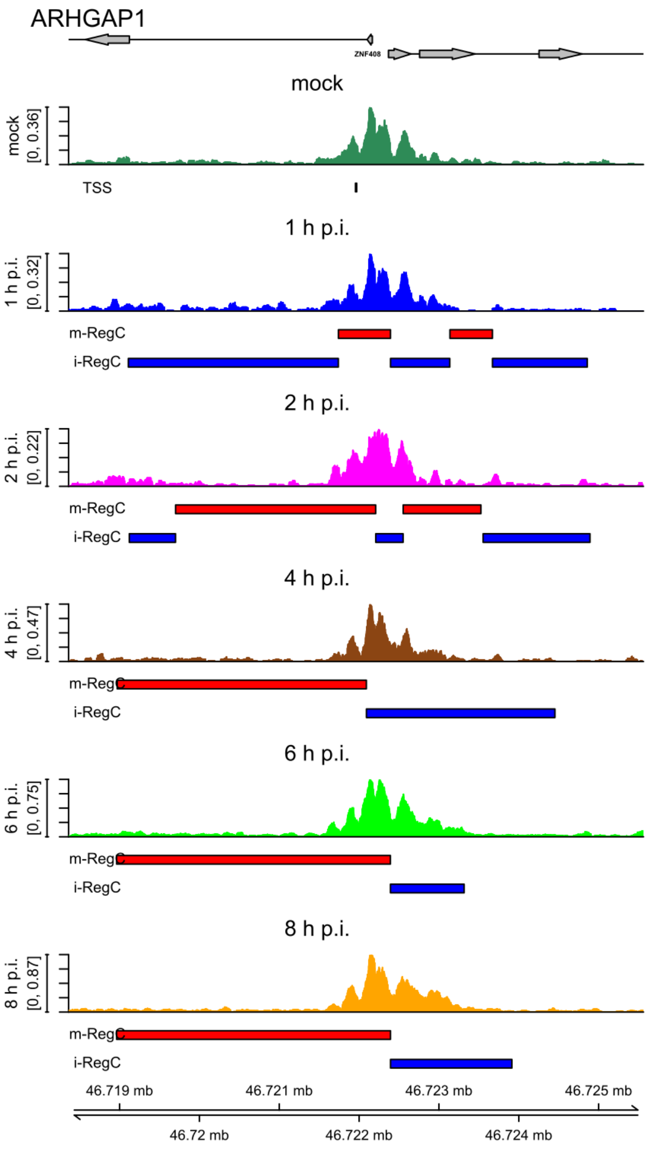


(f) pattern II, cluster 7

**(Continued on next page)**

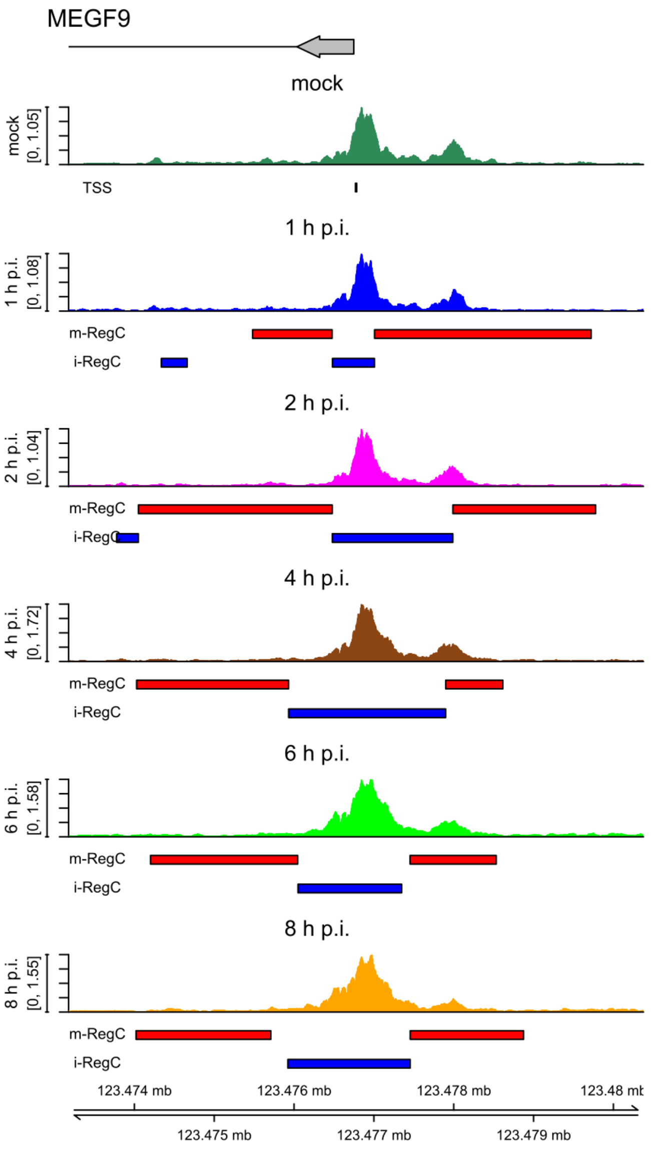


(g) pattern I + II, cluster 3

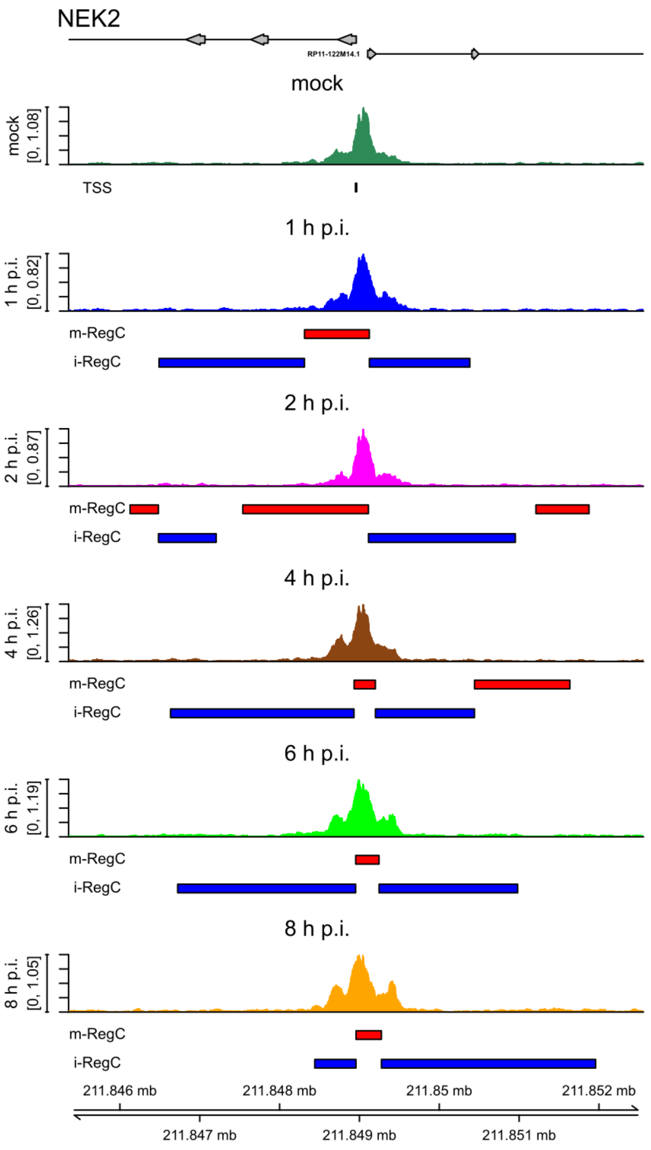


**(h)** pattern I + II, cluster 8

**Sup. Fig 23** Read coverage in a ± 3.6 kb window around the TSS in ATAC-seq data for mock (dark green), 1 h (blue), 2 h (magenta), 4 h (brown), 6 h (green), and 8 h p.i. (orange) for example genes with pattern I **(a-d)**, pattern II **(e,f)** and combined patterns I and II **(g,h)**. Read coverage was normalized to total number of mapped reads for each sample and averaged between replicates. The TSS used in the analysis is indicated by a short vertical line below the read coverage track for mock infection. Gene annotation is indicated at the top. Boxes represent exons, lines represent introns, and direction of transcription is indicated by arrowheads. The name of the gene whose promoter window was analyzed is indicated in larger font on the top left and sometimes beside the gene annotation. Names for other genes overlapping the input window are also indicated. Below each read coverage track m- (red bars) and i-RegCs (blue bars) are indicated for the comparison of the corresponding time-point of infection to mock. Corresponding read coverage plots in ATAC-seq data for mock, WT, ΔICP0, ΔICP22, ΔICP27, Δ*vhs* and WT+PAA infection are shown in **Sup. Fig. 4**.

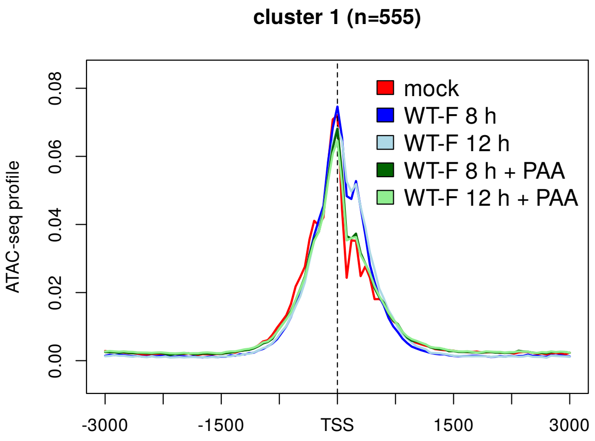


(a) pattern I

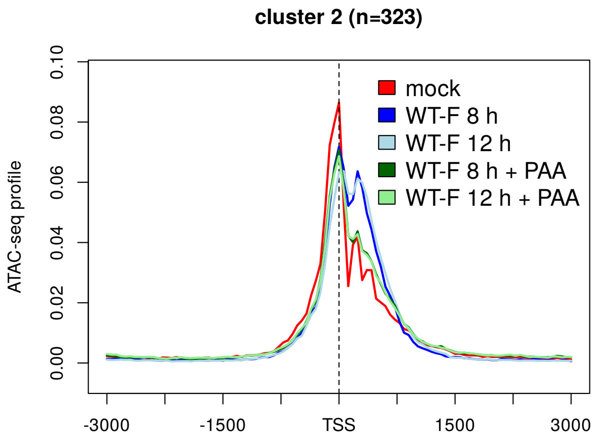


(b) pattern I

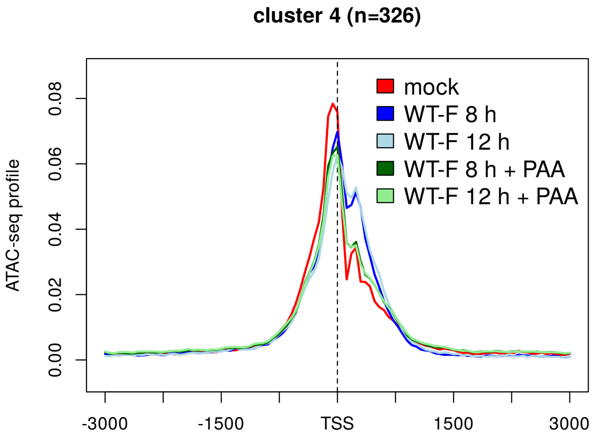


(c) pattern I

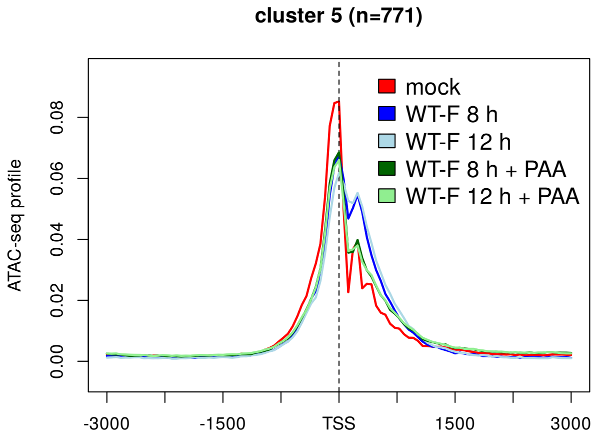


(d) pattern I

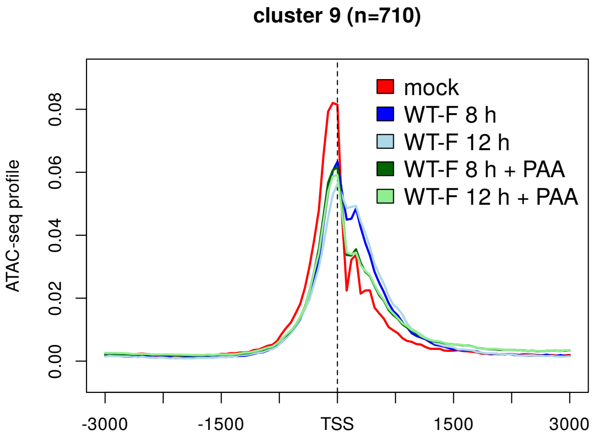


(e) pattern I

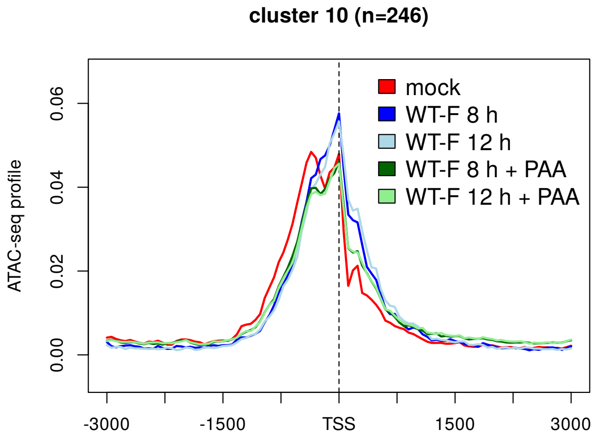


(f) pattern I

(Continued on next page)

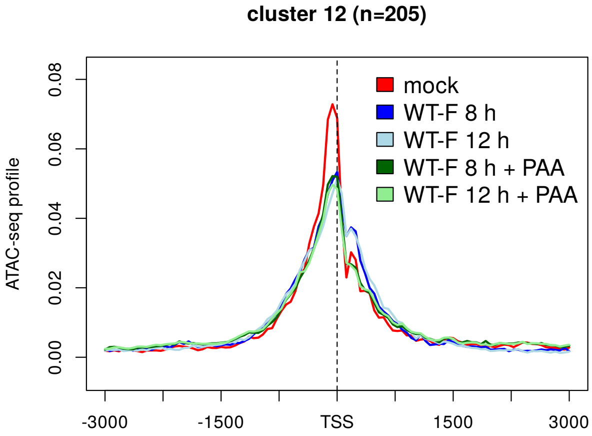


**(g)** pattern I

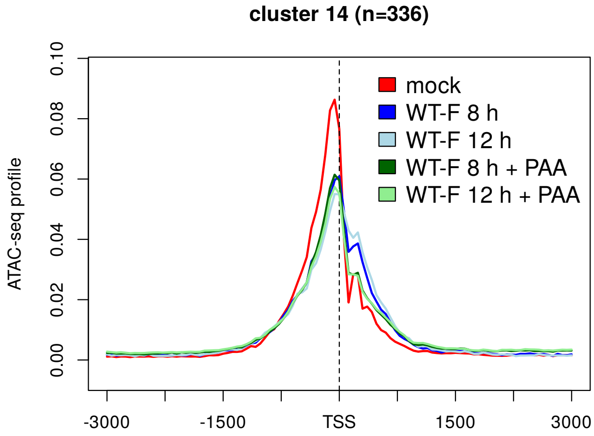


**(h)** pattern I

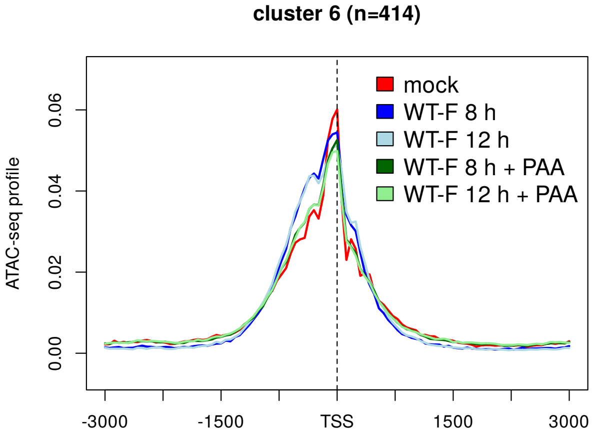


**(i)** pattern II

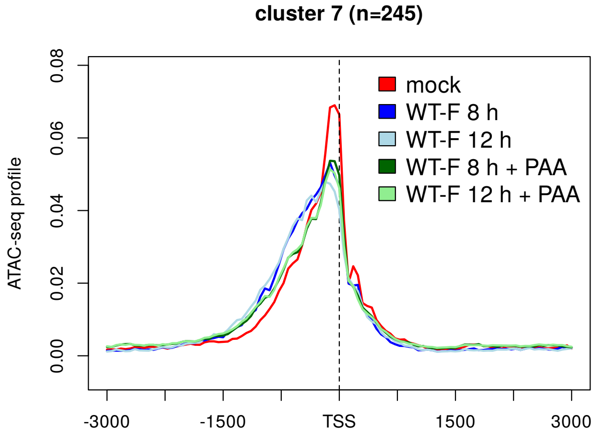


**(j)** pattern II

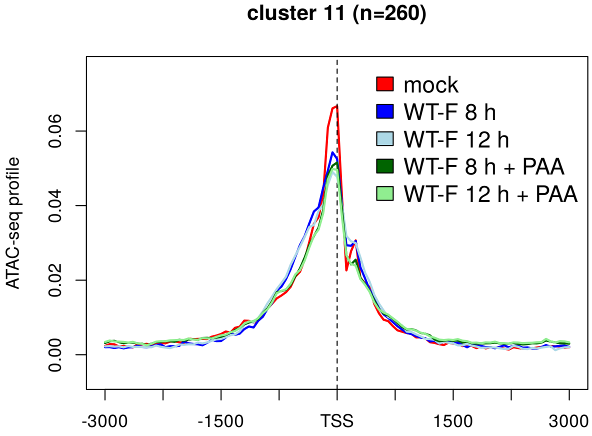


**(k)** pattern II

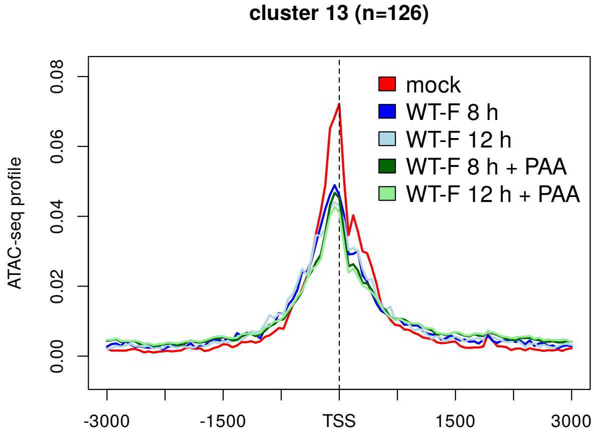


**(l)** pattern III

**(Continued on next page)**

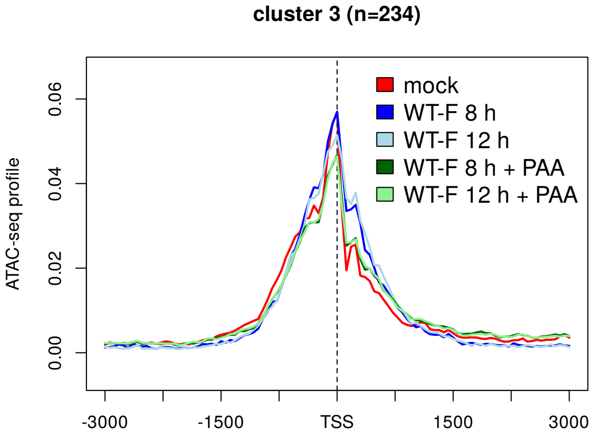


**(m)** pattern I + II

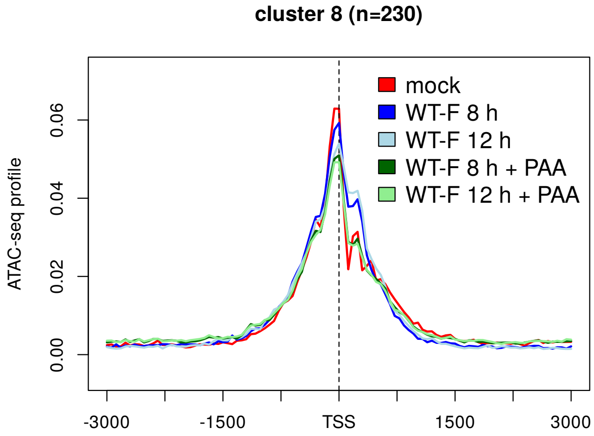


**(n)** pattern I + II

**Sup. Fig 24** Metagene plots showing ATAC-seq profiles in mock and WT-F infection at 8 and 12 h p.i. ± PAA for all clusters (indicated on top of subfigures) ordered according to patterns (indicated below subfigures). See Materials and Methods for a detailed description of metagene plots.

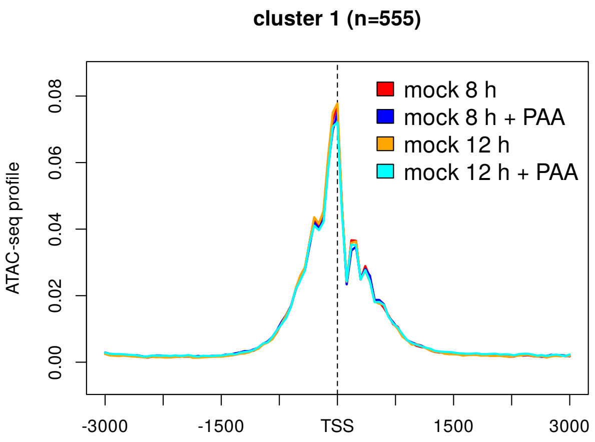


(a) pattern I

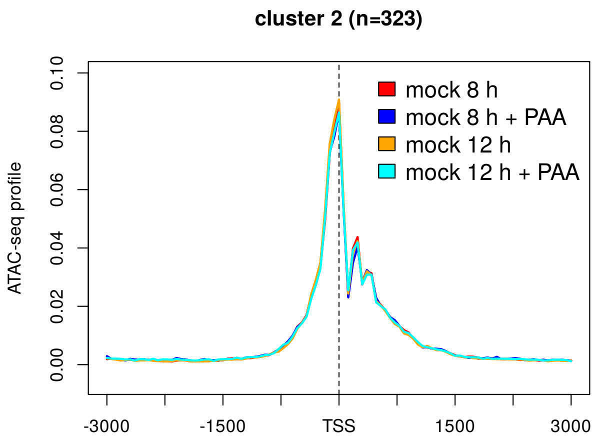


(b) pattern I

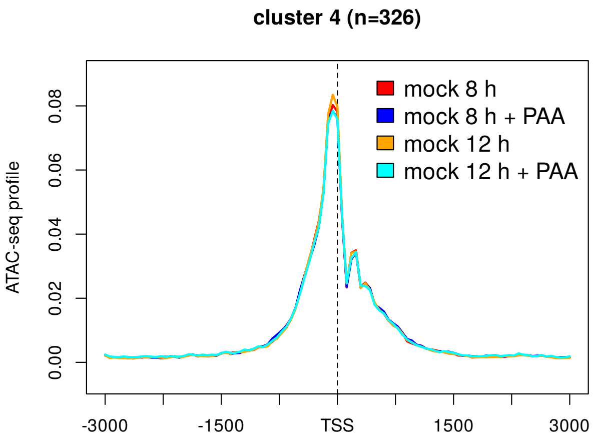


(c) pattern I

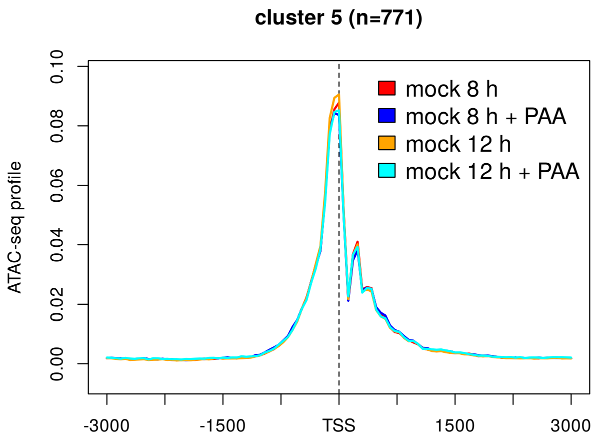


(d) pattern I

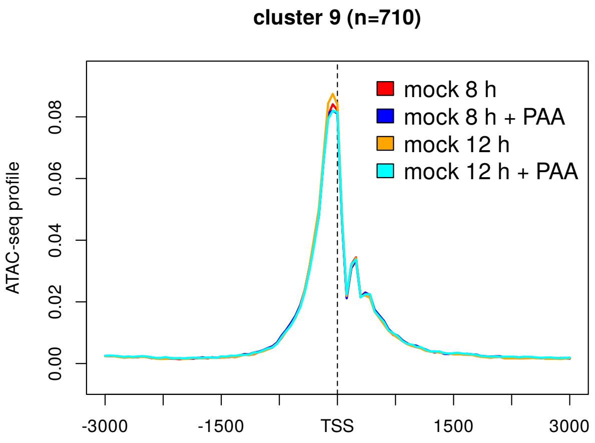


(e) pattern I

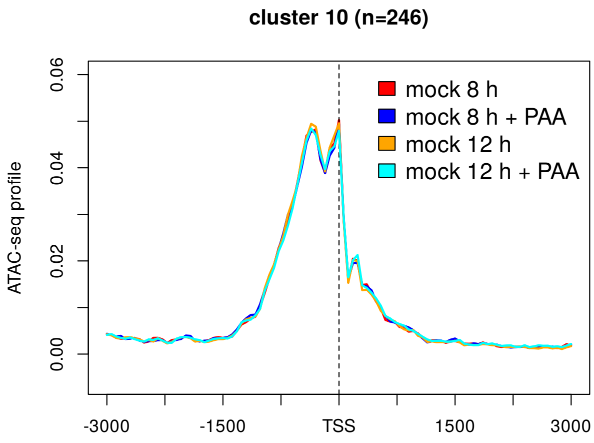


(f) pattern I

(Continued on next page)

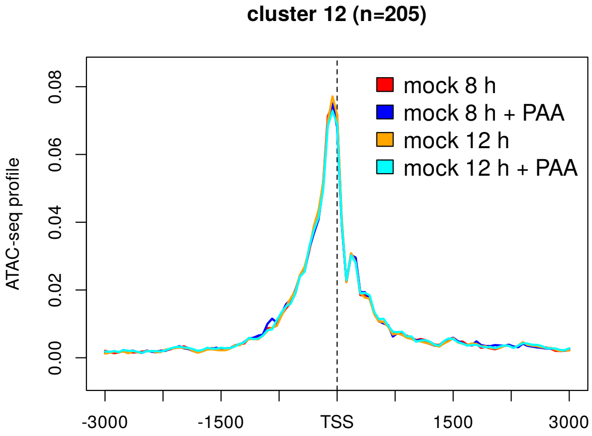


(g) pattern I

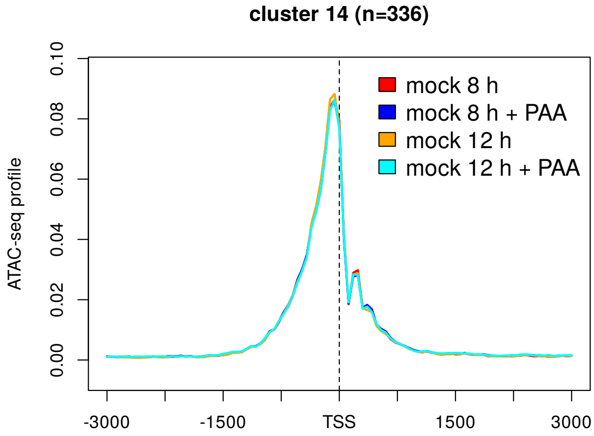


(h) pattern I

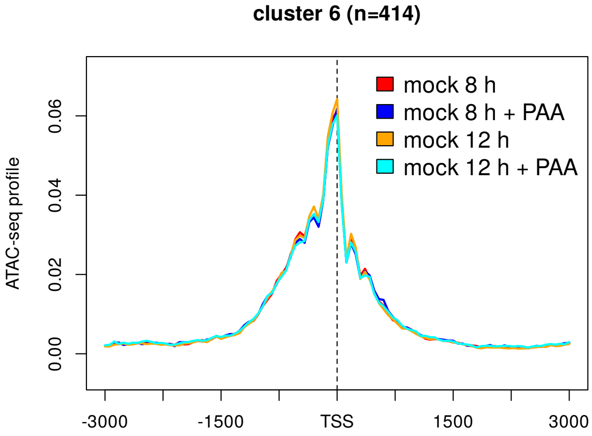


(i) pattern II

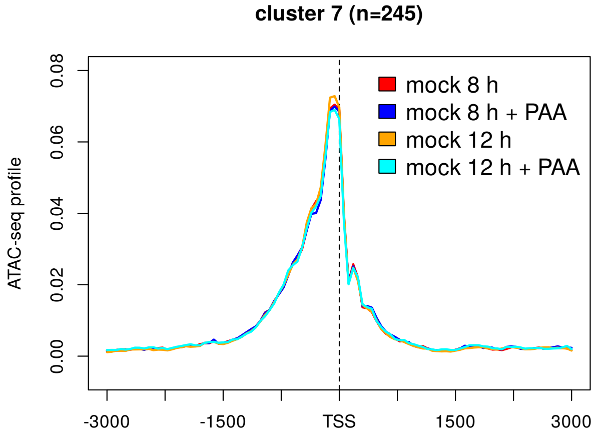


(j) pattern II

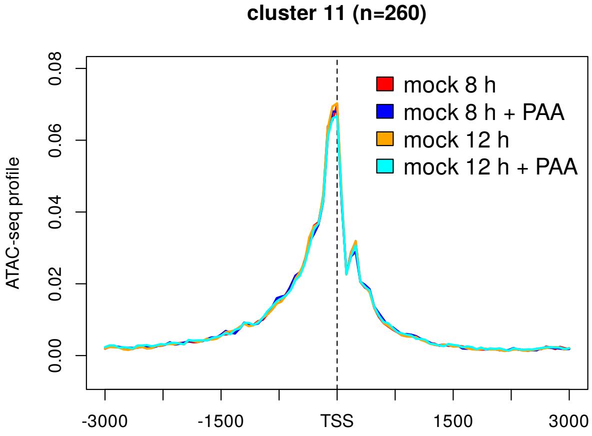


(k) pattern II

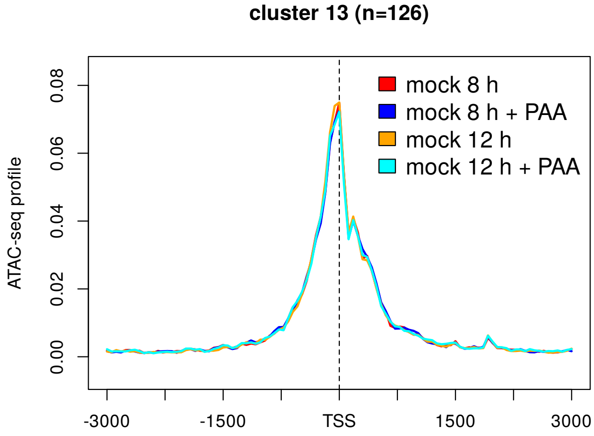


(l) pattern III

**(Continued on next page)**

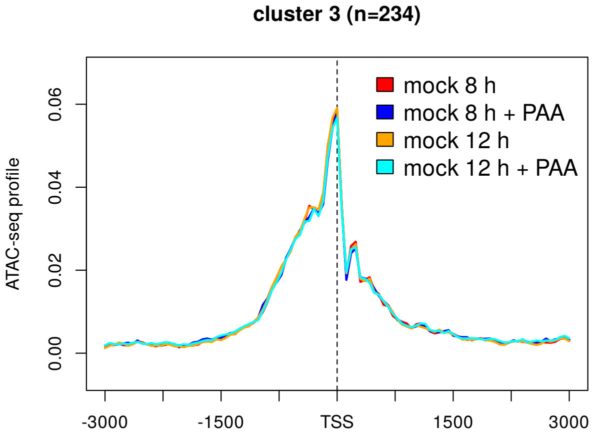


**(m)** pattern I + II

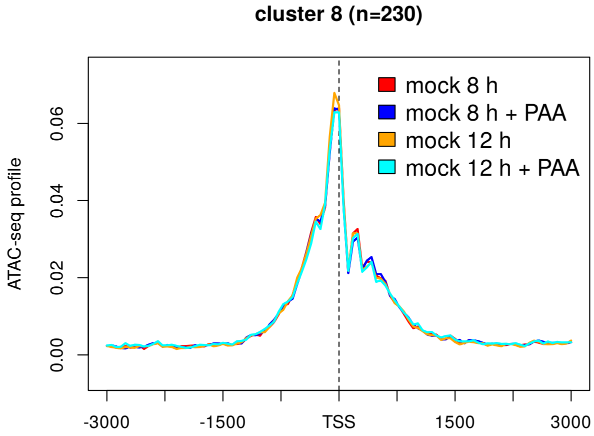


**(n)** pattern I + II

**Sup. Fig 25** Metagene plots showing ATAC-seq profiles in mock infection at 8 and 12 h p.i. ± PAA for all clusters (indicated on top of subfigures) ordered according to patterns (indicated below subfigures). See Materials and Methods for a detailed description of metagene plots.

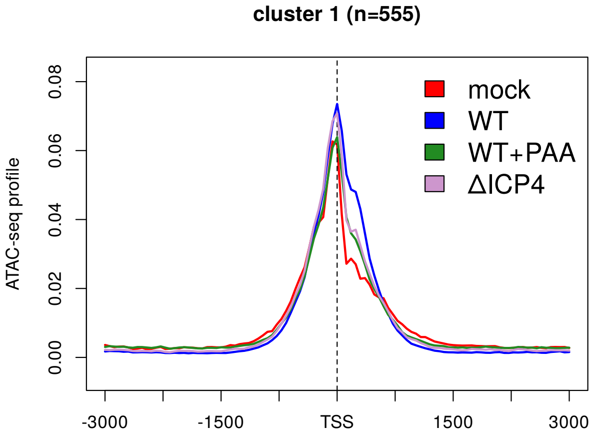


**(a)** pattern I

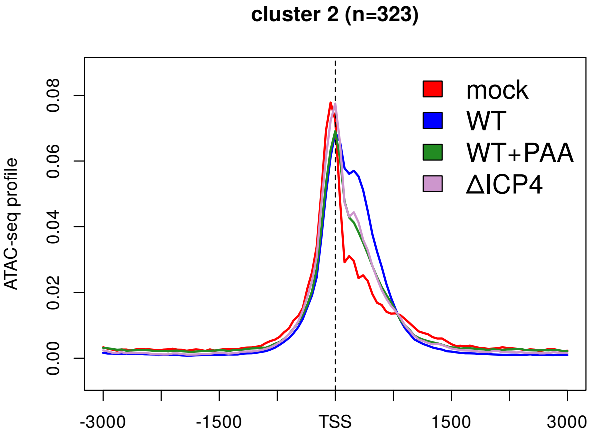


**(b)** pattern I

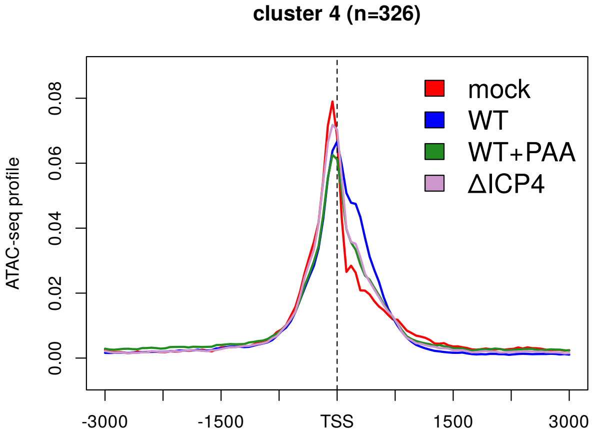


**(c)** pattern I

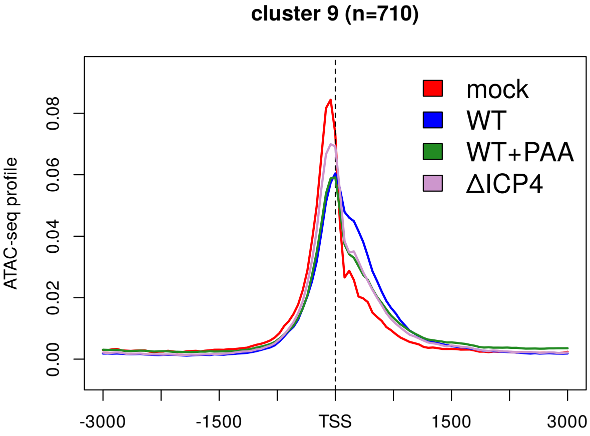


**(d)** pattern I

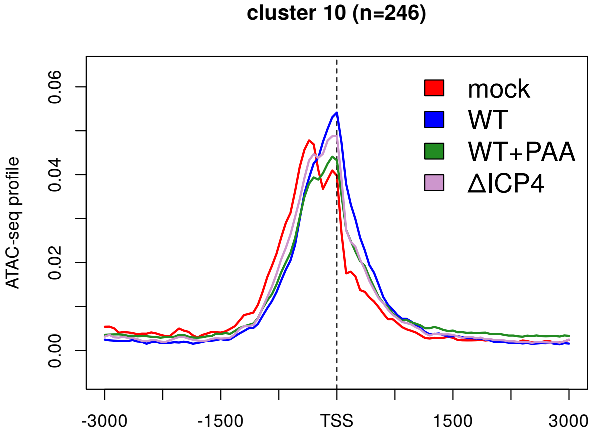


**(e)** pattern I

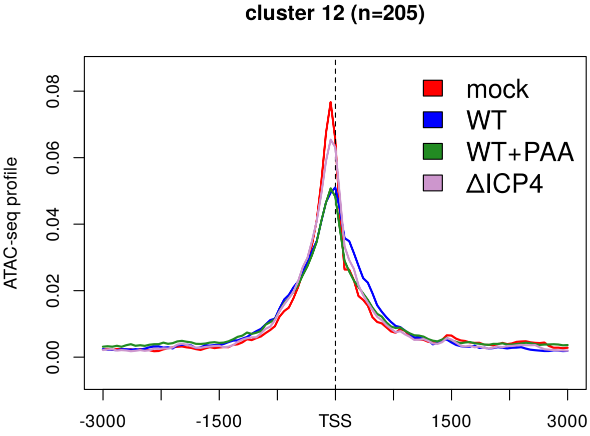


**(f)** pattern I

**(Continued on next page)**

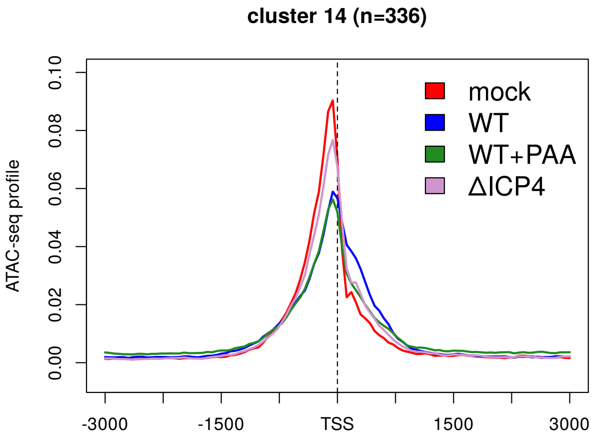


**(g)** pattern I

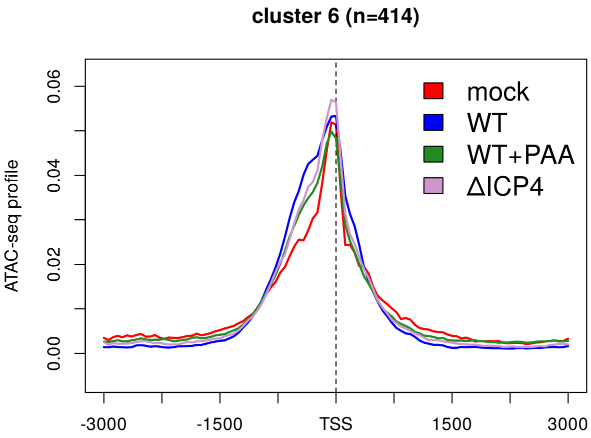


**(h)** pattern II

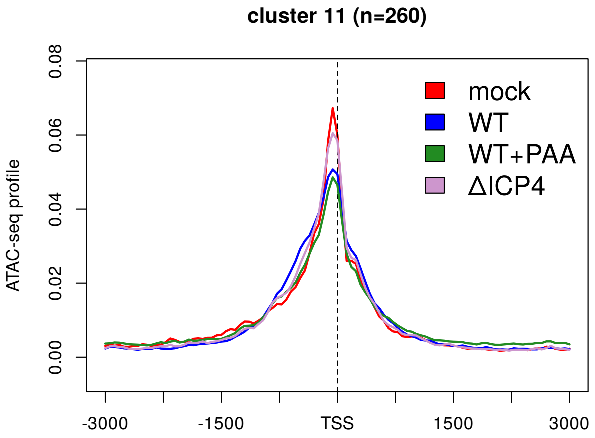


**(i)** pattern II

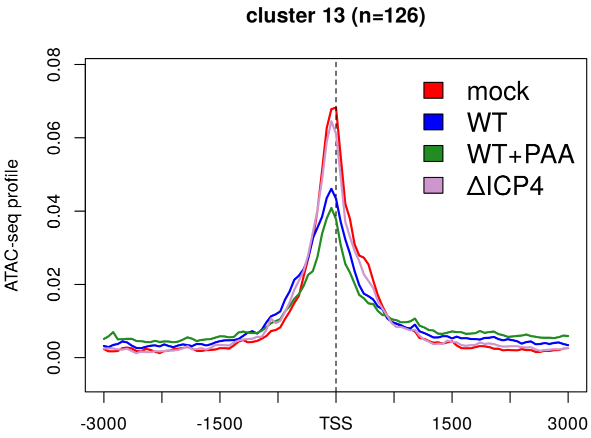


**(j)** pattern III

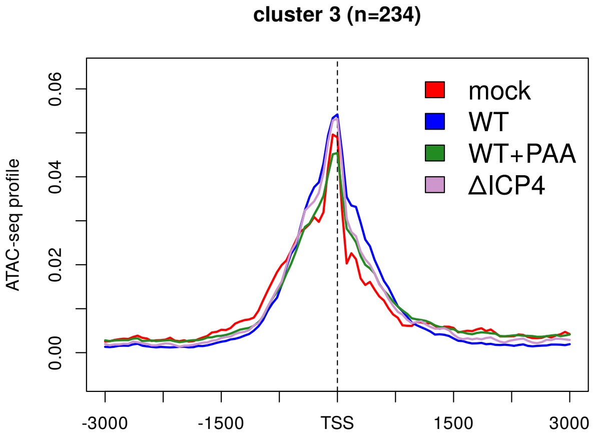


**(k)** pattern I + II

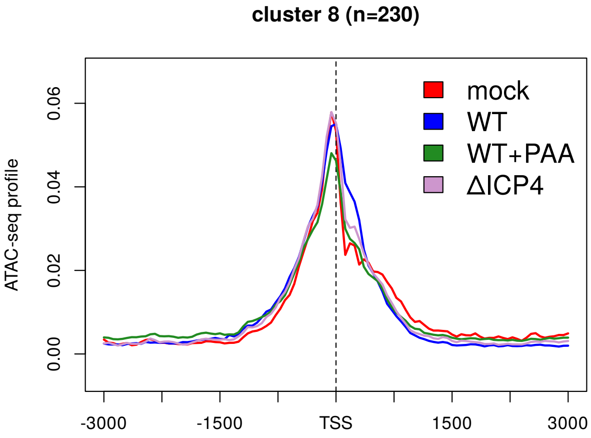


**(l)** pattern I + II

**Sup. Fig 26** Metagene plots of ATAC-seq profiles for mock (red), WT (blue), WT+PAA (green) und ΔICP4 infection (violet) for all clusters (indicated on top of subfigures) ordered according to patterns (indicated below subfigures). See Materials and Methods for a detailed description of metagene plots.

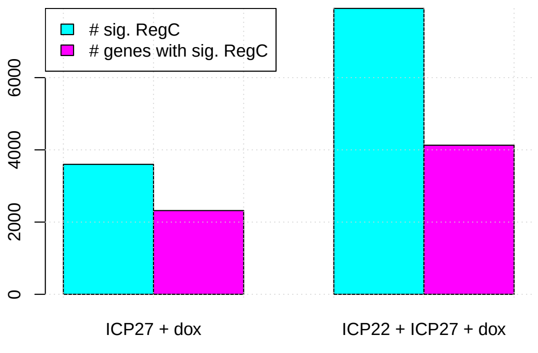


(a)

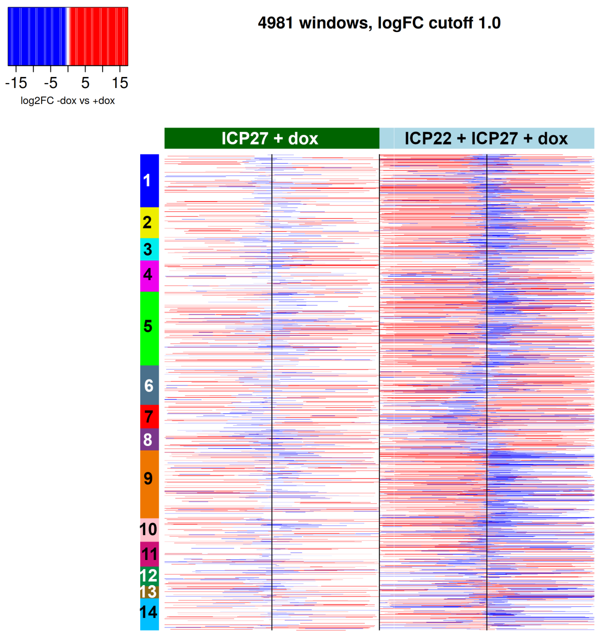


(b)

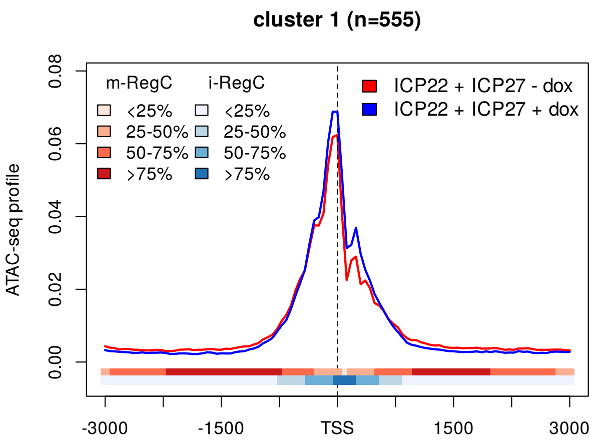


(c) pattern I

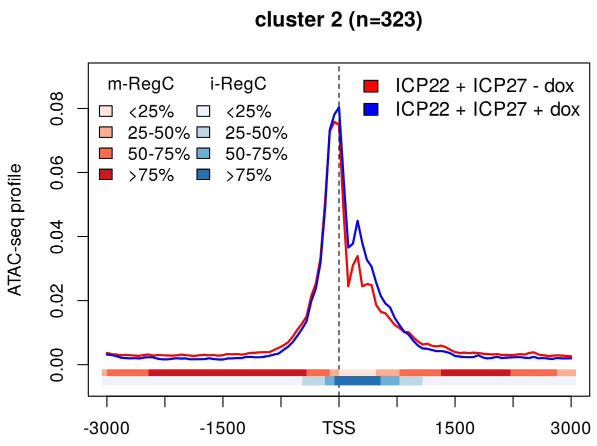


(d) pattern I

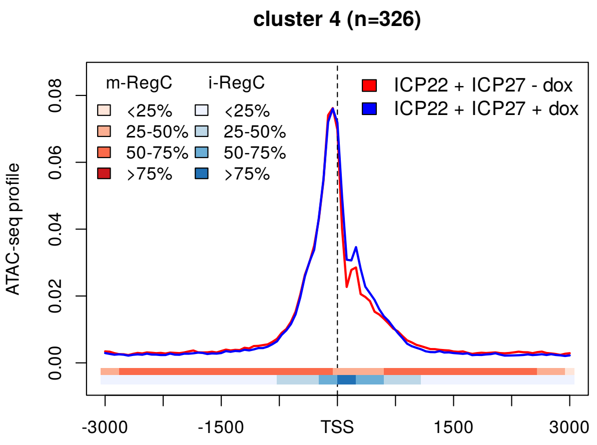


(e) pattern I

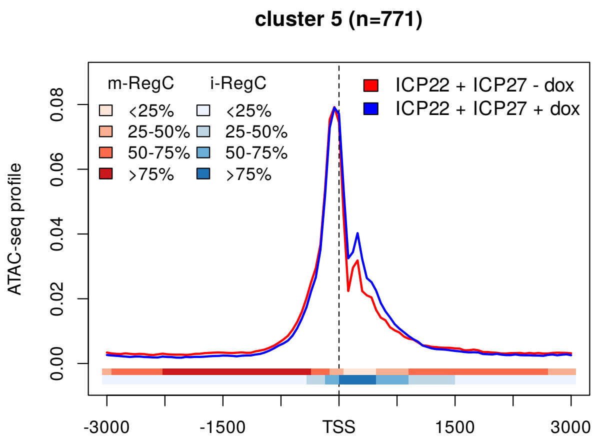


(f) pattern I

(Continued on next page)

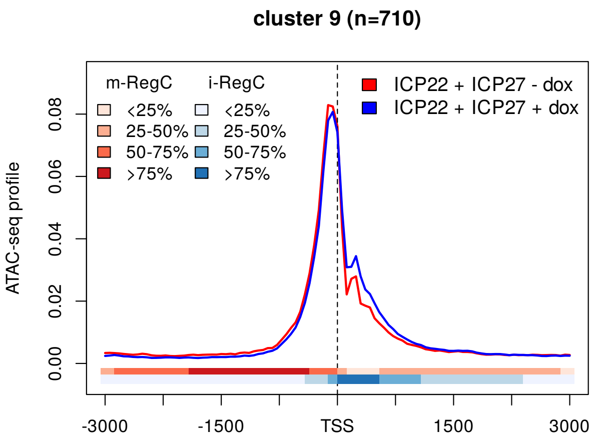


(g) pattern I

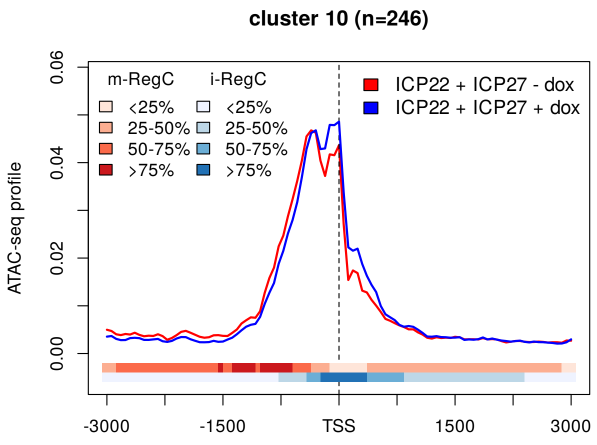


(h) pattern I

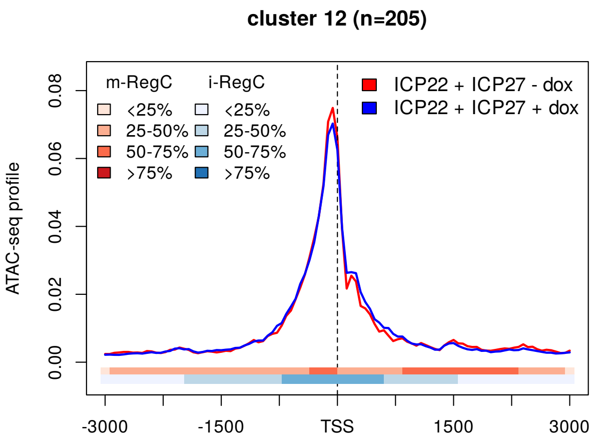


(i) pattern I

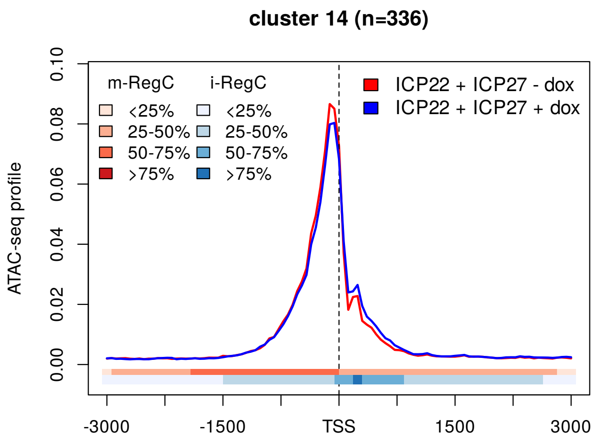


(j) pattern I

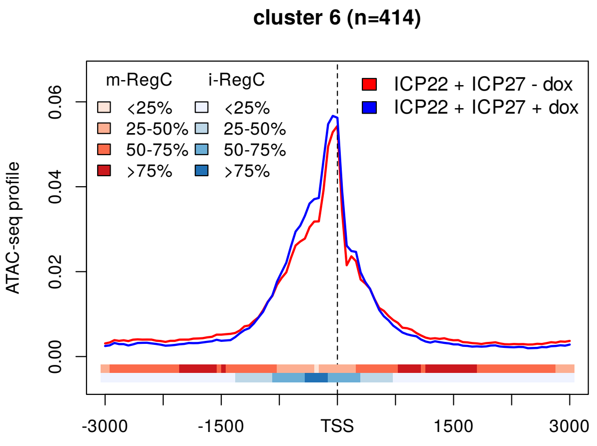


(k) pattern II

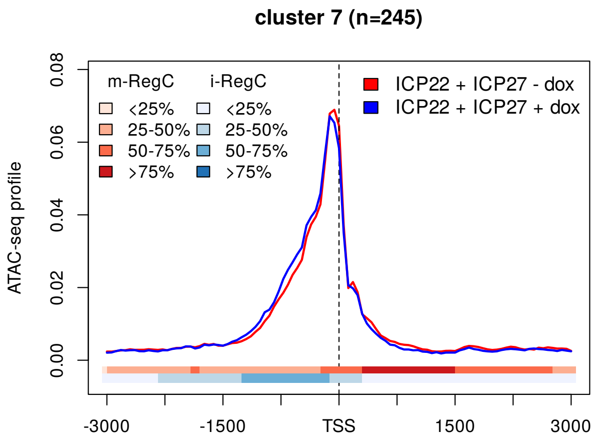


(l) pattern II

**(Continued on next page)**

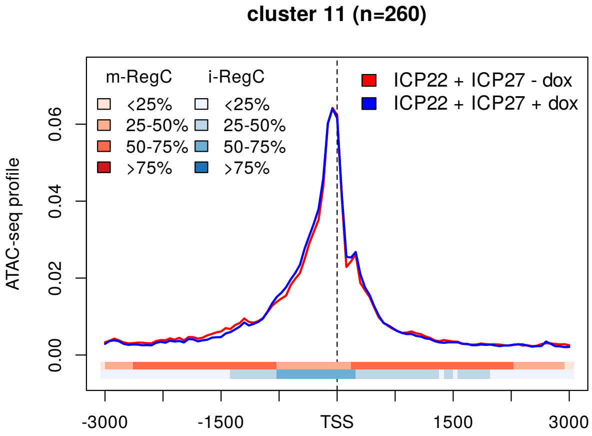


**(m)** pattern II

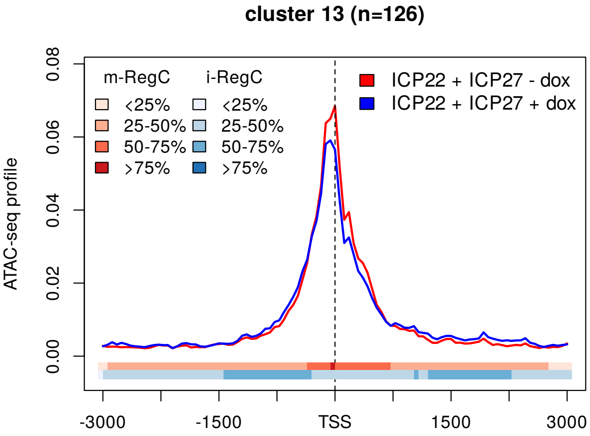


**(n)** pattern III

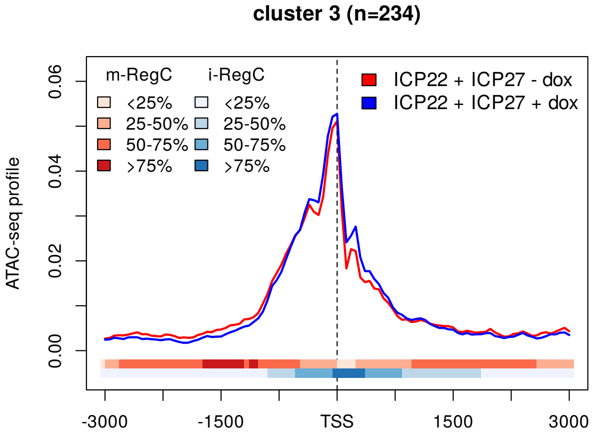


**(o)** pattern I + II

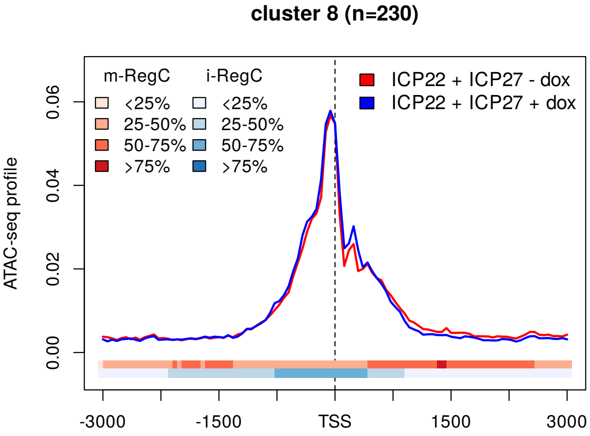


**(p)** pattern I + II

**Sup. Fig 27 (a)** Number of significant RegC and genes with significant RegC identified by RegCFinder for T-HF-ICP27 cells (left) and T-HF-ICP22/ICP27 cells (right) upon dox exposure. **(b)** Heatmap visualizing log2 fold-changes for the differential regions identified for T-HF-ICP27 cells (left half) and T-HF-ICP22/ICP27 cells (right half) upon dox exposure for the promoter windows included in **Fig. 1a**. One row of this heatmap represents results for a particular input window. Black vertical lines in the center of each half of the heatmap indicate the position of the TSS. Regions corresponding to statistically significant differential regions (adj. p. $< 0.01$) are colored according to the log2 fold-change determined by DEXSeq. Here, the color scale is continuous between -1 and 1 and log2 fold-changes $> 1$ are colored the same red and log2 fold-changes $< 1$ the same blue. Promoter windows are ordered as in **Fig. 1a** and clusters from **Fig. 1a** are shown as colored and numbered rectangles on the left. **(c-p)** Metagene plots of ATAC-seq profiles for T-HF-ICP22/ICP27 cells with (blue) and without (red) dox exposure for all clusters (indicated on top of subfigures) ordered according to patterns (indicated below subfigures). See Materials and Methods for a detailed description of metagene plots. The colored bands below the metagene curves indicate the percentage of genes having an m-RegC (red, decreased upon dox exposure) or i-RegC (blue, increased upon dox exposured) or at that position.

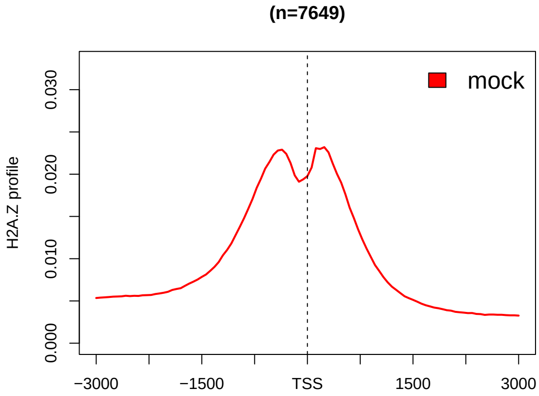


(a)

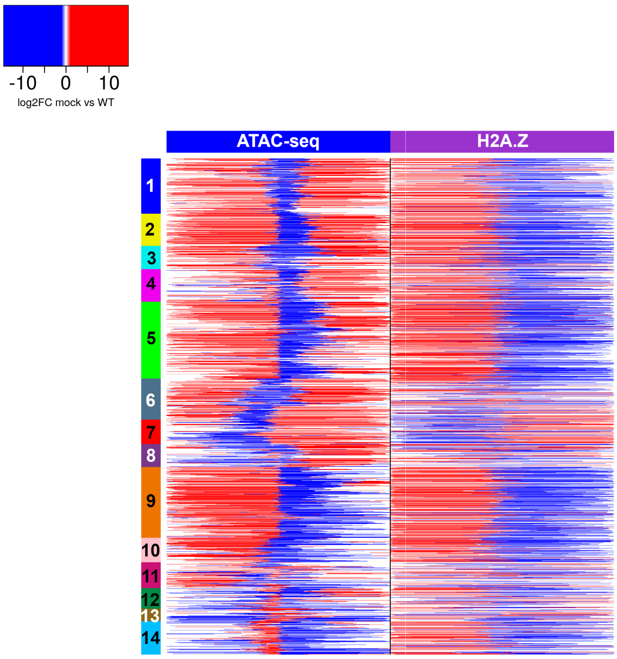


(b)

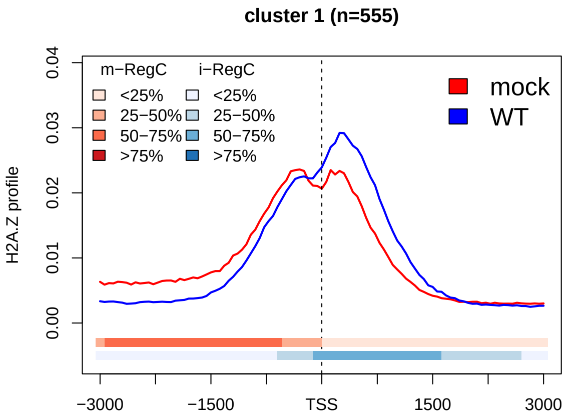


(c) pattern I

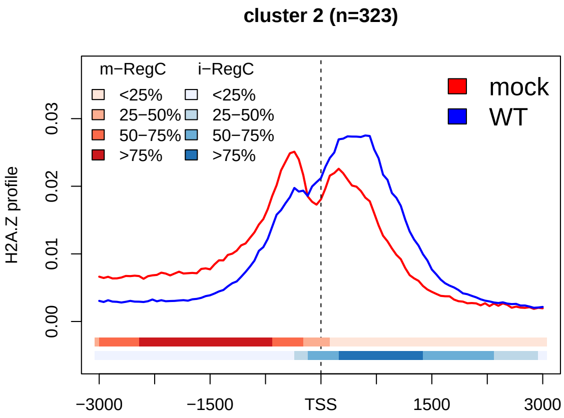


(d) pattern I

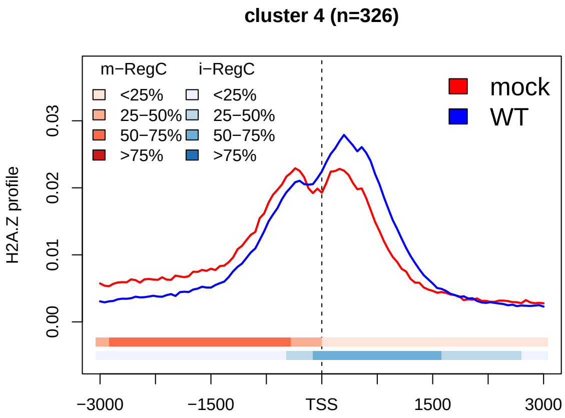


(e) pattern I

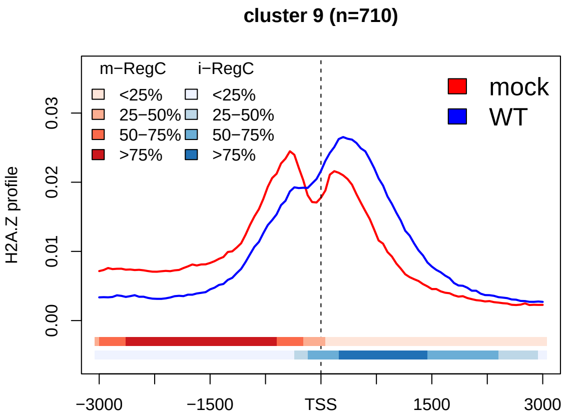


(f) pattern I

(Continued on next page)

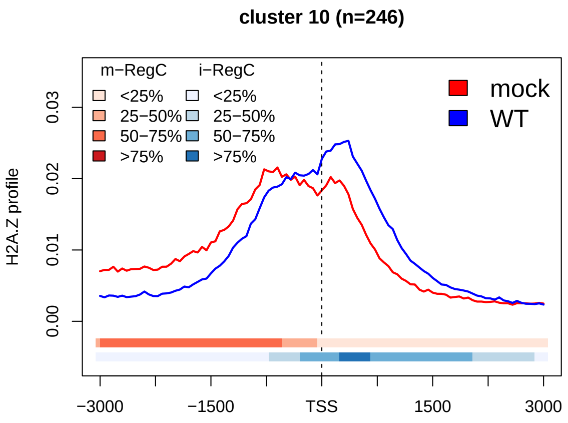


(g) pattern I

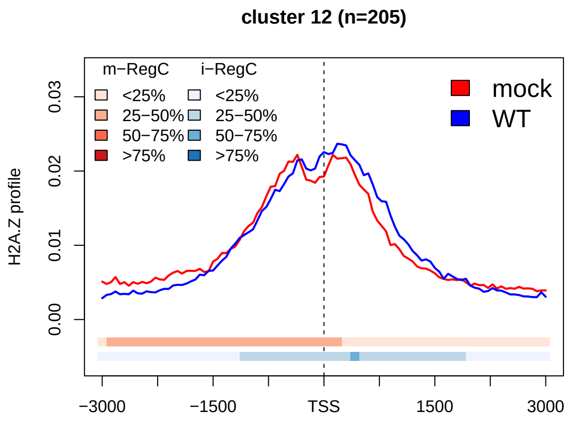


(h) pattern I

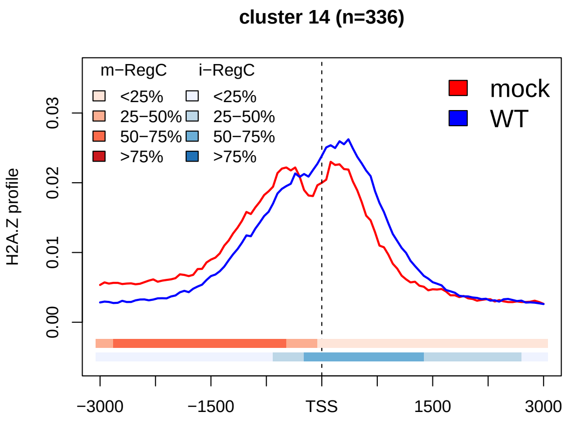


(i) pattern I

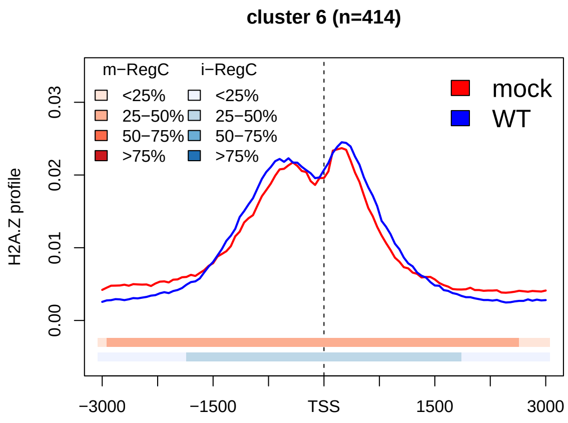


(j) pattern II

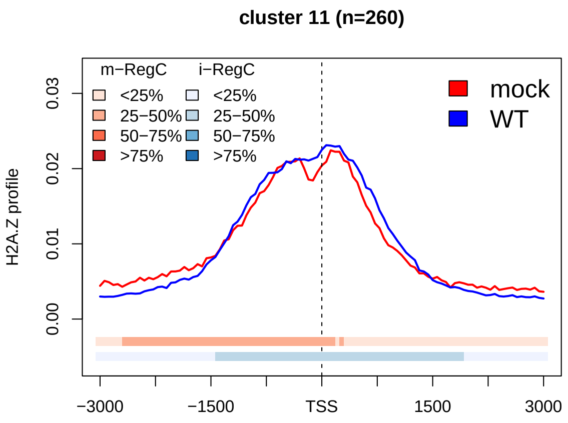


(k) pattern II

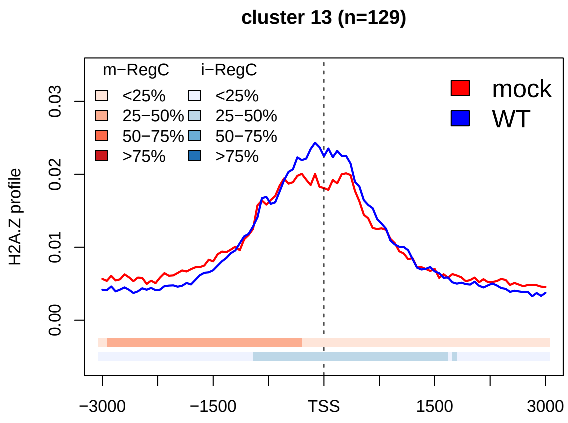


(l) pattern III

**(Continued on next page)**

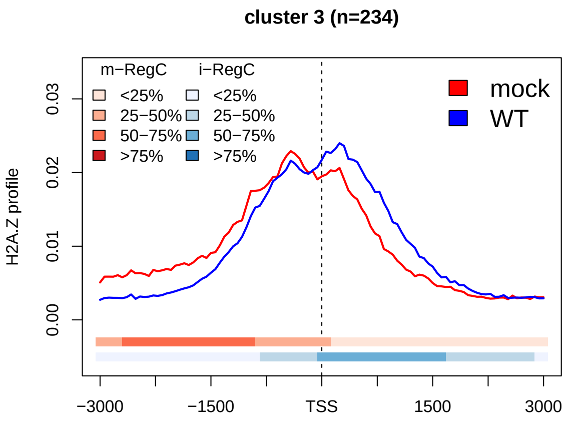


**(m)** pattern I + II

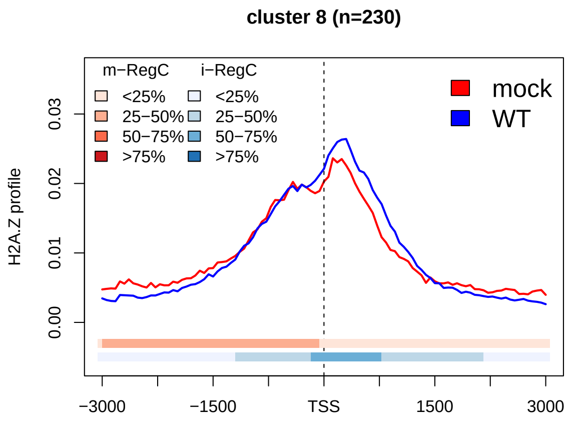


**(n)** pattern I + II

**Sup. Fig 28** (**a**) Metagene plot of H2A.Z profiles for mock infection for all analyzed promoter windows. See Materials and Methods for an explanation of metagene plots. **(b)** Heatmap showing log2 fold-changes for differential regions (m- and i-RegC) identified in the WT vs. mock comparison on the ATAC-seq (left half) and H2A.Z ChIPmentation data (right half) for the promoter windows included in **Fig. 1a**. Statistically significant (adj. p. $< 0.01$) differential regions are colored according to the log2 fold-change determined by DEXSeq. Here, the color scale is continuous between -1 and 1 and log2 fold-changes $> 1$ are colored the same red and log2 fold-changes $< 1$ the same blue. Promoter windows are ordered as in **Fig. 1a** and clusters from **Fig. 1a** are annotated as colored and numbered rectangles on the left. **(c-n)** Metagene plots of H2A.Z profiles in mock (red) and WT (blue) infection for clusters with pattern I (**c-i**, cluster 5 is shown in **Fig. 4d**), pattern II (**j,k**, cluster 7 is shown in **Fig. 4e**), pattern III **(l)** and combined patterns I and II **(m,n)**. See Materials and Methods for an explanation of metagene plots. The colored bands below the metagene curves in each panel indicate the percentage of genes having an m- or i-RegC at that position in the comparison of WT vs. mock in the H2A.Z ChIPmentation data.

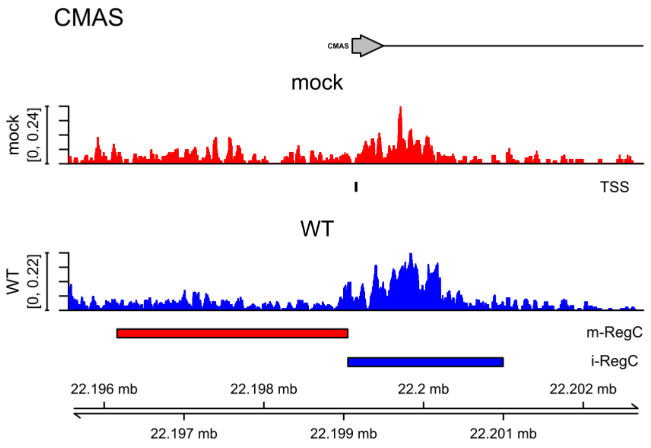


(a) pattern I, cluster 1

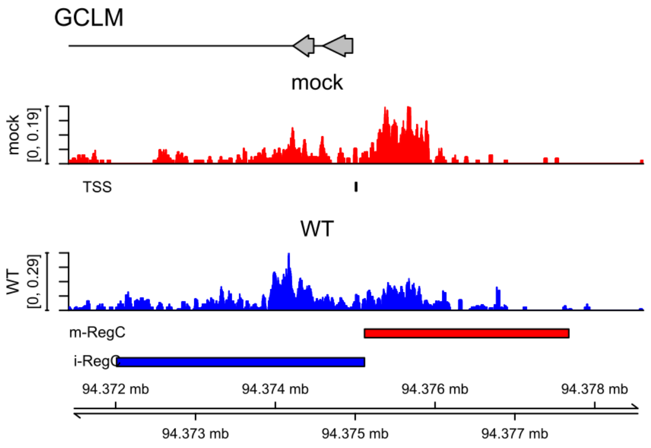


(b) pattern I, cluster 2

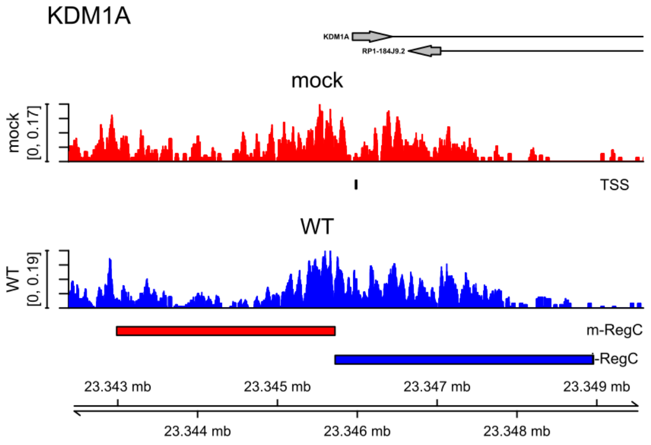


(c) pattern I, cluster 4

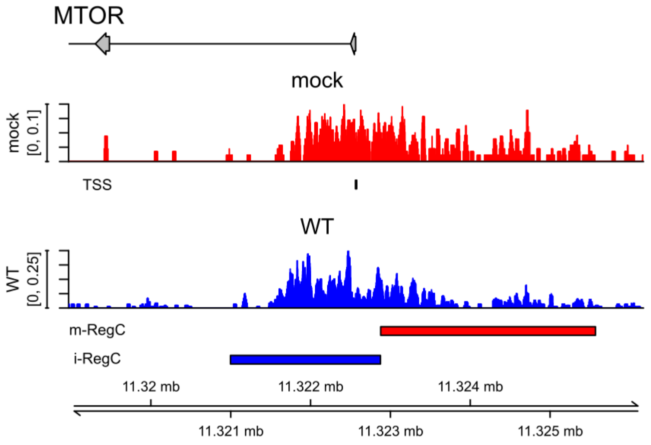


(d) pattern I, cluster 9

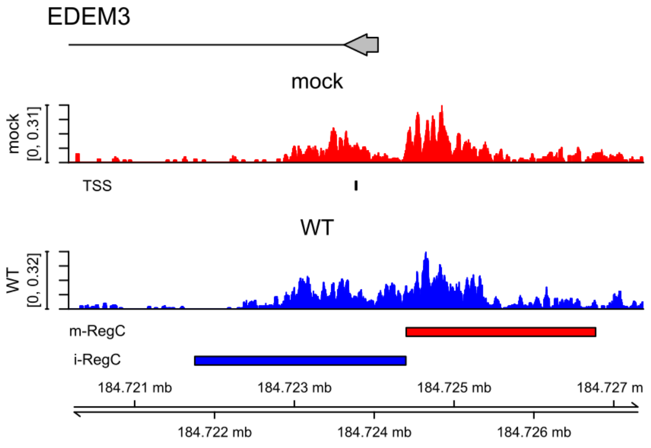


(e) pattern I, cluster 10

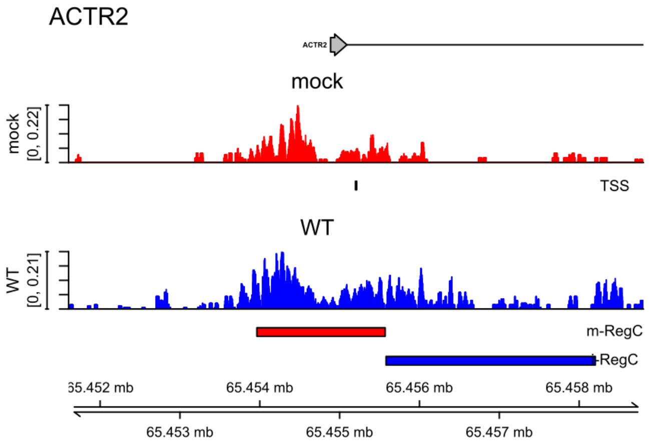


(f) pattern I, cluster 12

**(Continued on next page)**

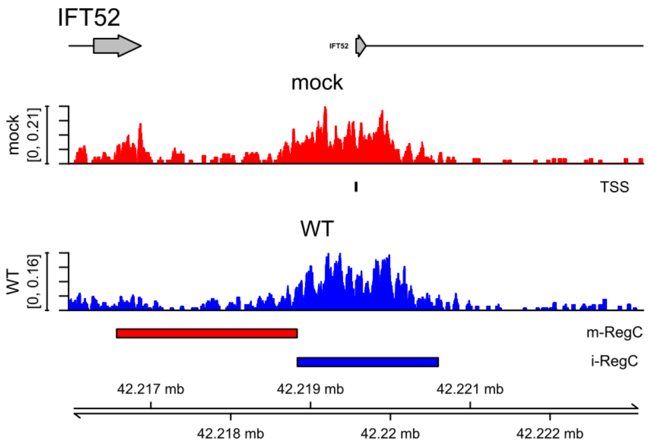


**(g)** pattern I, cluster 14

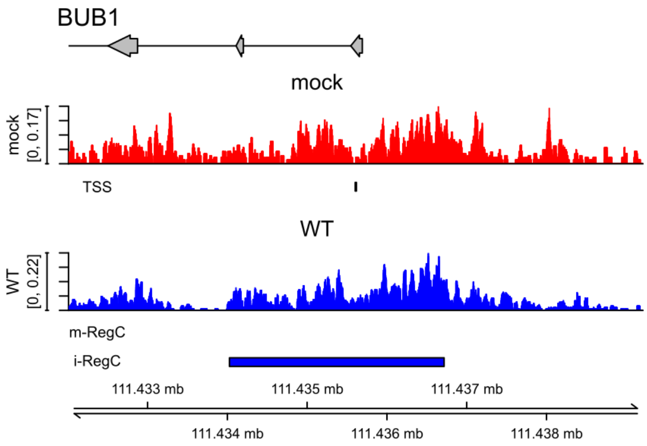


**(h)** pattern II, cluster 6

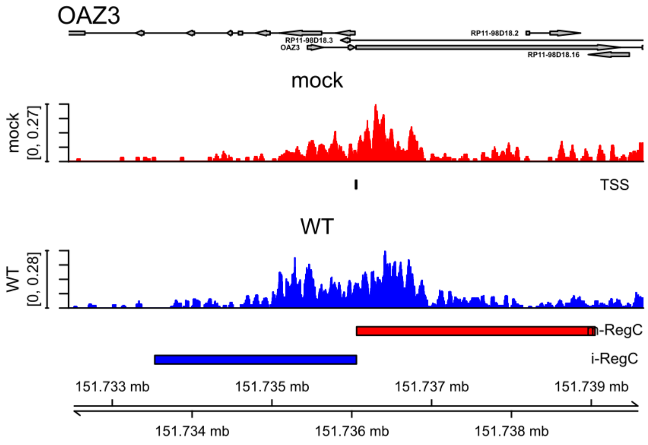


**(i)** pattern II, cluster 11

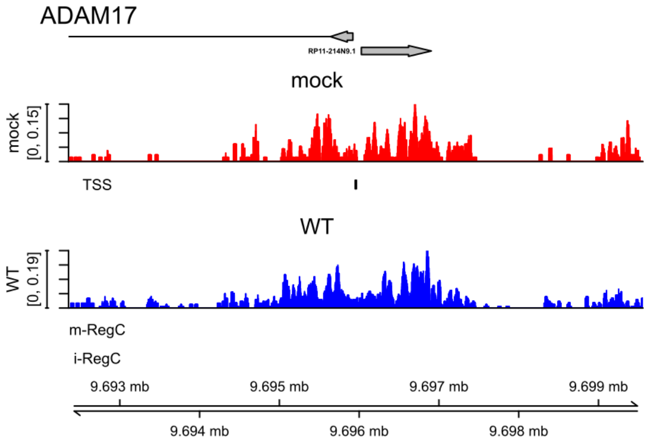


**(j)** pattern III, cluster 13

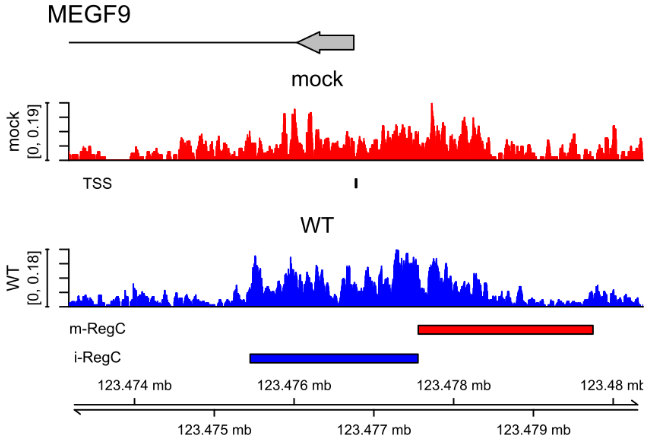


**(k)** pattern I + II, cluster 3

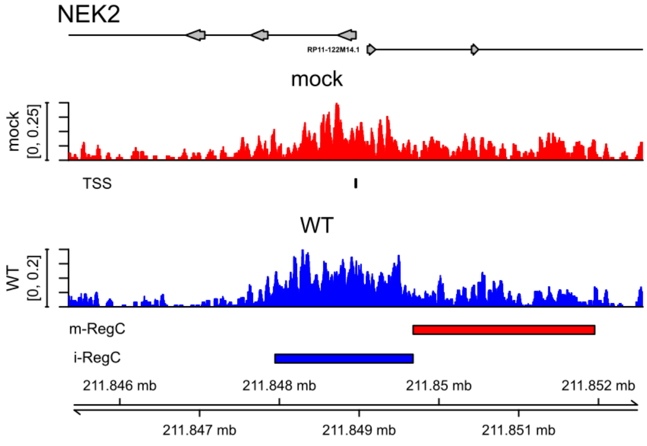


**(l)** pattern I + II, cluster 8

**(Continued on next page)**

**Sup. Fig 29** Read coverage in a ± 3.6 kb window around the TSS in H2A.Z ChIPmentation data for mock (red) and WT (blue) infection for example genes with pattern I **(a-g)**, pattern II **(h,i)**, pattern III **(j)** and combined patterns I and II **(k,l)**. Read coverage was normalized to total number of mapped reads for each sample and averaged between replicates. The TSS used in the analysis is indicated by a short vertical line below the read coverage track for mock infection. Gene annotation is indicated at the top. Boxes represent exons, lines represent introns, and direction of transcription is indicated by arrowheads. The name of the gene whose promoter window was analyzed is indicated in larger font on the top left and – if not clear from the context – beside the gene annotation. Names for other genes overlapping the input window are also indicated if necessary. Below each read coverage track m- (red bars) and i-RegCs (blue bars) are indicated for the comparison of WT vs. mock om the H2A.Z ChIPmentation data. Corresponding read coverage plots in ATAC-seq data for mock, WT, ΔICP0, ΔICP22, ΔICP27, Δ*vhs* and WT+PAA infection are shown in **Sup. Fig. 4**.

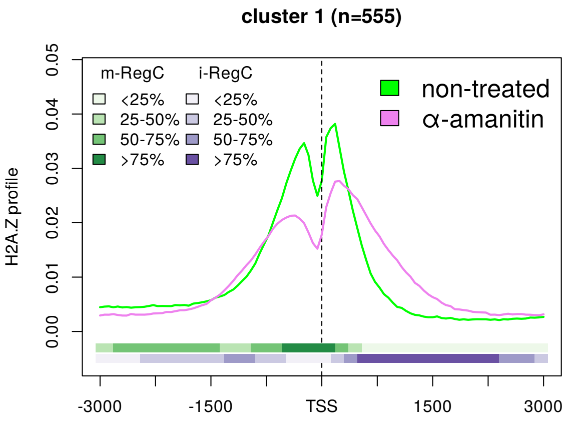


(a) pattern I

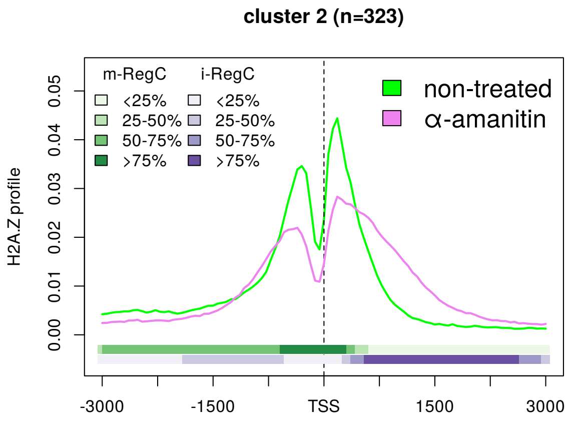


(b) pattern I

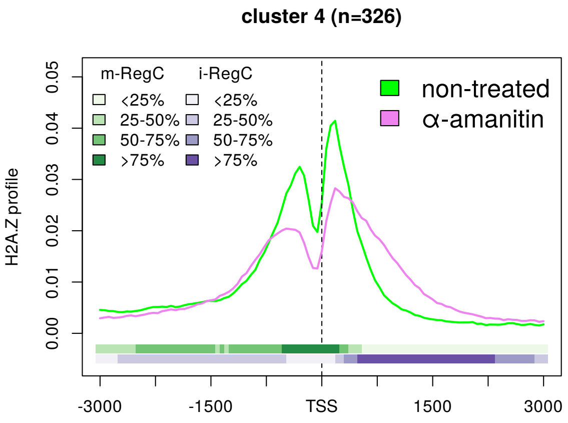


(c) pattern I

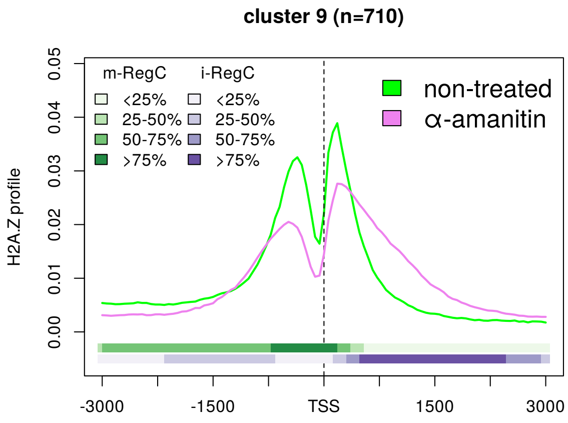


(d) pattern I

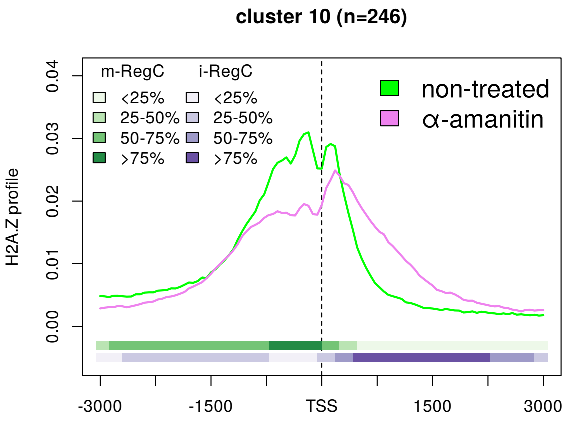


(e) pattern I

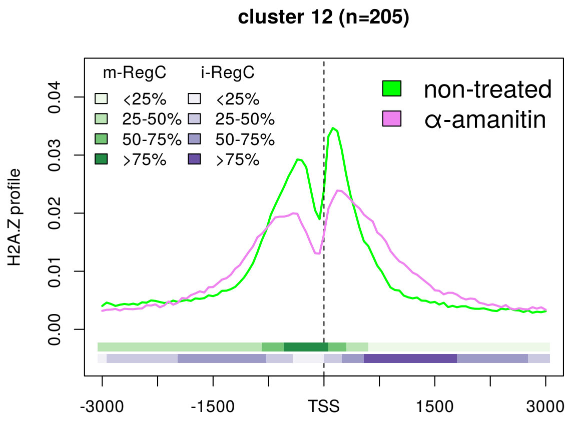


(f) pattern I

**(Continued on next page)**

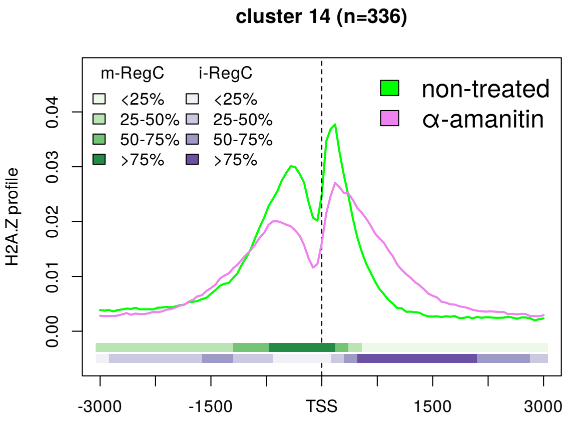


**(g)** pattern I

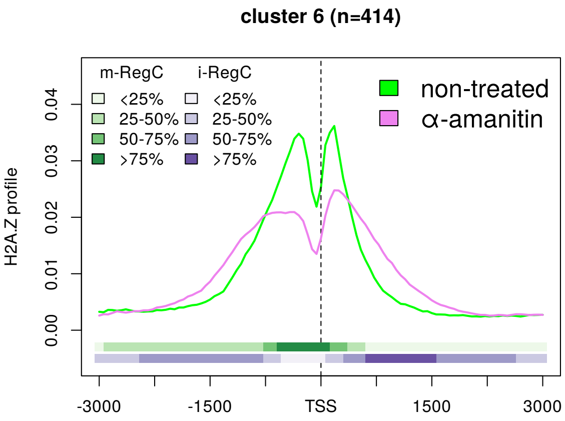


**(h)** pattern II

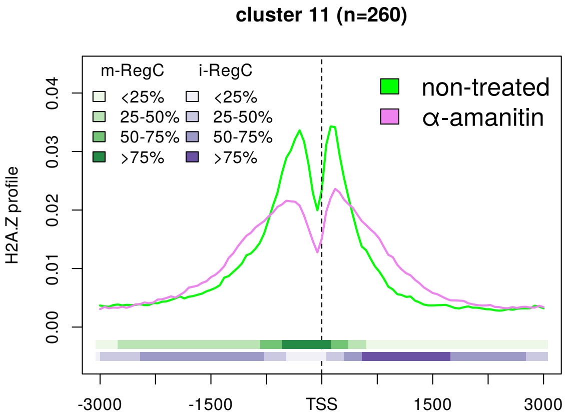


**(i)** pattern II

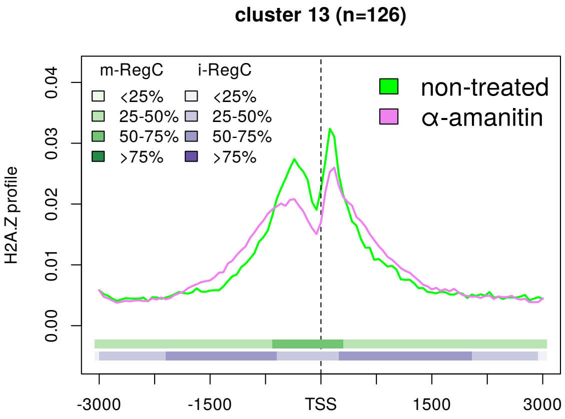


**(j)** pattern III

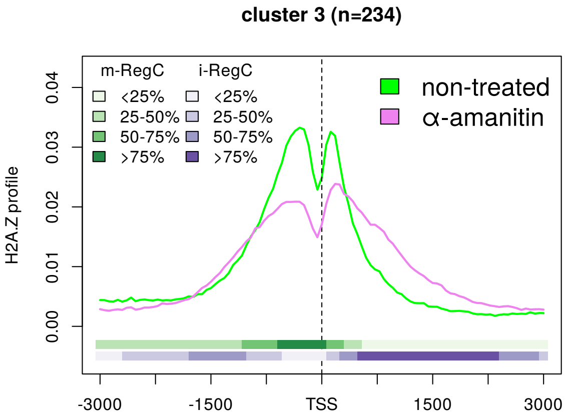


**(k)** pattern I + II

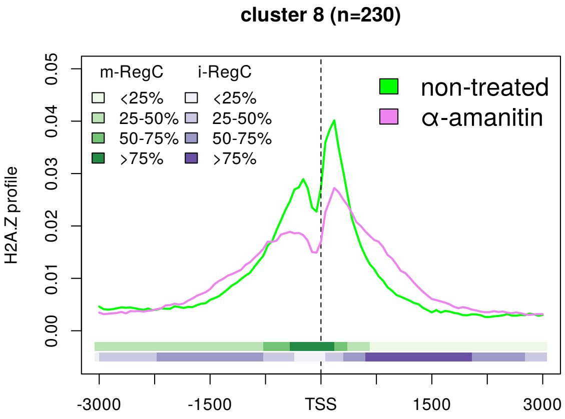


**(l)** pattern I + II

**Sup. Fig 30** Metagene plots of H2A.Z profiles for untreated (green) and $\alpha$-amanitin-treated (violet) HCT116 cells for clusters with pattern I (**c-g**, cluster 5 is shown in **Fig. 4f**), pattern II (**h,i**, cluster 7 is shown in **Fig. 4g**), pattern III **(j)** and combined patterns I and II **(k,l)**. See Materials and Methods for an explanation of metagene plots. The colored bands below the metagene curves in each subfigure indicate the percentage of genes having an m- or i-RegC at that position in the comparison of $\alpha$-amanitin-treatment vs. no treatment in the H2A.Z ChIP-seq data. Here, m-RegC are differential regions with relative read density higher in untreated cells and i-RegC differential regions with relative read density higher upon $\alpha$-amanitin-treatment.

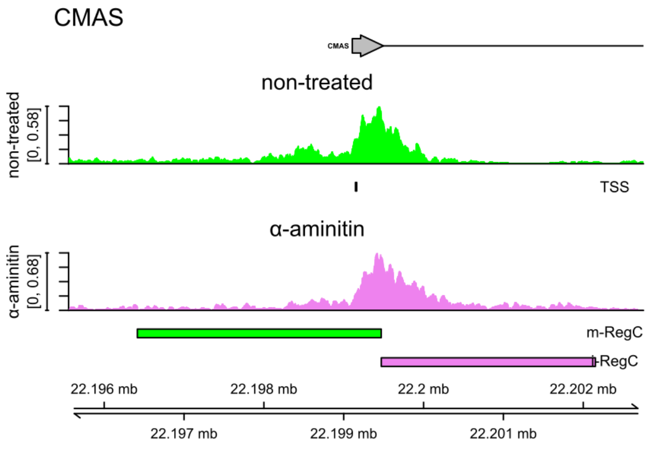


(a) pattern I, cluster 1

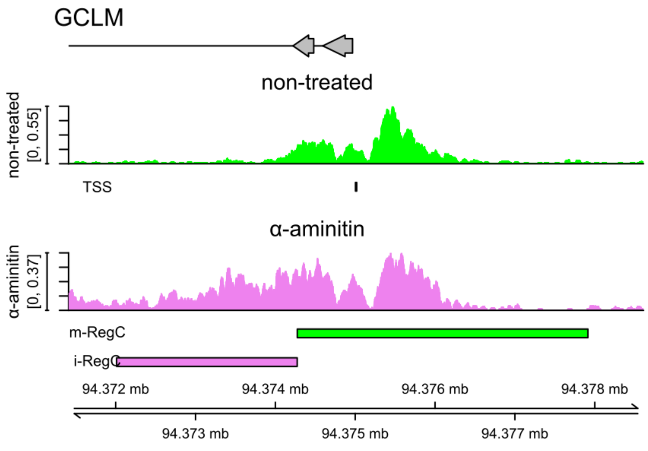


(b) pattern I, cluster 2

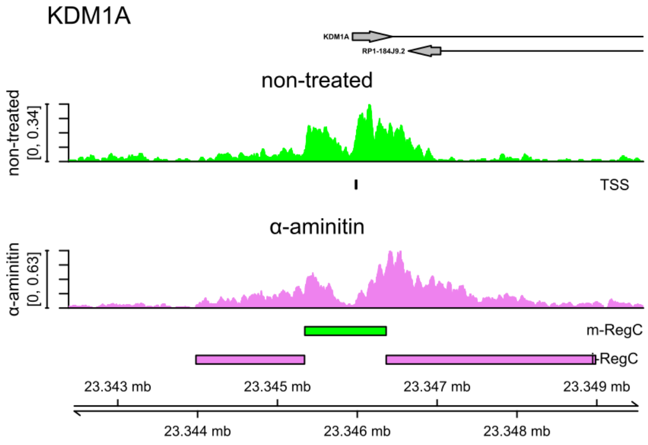


(c) pattern I, cluster 4

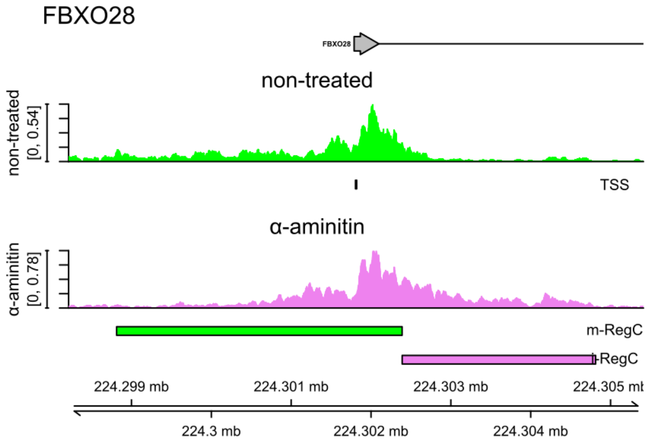


(d) pattern I, cluster 5

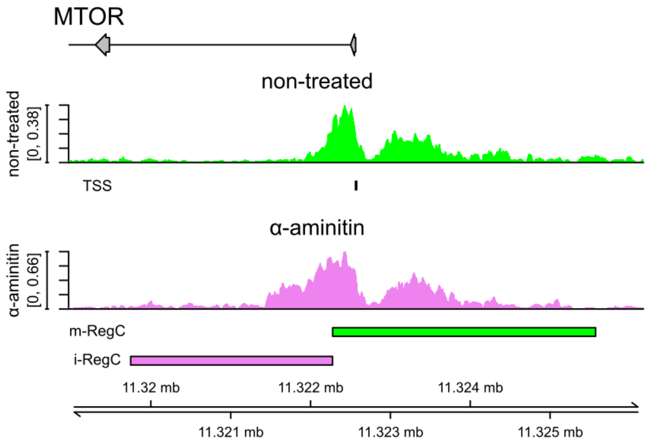


(e) pattern I, cluster 9

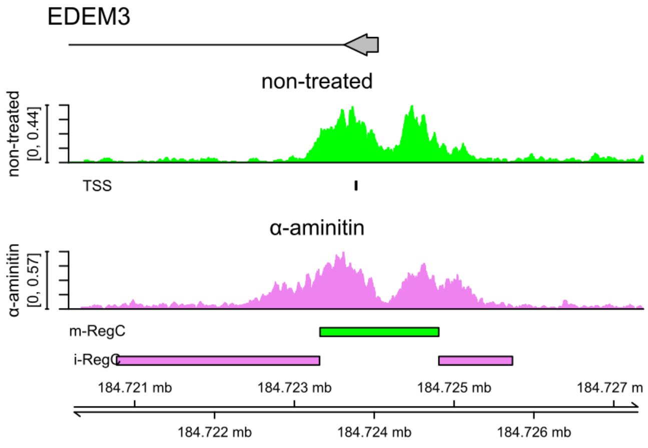


(f) pattern I, cluster 10

**(Continued on next page)**

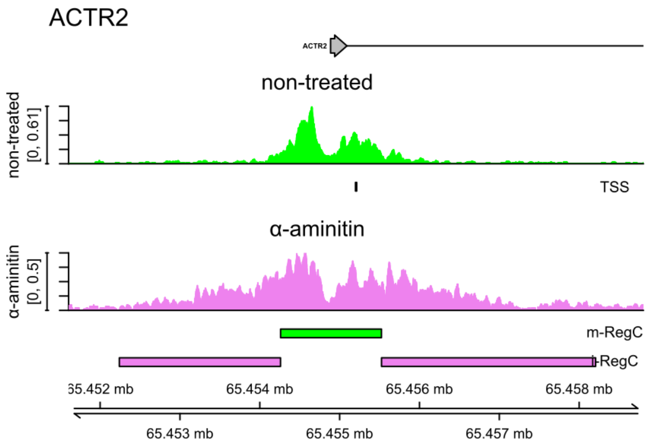


(g) pattern I, cluster 12

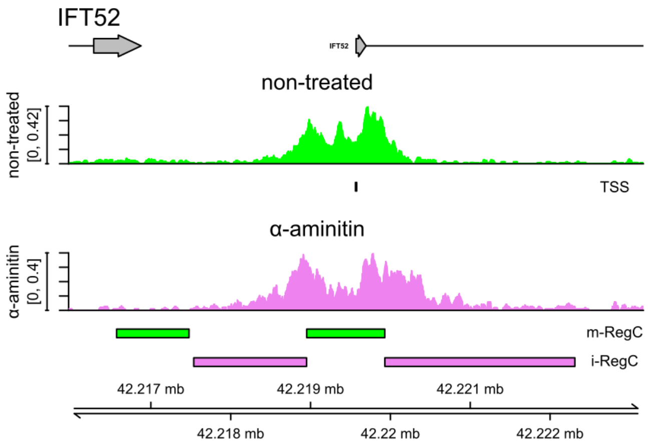


(h) pattern I, cluster 14

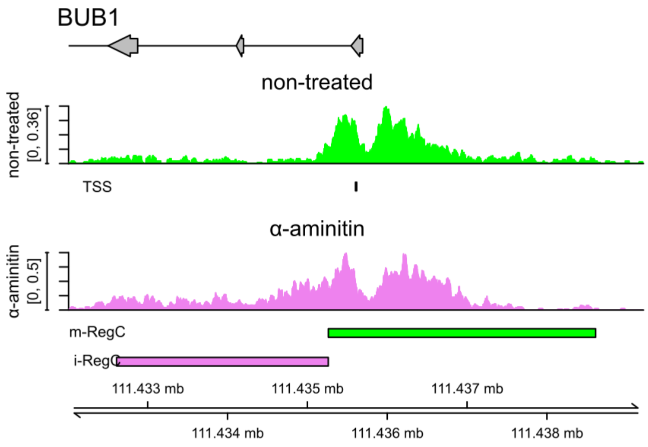


(i) pattern II, cluster 6

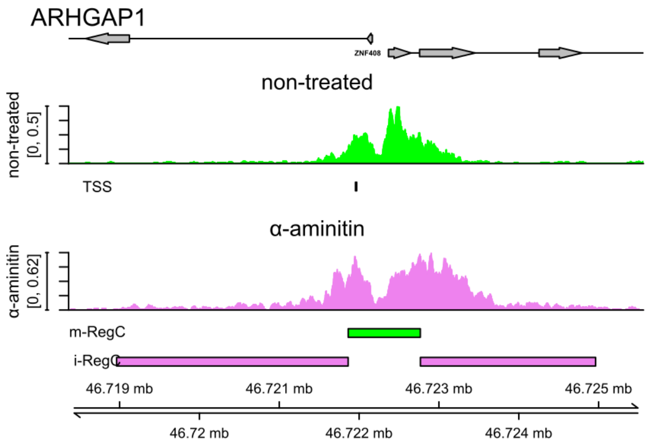


(j) pattern II, cluster 7

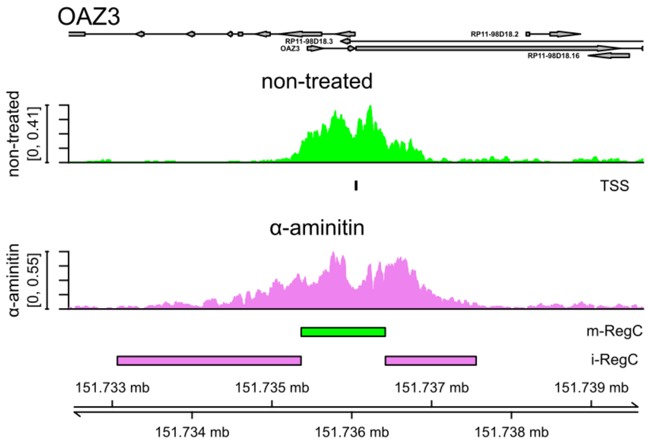


(k) pattern II, cluster 11

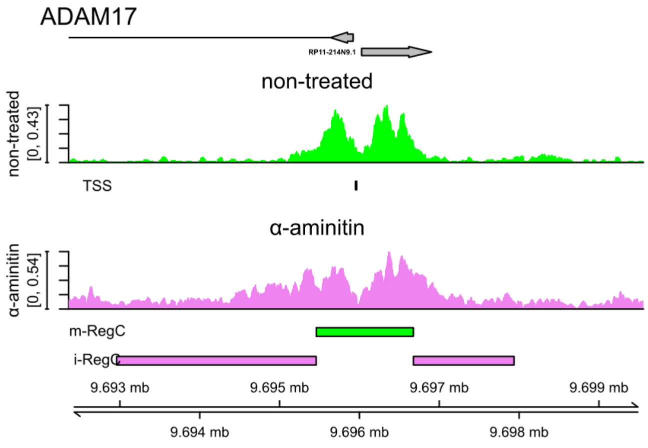


(l) pattern III, cluster 13

**(Continued on next page)**

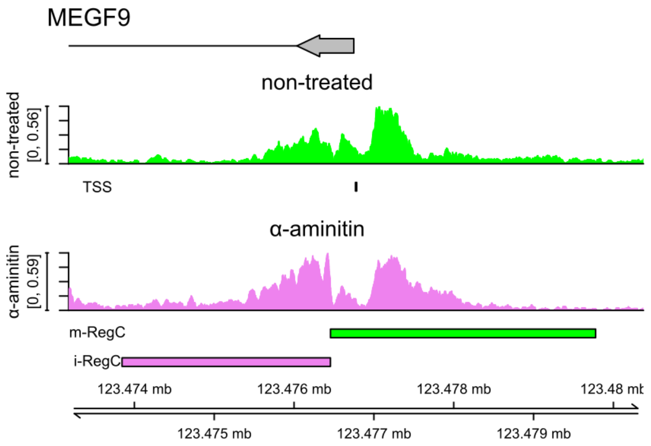


**(m)** pattern I + II, cluster 3

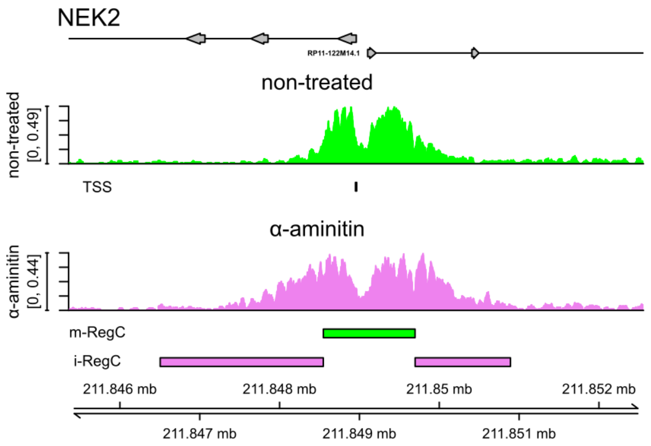


**(n)** pattern I + II, cluster 8

**Sup. Fig 31** Read coverage in a ± 3.6 kb window around the TSS in the H2A.Z ChIP-seq data for untreated (green) and $\alpha$-amanitin-treated (violet) HCT116 cells for example genes for all patterns and clusters. Pattern and cluster are indicated below each subfigure. Read coverage was normalized to total number of mapped reads for each sample and averaged between replicates. The TSS used in the analysis is indicated by a short vertical line below the read coverage track for untreated cells. Gene annotation is indicated at the top. Boxes represent exons, lines represent introns, and direction of transcription is indicated by arrowheads. The name of the gene whose promoter window was analyzed is indicated in larger font on the top left and – if not clear from the context – beside the gene annotation. Names for other genes overlapping the input window are also indicated when necessary. Below each read coverage track m-RegCs (i.e. differential regions with relative read coverage higher in untreated cells) and i-RegC (i.e. differential regions with relative read coverage higher upon $\alpha$-amanitin-treatment) are indicated.
